# EPIGENETIC MODULATION VIA THE C-TERMINAL TAIL OF H2A.Z

**DOI:** 10.1101/2021.02.22.432230

**Authors:** László Imre, Péter Nánási, Ibtissem Benhamza, Kata Nóra Enyedi, Gábor Mocsár, Rosevalentine Bosire, Éva Hegedüs, Erfaneh Firouzi Niaki, Ágota Csóti, Zsuzsanna Darula, Éva Csősz, Szilárd Póliska, Beáta Scholtz, Gábor Mező, Zsolt Bacsó, H. T. Marc Timmers, Masayuki Kusakabe, Margit Balázs, György Vámosi, Juan Ausio, Peter Cheung, Katalin Tóth, David Tremethick, Masahiko Harata, Gábor Szabó

## Abstract

H2A.Z-nucleosomes are present in both euchromatin and heterochromatin and it has proven difficult to interpret their disparate roles in the context of their stability features. Using an *in situ* assay of nucleosome stability and DT40 cells expressing engineered forms of the histone variant we show that native H2A.Z, but not C-terminally truncated H2A.Z (H2A.ZΔC), is released from nucleosomes of peripheral heterochromatin at unusually high salt concentrations. H2A.Z and H3K9me3 landscapes are reorganized in H2A.ZΔC-nuclei and overall sensitivity of chromatin to nucleases is increased. These tail-dependent differences are recapitulated upon treatment of HeLa nuclei with the H2A.Z-tail-peptide (C9), with MNase sensitivity being increased at specific regions including promoters. Introduced into live cells C9 elicits down-regulation of ∼560 genes with nonrandom chromosomal band-localization and pathway-spectrum. Thus, tail-dependent heterogeneity of H2A.Z-nucleosomes is revealed at all organization levels of chromatin and epigenetic modulation can be achieved by targeting molecular interactions involving its C-terminal tail.

## INTRODUCTION

The variant histone H2A.Z is involved in the regulation of diverse and basic cell functions, with activating as well as repressive effects that have proved difficult to reconcile with the structural information available for the nucleosomes containing H2A.Z. It contributes to the regulation of transcriptional initiation and elongation, DNA replication, heterochromatin organization, DNA repair, cell differentiation and cell cycle, and epithelial mesenchymal transition in embryonic development. It is also up-regulated in a variety of different types of cancers, has important roles in intestinal epithelial cell homeostasis, Wnt, Notch, Nanog, Gli1 signaling, and in central nervous system development and function (for reviews see ^1, 2^). H2A.Z exhibits about 60% identity to canonical H2A and constitutes approximately 5% of the total H2A histone pool in vertebrates ^3^.

Since the nucleosomal structure is, in general, repressive for transcription, replication and repair, one strategy of eukaryotic cells to regulate these activities involves de-repression by destabilizing or mobilizing particular nucleosomes. As such, the stability of nucleosomes is of utmost regulatory importance, and it can be modulated by posttranslational modifications (PTMs) on histones, the reader proteins binding to them as well as by histone variant composition ^4^. In this context, destabilizing effects are expected in the case of activating functions, while nucleosome stabilization would suit repressive roles. In a puzzling manner, there are observations suggesting that the presence of H2A.Z in nucleosomes can increase ^5, 6, 7, 8, 9, 10, 11^, or decrease ^6, 12, 13, 14, 15, 16, 17, 18, 19^ nucleosome stability, which underscore the context-dependent complexity of this variant ^20, 21^. Even similar experimental approaches have led to disparate conclusions, such as the different findings from Förster resonance energy transfer measurements on reconstituted nucleosomes ^5, 20^ or magnetic tweezer unzipping measurements ^11, 15^. On the other hand, in genomics studies, destabilizing effects are regularly described. For example, the presence of H2A.Z destabilizes local nucleosome structure in ES cells, leading to decreased nucleosome occupancy and increased chromatin accessibility, particularly at enhancers ^16^. In line with this report, H2A.Z deposition creates wide ATAC-Seq positive promoter regions in mouse fibroblasts genome-wide ^18^. Furthermore, decreased unwrapping of the +1 nucleosomes has been found recently by MNase-X-ChIP-seq upon depletion of H2A.Z ^19^. Unusually labile (i.e. sensitive to low concentration of NaCl) H2A.Z-containing nucleosomes have been detected at the TSS (transcription start site) region of transcriptionally active promoters ^14^, particularly those flanking nucleosome-free regions at TSSs that contain H2A.Z paired with the H3.3 variant. Context-dependent variation of H2A.Z nucleosome stability has been demonstrated by a differential MNase digestion approach revealing that H2A.Z nucleosomes upstream or downstream of the TSS are much more resistant to MNase than those at the TSS ^22^. One possible explanation for some of the above differences is how nucleosome stability is defined and measured. On the other hand, since H2A.Z-containing nucleosomes are also present in the transcriptionally repressed heterochromatin ^23, 24^ with stability features likely different from those of the euchromatic localization, the different results may also be related to the possible intranuclear heterogeneity of H2A.Z nucleosomes.

In human cells, H2A.Z is loaded onto chromatin by the Tip60/p400 and SRCAP chromatin remodeling complexes, while ANP32E and INO80 are responsible for its eviction. In addition to the INO80 type remodelers, ISWI and CHD family protein complexes have also been implicated in the replication independent dynamics of the variant (for review see ^25^). These activities must be superimposed on and also influence, the nucleosome-autonomous stability features.

H2A.Z has two isoforms, H2A.Z.1 and H2A.Z.2.1, which differ in only 3 amino acids ^26^ and are encoded by separate genes. The two isotypes have indispensable and selective functions during development ^27, 28^ and appear to have distinct roles in controlling normal and cancer cell proliferation ^29, 30^. Only subtle structural differences between canonical and H2A.Z-containing nucleosomes were revealed by X-ray crystallography ^31, 32^. Alternative splicing gives rise to the hypervariant H2A.Z.2.2 that is shorter than H2A.Z.1 or Z2 by 14 amino acids and markedly destabilizes nucleosomes relative to H2A.Z.1 or H2A ^32^. Through the spectacles of mobility features of nucleosomal DNA termini, cryo-EM studies ^21^ support the notion of a destabilizing role of H2A.Z which depends on its C-terminal tail; however, the same study has also demonstrated the formation of more condensed chromatin fibers in the presence of H2A.Z (recently reviewed in ^33^). In a recent high-speed atomic microscopic (AFM) study ^34^, interaction between the N-terminal region of H2A.Z.1 and the DNA was found to be responsible for nucleosome sliding, at variance with ^21^. In summary, the different approaches addressing the relationships between the stability features of H2A.Z-containing nucleosomes and their other characteristics, including isoform composition, post-translational modifications (PTMs) and their roles in euchromatin vs. heterochromatin, have not yet led to a fully coherent picture ^2, 29^.

In this work, we made use of an *in situ* assay of nucleosome stability, QINESIn (Quantitative Imaging of Nuclei after Elution with Salt/Intercalators) ^35^ to gain insights into the stability features of H2A.Z-containing nucleosomes in close to native conditions of chromatin. This quantitative imaging cytometry-based assay delivers histone type, PTM- and cell cycle phase-specific information on the stability features of nucleosomes consisting of native endogenous or ectopically expressed histones, in populations of individual nuclei. A unique genetic complementation system ^36^ was also employed, involving H2A.Z.1 and Z2 double knock out (DKO) DT40 cell lines expressing transduced H2A.Z.1, Z2, a nonacetylatable Z1 mutant or a truncated H2A.Z.1 missing its C-terminal 9 amino acids ^26, 36, 37^, to learn what role the related factors may have in determining nucleosome stability *in vivo*. Having observed remarkable changes of nucleosome stability, nuclear architecture and accessibility features of chromatin in the case of the C-terminal deletion, the effect of a peptide representing the tail (C9) on these features was tested in permeabilized nuclei, and its binding to reconstituted nucleosomes was studies by fluorescence correlation spectroscopy (FCS). Based on these observations and the effect of C9 introduced into live cells on gene expression, a central role of the alternative engagements of the C-terminal H2A.Z tail in the functioning of this histone variant emerges, amenable to modulation by introducing C9 into live cells.

## RESULTS

### H2A.Z-containing nucleosomes exhibit intranuclear heterogeneity reflected by their stability features and intranuclear localization

Using QINESIn (Suppl. Fig.1A, B), we show that H2A.Z is unusually stably chromatin associated (Fig. 1A), exhibiting salt elution profiles typical of H3 or H4 (see ^35^, and refs. cited therein), as compared to canonical H2A or H2A.X, in every phase of the cell cycle (Fig. 1B). In QINESIn, the curves represent the average immunofluorescence of agarose-embedded nuclei remaining after their treatment with the indicated concentration of salt. The apparent stability of the H2A.Z-containing nucleosomes depends on which anti-H2A.Z antibody is applied to visualize the histone variant: an Abcam antibody (generated against the C-terminus of the histone starting with amino acid 65, comprising the H2A.Z docking domain and the C-terminal unstructured tail; see Suppl. Fig. 1C ; designated ZAbA from here on) and three others (Suppl. Fig. 2A-D) detect the stable H2A.Z-containing nucleosomes, while the Thermo Fisher Scientific antibody PA5-17336 (generated against a peptide sequence surrounding amino acid residue 118 of the docking domain, near the tail; designated ZAbB) detects H2A.Z in nucleosomes behaving similarly to those containing H2A (see dashed line in Fig. 1C), or to fluorescent protein-tagged H2A.Z (see below). (Since all the antibodies used detect both isoforms, the isoform-specific designations of the histone will be used in the text only when H2A.Z.1 and H2A.Z.2.1 could be distinguished.) Different antibodies detect H2A.Z at different nuclear localizations: ZAbA recognizes H2A.Z nucleosomes mainly at the nuclear periphery, while ZAbB staining gives a more scattered pattern (Fig. 1D, E). Suppl. Fig. 2E demonstrates that the H3-like stability of H2A.Z approached the H2A-like features when the nuclei were pretreated with a frequent-cutter nickase; we have previously shown that topological relaxation via nicking destabilizes nucleosomes to salt such that eviction of the dimers is facilitated ^35^. Thus, the same antibody is able to identify H2A.Z within stable or unstable nucleosomes depending on the superhelical state of the DNA. Antibody labeling follows the salt elution step in QINESIn, so it cannot affect the elution characteristics of a histone.

**Figure 1.**
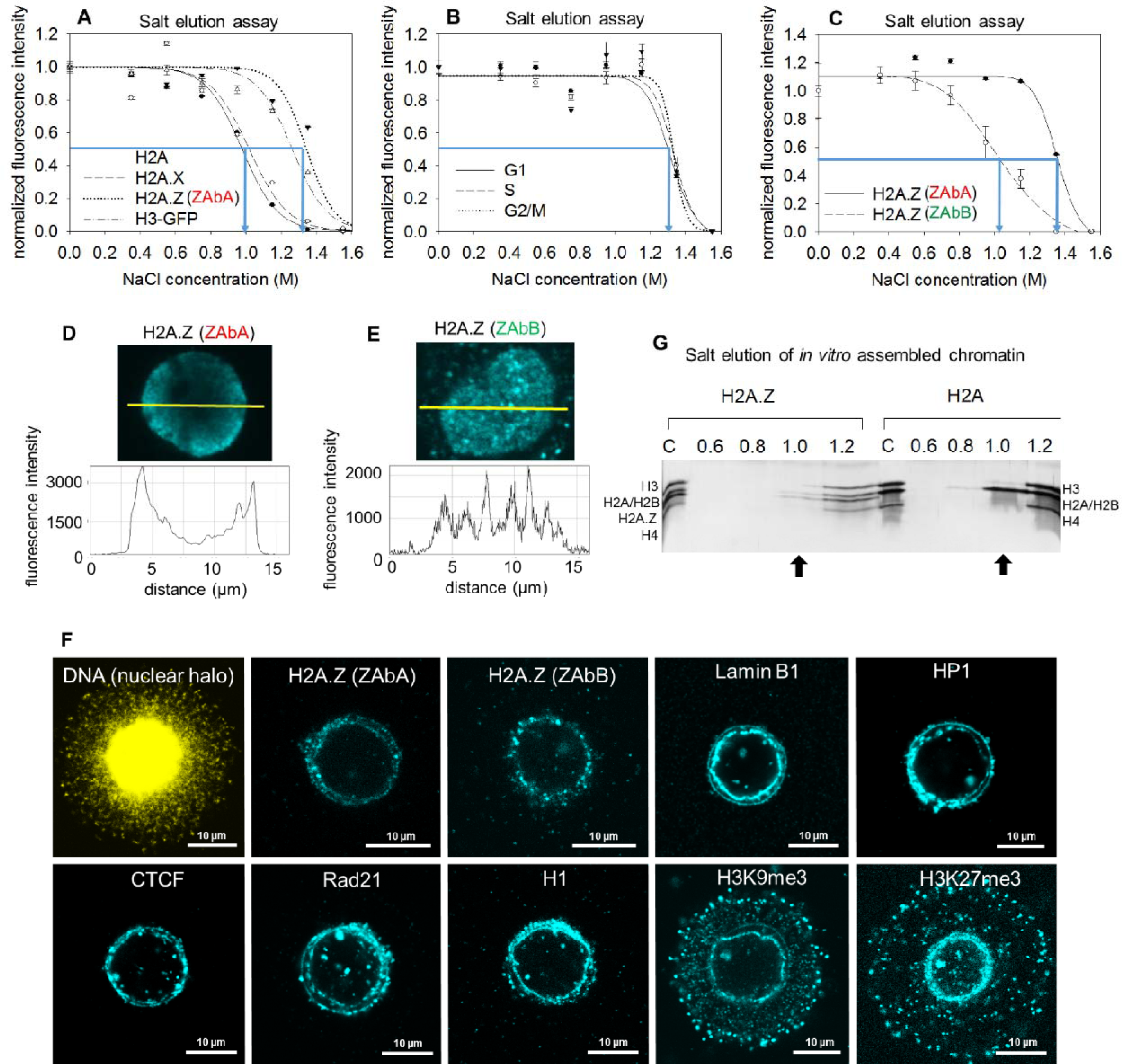
Intranuclear heterogeneity of H2A.Z. (A) Comparison of the salt elution profiles, measured by QINESIn (as in Suppl. Fig. 1A), of H2A, H2A.X, H2A.Z (detected by the antibody ZAbA; Abcam, ab 97966) and of H3-GFP (used as an internal control) in HeLa nuclei. (B) Salt elution profile of H2A.Z detected by ZAbA in HeLa nuclei of different cell cycle phases. (C) Salt elution curves of H2A.Z detected by ZAbB (Thermo Fisher Sci.) and ZAbA, measured separately in HeLa nuclei. In panels A-C, the elution curves refer to G1 phase nuclei gated according to their DNA fluorescence intensity distribution and the error bars represent SEM of ∼600 nuclei measured by LSC. Blue arrows on the elution curves indicate EC50 values (also in the other figures). (D, E) CLSM images and line-scans showing nuclear localization of H2A.Z as recognized by ZAbA (D), or by ZAbB (E). (F) Immunofluorescence staining of H2A.Z (using ZAbA or ZAbB), Lamin B1, HP1, CTCF, Rad21, H1, H3K9me3 and H3K27me3 in halo samples of HeLa nuclei. Representative images are shown. (G) Hydroxyapatite dissociation chromatography analyses of chromatin assembled *in vitro* using *Xenopus laevis* N1/N2-(H3, H4) and recombinant H2A/H2B or H2A.Z.1/H2B (see Materials and Methods). The H2A/H2B dimers run on the gel as a single band. See full gel image in Suppl. Fig. 3D. Arrows point at the histones eluted at 1 M salt.

H2A.Z isotype composition, PTMs that can mark H2A.Z, and the presence or absence of the C-terminus binding reader protein PWWP2A ^37^ could all affect the stability of nucleosomes containing H2A.Z. However, employing inhibitors and cell lines expressing different isoforms or mutants of H2A.Z in a double KO (DKO) DT40 background ^36^ we have shown that the stability features measured do not depend on H2A.Z isotype specificity, acetylation, ubiquitination or sumoylation (Suppl. Fig. 2G-L, Suppl. Fig. 3A-C). The finding that the H2A.Z elution curves were similar at different PWWP2A expression levels is not surprising in view of the fact that the reader is eluted at rather low salt, so it could not affect the stability features measured in our assay (see Suppl. Fig. 3C). Due to the same reason, the unusual stability features of H2A.Z as detected by ZAbA cannot be due to PWWP2 binding either. The above negative data are in line with the observations made on reconstituted nucleosomes (Fig. 1G). When the stability of the association of recombinant H2A.Z/H2B versus H2A/H2B dimer in nucleosomes was analysed by hydroxyapatite chromatography ^38^, a higher concentration of NaCl was required to elute H2A.Z as compared to H2A, being co-eluted with histone H3 (Fig. 1G, Suppl. Fig. 3D), recapitulating the salt elution profile in HeLa cells (Fig. 1A-C). These observations suggest that the unusual behavior of H2A.Z detected by several antibodies (Abcam 97966, 4174, 18262 and Millipore 07-594) is an autonomous feature of the nucleosomes containing the variant histone.

When the salt elution curves of Fig. 1A were plotted without normalization to zero endpoint (Suppl. Fig. 2F), a ∼10 % fraction of H2A.Z remaining in the nuclei after the highest concentration of salt could be detected. This fraction was apparently bound to the nuclear lamina according to the confocal laser scanning microscopic (CLSM) images of nuclear halo samples, as shown in Fig. 1F. H2A.Z was present in two concentric layers of the lamina, recapitulating its superresolution structure ^39^. This fraction is referred to from here on as H2A.Z^lmn^. Interestingly, beside H2A.Z, CTCF, H1, H3K9me3, H3K27me3, HP1 and the cohesin subunit Rad21 were also present in the two layers. Most of these proteins were also detected by mass spectrometric analyses of the halo samples (Suppl. Table 1).

Interestingly, the fluorescent protein-tagged forms of H2A.Z.1 and Z2 both show a destabilized character, similarly to what was observed through the spectacles of ZAbB, and this was invariant to the N- or C-terminal localization of the protein-tag (Suppl. Fig. 4A and B). As the co-labeling experiment of Suppl. Fig. 4C shows, the ZAbA-detected chromatin elements exhibit peripheral localization, while the tagged forms, similarly to the ZAbB-detected histones, are distributed in a scattered fashion in the nucleus, suggesting that ZAbA and ZAbB preferentially recognize different subsets of the variant in the nucleus. The intranuclear localization and stability features of H2A.Z-nucleosomes were both sensitive to even small tags, particularly when present at the C-terminus (Suppl. Fig. 4D-F). The specificity of the antibodies was demonstrated in Western blots using recombinant histones ^40^ (see Suppl. Fig. 5A-C), and also confirmed in experiments where silencing of H2A.Z using shRNA ^23^ has led to decreased H2A.Z immunofluorescence before cell death would occur (Suppl. Fig. 5D-G). H2A.Z silencing induced reorganization of the heterochromatin (Suppl. Fig. 5H, I), concomitant with the disappearance of the histone variant from the H3K9me3-rich peripheral heterochromatin (Suppl. Fig. 5G, H). Suppression of the binding of ZAbA by recombinant H2A.Z.1 protein was also demonstrated (Suppl. Fig. 5J). Silencing was confirmed also in western blots (Suppl. Fig. 6A-E) and by co-transfection of the silencing construct with a GFP-expressor plasmid allowing the separate gating of successfully transfected cells in the laser scanning cytometric (LSC) analysis of H2A.Z expression (Suppl. Fig. 6F-H).

Thus, two kinds of H2A.Z containing chromatin regions can be distinguished: ZAbA-detected salt-stable ones likely associated with the peripheral heterochromatin (designated H2A.Z^hc^; see Fig. 1D), and those with stability features of canonical nucleosomes being more scattered or more centrally located in the nuclei, resembling euchromatin (designated H2A.Z^eu^; see Fig. 1E). (The third nuclear compartment containing the variant histone is H2A.Z^lmn^; see above.)

The localization patterns of the histone variant in the agarose-embedded nuclei labeled with ZAbA and ZAbB before fixation are similar to those of the prefixed specimen shown in the manufacturers’ datasheets. The CFP or YFP-tagged H2A.Z.1 and Z2 containing nucleosomes appear to be scattered in the nucleus ^41^, similarly to the images obtained with ZAbB and the GFP-tagged histone (Fig. 1E, Suppl. Fig. 4C).

In agreement with the ZAbA-detected nucleosomes exhibiting H3-like, i.e. decreased, salt-sensitivity in the salt elution experiments, H2A.Z was detected by mass spectrometry (MS) in the salt resistant fraction (see Suppl. Fig. 7A and Suppl. Table 2). Proteins associated with euchromatic functions as well as others implicated in heterochromatin organization ^42, 43^ were both detected, just like in a different experimental set-up, using the CUT&RUN protocol ^44^ followed by MS (flow-charts and titration of MNase concentration are shown in Suppl. Fig. 7B and C-E, respectively). According to the MS data, ZAbA and ZAbB recognize nucleosomes associated with an overlapping spectrum of proteins, many of which were found to be H2A.Z-associated also in other studies. There was an overlap also between the list of hits in the two different approaches (H2A.Z, HP1, MATR3, LMNA, H1.0).

### Effect of the H2A.Z C-terminus on nucleosome stability and localization

In view of the fact that the C-terminal region of the unstable H2A.Z.2.2 isoform is 14 residues shorter than either isoform of H2A.Z ^4^, where the last six residues of Z.2.2 form a motif facilitating nucleosome assembly ^45^, we conducted experiments with cells expressing H2A.Z.1 with a 9 amino acid deletion at the end of their C-terminus (between amino acids 119-128, Suppl. Fig. 1C), on a double, H2A.Z.1, H2A.Z.2 KO (DKO) DT40 background ^36^. The H2A.ZΔC containing nucleosomes were less stable as compared to nucleosomes containing full-length H2A.Z, as studied by QINESIn in the agarose-embedded nuclei of the H2A.ZΔC and the H2A.Z.1-expressor control cells, respectively; Fig. 2A, Suppl. Fig. 8A), while the stability of nucleosomes carrying H3K27me3 or H3K9me3 was not affected significantly (Suppl. Fig. 8B, C). The elution profile of the C-terminally truncated H2A.Z-nucleosomes exhibiting canonical, H2A-like behavior was affected much less by nicking treatment, as compared to the nucleosomes containing full-length H2A.Z (using ZAbA; Suppl. Fig. 8D).

**Figure 2.**
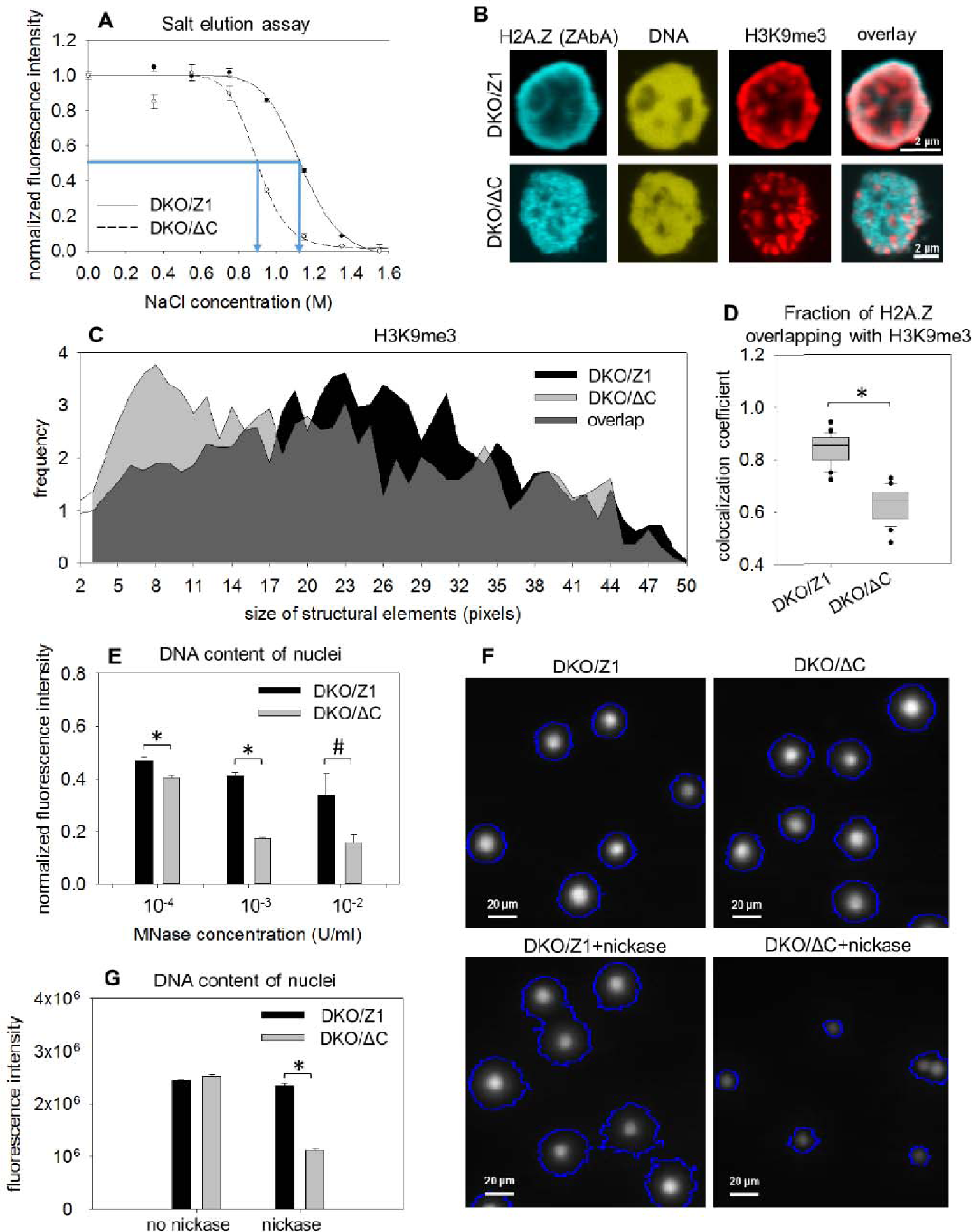
Effect of C-terminal truncation on the salt sensitivity, nuclear localization and chromatin accessibility features. (A) Salt elution profiles of nucleosomes in H2A.Z.1ΔC (DKO/ΔC) and H2A.Z.1 expressor DKO DT40 cells (DKO/Z1), detected by ZAbA. The elution curves refer to G1 phase nuclei gated according to their DNA fluorescence intensity distribution and the error bars represent SEM of ∼600 nuclei measured by LSC. (B) Representative CLSM images showing the nuclear localization of H2A.Z recognized by ZAbA, and H3K9me3 co-labeled with H2A.Z in DKO H2A.Z.1ΔC (DKO/ΔC) and H2A.Z.1 (DKO/Z1) nuclei. (C) Texture analysis (see Materials and Methods) of ΔC (DKO/ΔC) and CTRL (DKO/Z1) nuclei showing the size distribution of structural elements containing H3K9me3. (See also Suppl. Fig. 11A-D). (D) Colocalization analysis of H2A.Z and H3K9me3 in ΔC (DKO/ΔC) and CTRL (DKO/Z1) nuclei. The Manders colocalization coefficient reflecting the fraction of H2A.Z overlapping with H3K9me3 are shown. Box-and-whisker plot was created from the data of 25 nuclei. Statistical analysis was done using one-way ANOVA (* p≤0.001). For the expression levels of the proteins in the two cell lines, see Suppl. Fig. 11E. (E) MNase sensitivity of ΔC (DKO/ΔC) and CTRL (DKO/Z1) nuclei. Total DNA content per nuclei was measured by LSC before and after endonuclease treatment, as indicated on the figure. (F and G) Comparison of chromatin sensitivities to a frequent cutter nickase (Nt.CviPII). DNA content (G) and halo size (Suppl. Fig. 13B) of ΔC and CTRL nuclear halos were measured by LSC before and after nickase treatment of the nuclei (see Materials and Methods), as indicated on the figure. Error bars represent SEM of ∼600 nuclei. Statistical analysis was done using one-way ANOVA (* p≤0.001, # p≤0.05). Box-and-whisker plot shows the median, 25^th^ and 75^th^ percentiles as vertical boxes with error bars, 5^th^, 95^th^ percentiles and outliers as dots (also in the other figures).

Remarkably, not only the stability of H2A.Z containing nucleosomes but also their nuclear localization pattern was C-terminus dependent (Fig. 2B-D). In the H2A.ZΔC-expressing cells, the localization of ZAbA-detected H2A.Z is changed to a much more scattered and less peripheral topography in the same DKO background, while there was no such difference in the case of the ZAbB-detected ones (compare Suppl. Fig. 9 with Suppl. Fig. 10). The spatial distribution of H3K9me3 marked nucleosomes also changed, as reflected by the altered texture of H3K9me3 containing structural elements (Fig. 2C; Suppl. Fig. 11A,B) in comparison with that of H3K27me3 (Suppl. Fig. 11C, D), and the colocalization data of Fig. 2B, D, Suppl. Figs. 9-10. Manders colocalization coefficient (MCC) was used to characterize the pixel-overlap of H2A.Z with hetero- and euchromatin markers. The H2A.ZΔC levels expressed in the nuclei of these DKO DT40 cells were about half of the H2A.Z levels in the same DKO background, while the total number of H3K9me3, H2B nucleosomes and also the amount of DNA in the nuclei of the two cell types were similar (Suppl. Fig. 11E). The degree of colocalization between ZAbA-detected H2A.Z and H3K9me3 was significantly lower in the H2A.ZΔC than in H2A.Z.1 DKO nuclei (Fig. 2D). The difference between the two cell lines was smaller in the case of H2A.Z and H3K27me3 colocalization (Suppl. Figs. 9-10). The somewhat elevated levels of H3K27me3 in H2A.ZΔC cells (Suppl. Fig. 11E) may account for the altered degree of colocalization between H3K9me3 and H3K27me3 (Suppl. Fig. 10D). The conspicuous decrease of colocalization between H2A.Z and H3K9me3 in H2A.ZΔC is present in spite of the fact that the ZAbA-detected H2A.ZΔC levels are lower than those of H2A.Z detected in DKO/Z1 cells (Suppl. Fig. 11E). The higher Manders coefficient indicative of more H2B colocalizing with ZAbA-detected H2A.Z in the ΔC nuclei relative to those harboring the full-length form (Suppl. Fig. 9C) can be explained by the decrease in peripherally localized H2A.Z levels, but not of H2B. At the same time, there was no change in the colocalization coefficients in the case of PWWP2A and H2B (Suppl. Fig. 9D).

In contrast with the above observations, when the histone variant was labeled using ZAbB, preferentially recognizing euchromatic H2A.Z as shown above, no significant differences were measured between the two kinds of cells in terms of colocalization of the histone variant with either the euchromatic mark H3K4me3 or the heterochromatic mark H3K27me3 (Suppl. Fig. 10A, B). Although the ZAbB-detected H2A.Z was similarly scattered in both wild-type (wt) and ΔC nuclei, the fraction of ZAbB-detected H2A.Z colocalizing with H3K9me3 decreased significantly in the ΔC nuclei (Suppl. Fig. 10C), similarly to the decrease of colocalization between ZAbA-detected H2A.Z and H3K9me3 (Fig. 2D). Furthermore, the degree of colocalization between the two heterochromatic marks H3K27me3 and H3K9me3 was significantly different in the H2A.Z.1 vs ΔC nuclei (Suppl. Fig. 10D), suggesting that the global nuclear organization has changed in the ΔC nuclei. The topographic characteristics of ZAbA- and ZAbB-detected H2A.Z in the DKO/Z1 and H2A.ZΔC nuclei described above were also apparent in superresolution microscopic images, as shown in Suppl. Fig. 11F, G. There was a difference between the wt and ΔC DT40 nuclei also in the distribution of H3K9me3 remaining lamina-attached in the halo samples, while no such difference was detected in the case of H3K27me3 (Suppl. Fig. 12).

### Consequences of H2A.Z C-terminal truncation on accessibility of DNA in chromatin and on gene expression

The architectural changes related to the C-terminal truncation of H2A.Z were also reflected in global effects of functional relevance. As compared to DKO H2A.Z.1 nuclei, those of the DKO H2A.ZΔC cells were more readily digested using a frequent cutter nickase, MNase and DNAse I (Fig. 2E-G and Suppl. Fig. 13A, B). These H2A.Z tail-dependent changes in accessibility features were also reflected in differences of gene expression patterns, comparing wt, DKO H2A.Z.1 and DKO H2A.ZΔC cells (Suppl. Fig. 14). Among the genes substantially up- or down-regulated in DKO H2A.ZΔC relative to DKO H2A.Z.1 or wt, those that have been implicated with pathways where H2A.Z is known to participate in, are listed in Suppl. Table 3. These pathways include processes directly involving H2A.Z such as DNA repair ^46^. Indeed, DKO H2A.ZΔC cells appear to exhibit a diminished DNA damage response and relatively poor survival upon exposure to DNA damaging agents (Suppl. Fig. 15).

### Effects of C9 treatment of permeabilized nuclei

As Fig. 3, Suppl. Fig 13C-E, Suppl. Figs. 16 and 17 demonstrate, the destabilized character, increased nuclease sensitivity and altered intranuclear distribution of nucleosomes containing H2A.ZΔC were emulated when a peptide representing the last 9 amino acids of the C-terminus (C9) was added to nuclei, suggesting that these differences are the direct consequences of molecular interactions involving the tail region; shorter peptides (C6-C8) or a scrambled control peptide (SCR; see Materials and Methods) were not, or much less effective. Upon treatment with C9, the H2A.Z-containing nucleosomes fell apart to a higher extent at 1 M salt (Fig. 3A), while addition of the peptide to nuclei at low salt elicited no eviction (Suppl. Fig. 16A, B). The H2A.Z-containing chromatin regions became partially separated from the H3K9me3-rich periphery based on the significantly decreased Manders coefficient (Fig. 3B, C, Suppl. Fig. 16C, D) and treatment of the nuclei with the C9 peptide increased sensitivity to nickase and MNase (Fig. 3D-F, Suppl. Fig. 17A, B) used at 10^−3^ U/ml concentration (see Fig. 2E). This MNase concentration is in the lower end of the usual concentration range of the enzyme applied in MNase-seq studies when performed on fixed ^47^ or nonfixed cells ^48^; however, the enzyme was applied in the absence of loosely bound proteins that were released upon preparation of the nuclei in our studies.

**Figure 3.**
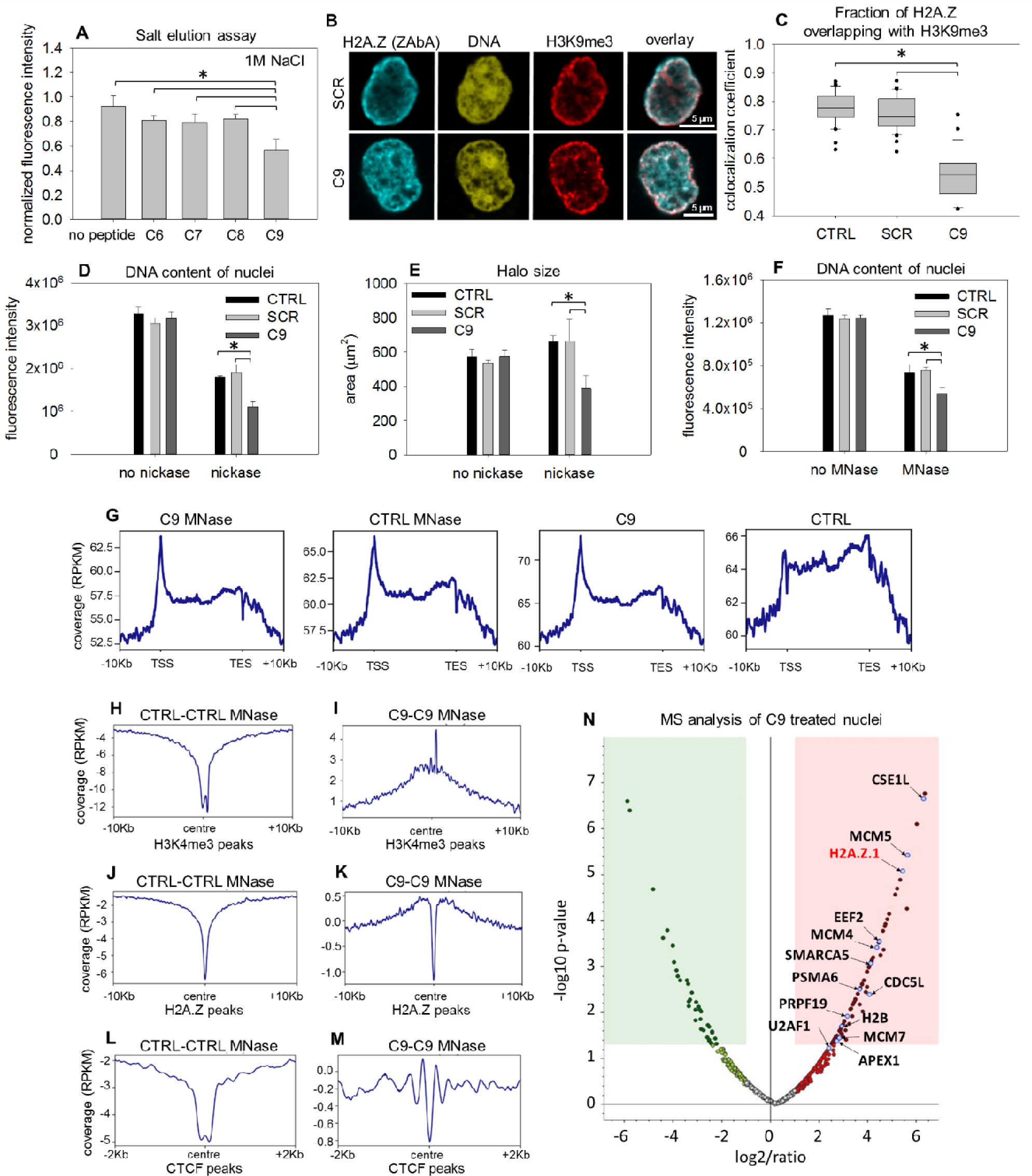
The effect of peptides containing the last 6-9 amino acids of the H2A.Z.1 C-terminus on the stability, nuclear localization and MNase digestability of H2A.Z.1-containing nucleosomes. (A) Stability of H2A.Z nucleosomes in permeabilized HeLa nuclei exposed to 1 M NaCl in the presence of different length peptides (C6-C9). ∼600 G1 nuclei were measured in each samples by LSC. Error bars represent SD of 3 parallel measurements. Statistical analysis was done using one-way ANOVA (* p≤0.001) (B) Representative CLSM images showing nuclear localization of H2A.Z recognized by ZAbA and of H3K9me3 co-labeled with H2A.Z, in HeLa nuclei treated with C9 or SCR. (C) Colocalization analysis of H2A.Z and H3K9me3 on CLSM images from experiment on panel (B). The Manders colocalization coefficients representing the fraction of H2A.Z overlapping with H3K9me3 are shown. Box-and-whisker plot was created from the data of ∼30 nuclei. Statistical analysis was done using one-way ANOVA (* p≤0.001) (D, E) Global nickase sensitivity of HeLa nuclei after treatment with C9 or SCR, measuring DNA content based on SYBR Gold staining or halo size, by LSC. Statistical analysis was done using one-way ANOVA (* p≤0.001) (F) Global MNase sensitivity of HeLa nuclei after treatment with the scrambled control peptide (SCR) or with C9 (C9), measuring SYBR Gold-stained DNA content with LSC. Error bars represent SD of 4 parallel measurements, where events were collected from the whole cell population. Statistical analysis was done using one-way ANOVA (* p≤0.001) (G) Metaplots of sequencing read coverages over the regions spanning the interval between the TSS and TES of the genomic DNA prepared from C9 treated, C9-untreated and MNase-digested or MNase-undigested samples (see also Suppl. Fig. 17A). The Y scales maximize resolution in each metaplot. (H-M) Anchor plots of CTRL – CTRL(MN) (H, J, L panels) and C9 – C9(MN) read coverage differences (I, K, M panels) around H3K4me3 (H and I), H2A.Z (J and K) and CTCF (L and M) peaks. (N) Volcano plot illustrating protein fold changes in the C9-treated sample compared to the untreated control identified by mass spectrometry in a ZAbA-MNase-pA/G-CUT&RUN experiment using HeLa nuclei (see Materials and Methods). Proteins listed in Suppl. Table 4 are highlighted.

In order to determine which chromatin regions were sensitized preferentially by C9 to MNase, the DNA remaining in the agarose-embedded nuclei after digestion was isolated, sequenced and the distribution of read coverages were compared. Note: we sequenced the DNA remaining in the agarose-embedded nuclei after digestion (see Materials and Methods), differently from the ‘MNase-seq’ procedures where the DNA of the whole chromatin digest is the input for sequencing, among the several other differences (see e.g ref. 48). The MNase-digested and undigested samples containing identical amounts of DNA were sequenced yielding approximately identical overall coverage; therefore, only the shape of the curves, not their relative position along the Y axis, is compared. As the metaplot analyses of Fig. 3G shows, coverage is increased at the TSS relative to the gene bodies upon C9 treatment, indicating decondensation of chromatin ^49^; there was no further change of the shape of the metaplot following MNase treatment as judged by the naked eye. When the effect of C9 on the read coverage-differences between MNase digested (MN) and undigested (CTRL) nuclei, i.e. on MNase accessibility, were calculated, conspicuous changes were detected at large regions and also at particular sites; see Suppl. Fig. 17C-K. C9 had an effect on MNase-digestibility around the H3K4me3 nucleosomes corresponding to TSS, the promoter-proximal H2A.Z peaks (compare Fig. 3H-K and Suppl. Fig. 17L-O) and also at the binding sites of the insulator protein CTCF (Fig. 3L, M). In both of the above locations MNase sensitivity was increased and its pattern characteristically changed. The effect of C9 treatment on MNase digestability over the total genomic DNA was closely recapitulated when the read coverages over the LINE1 repetitive elements, the H3K9me3-rich regions or the centromeric regions were considered, while the peptide had no effect on MNase digestion over the ALU repeats and H3K27me3-rich regions (Suppl. Fig. 17C). Thus, MNase sensitivity was increased at specific sites in euchromatin and certain large areas of heterochromatin were also sensitized to the nuclease upon treatment of nuclei with C9.

The result of C9 treatment of HeLa nuclei was also studied by ZAbA-targeted CUT&RUN (see Materials and Methods) combined with mass spectrometry. As Fig. 3N and Suppl. Table 4 demonstrate, an increased presence of several nuclear proteins associated with euchromatic functions, including H2A.Z, was observed, in line with decondensation and increased accessibility to ZAbA and/or to MNase in the chromatin around H2A.Z-containing nucleosomes in the euchromatin. The H2A.Z-nucleosomes of peripheral heterochromatin remained in the nucleus in these experimental conditions (Suppl. Fig. 17P).

### Interaction of reconstituted nucleosomes with C9 as studied by fluorescence correlation spectroscopy (FCS)

FCS measurements were conducted to further investigate to what extent the carboxyfluorescein-labeled C9 (C9-CF) is able to bind reconstituted H2A or H2A.Z-containing nucleosomes and/or the Cy5-labeled nucleosome-positioning Widom sequence used for reconstruction.

The autocorrelation function (ACF) of the samples with nucleosomes was shifted to longer lag times suggesting slower diffusion due to binding (Fig. 4A, B). The fits of the ACFs allowed us to identify a fast-diffusing population in all samples corresponding to non-bound peptides and a slower diffusing population in the nucleosome containing samples attributed to peptides bound to nucleosomes. A slow component was also observed in samples containing naked DNA. The relative proportion of the slow, bound component was also obtained from the fits (Fig. 4C, Suppl. Fig. 18). The ACF curves of C9-CF alone could be fitted reasonably well with a single diffusing component (see Fig. 4B); therefore, no C9-C9 binding is assumed to occur in our conditions.

**Figure 4.**
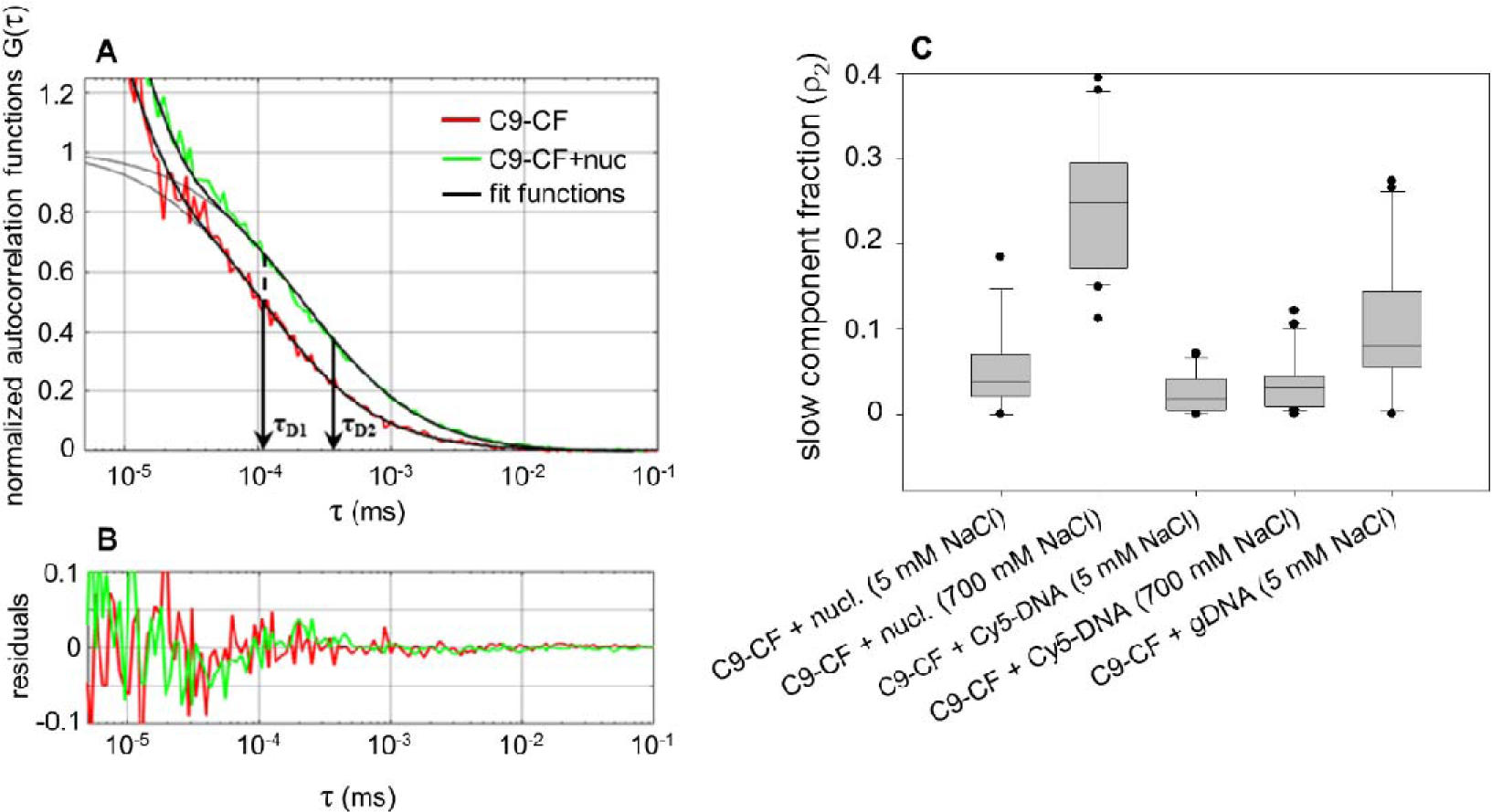
FCS analysis of C9-CF binding to reconstituted mononucleosomes. (A) Representative normalized autocorrelation functions (ACF-s) of C9-CF are shown; the correlation curves were acquired in the absence (red) or presence (green) of nucleosomes, at 700 mM NaCl concentration. Fits assuming triplet transition and a single diffusing species for a sample without nucleosomes and two diffusing populations with nucleosomes were used (black solid lines). The grey line indicates the fit excluding triplet state transition. τ_D1_and τ_D2_ are the diffusion times of the fast and slow components, characterizing the diffusion of freely diffusing and nucleosome-bound C9-CF peptides, respectively. The ACF-s shifted toward slower diffusion times indicating the presence of a nucleosome-bound fraction. (B) Fit residuals. (C) Slow fraction of C9-CF. The peptide was incubated with nucleosomes at 5 mM or 700 mM NaCl concentration, with Cy5-labeled Widom-sequence at 5 mM or 700 mM NaCl, or with genomic DNA at 5 mM NaCl. The slow fraction ρ_2_ was calculated in ACF fits as described in the Materials and Methods and presented as a box-and-whisker plot.

In fitting the autocorrelations curves, the diffusion time of the slow fraction was determined based on the diffusion of the Cy5-labeled DNA, after correction for the difference in focal volumes at the different wavelengths (following the pipeline shown in Suppl. Fig. 19). These measurements were conducted at equilibrium conditions established at a 5-fold molar excess of nucleosomes or DNA over C9-CF (or the labeled scrambled peptide, used as a control).

As Fig. 4C demonstrates, at low salt condition about ∼5 % of the peptide was found in the slower fraction representing C9-CF bound to any of the macromolecules, in several independent experiments. The CF-labeled scrambled control peptide did not bind to nucleosomes or DNA, while binding of C9-CF to nucleosomes built up from either canonical H2A or H2A.Z was well detectable (Suppl. Fig. 18).

From the 5 % value, a K_d_ of ∼2 µM was calculated based on the law of mass action (as in ^50^). When the FCS measurements were conducted in solutions containing 700 mM NaCl, expected to loosen the binding of the canonical histone dimers ^51^, binding of the peptide to the nucleosomes was considerably increased, yielding ∼35 % of the peptide in the slower fraction, which translates into a K_d_ = 90 nM. Binding of the peptide to DNA was not increased significantly at this salt concentration. These data raise the possibility that the C-terminal tail may be involved in establishing intra- or internucleosomal associations.

### Effects of C9 introduced into live cells

Under conditions that did not significantly affect cell viability (Suppl. Fig. 20), C9-CF could be efficiently introduced into live HeLa cells using the cyclodextrin derivative SBECD as shown in Fig. 5A and Suppl. Fig. 21. The dotted CLSM pattern of C9 accumulation when added to live cells (observed using C9-CF; Fig. 5A) is probably the consequence of the uptake process. The nuclear features that appear to depend on the C-terminus of H2A.Z were susceptible to modulation when C9 was introduced into live cells. As Fig. 5B, C and Suppl. Fig. 21A-F demonstrate, reorganization of peripheral H2A.Z-containing heterochromatin relative to H2B-GFP or H3K9me3, decreased nucleosome stability (Fig. 5G, Suppl. Fig. 21G) and increased nuclease (nickase) sensitivity (Fig. 5E, Suppl. Fig. 21H) could be detected in the nuclei of C9-treated HeLa cells one day after a single dose of peptide treatment. Similar though less conspicuous changes were demonstrated in the nuclei of two melanoma cell lines (^30, 52^; see Fig. 6A,B), with the reorganization of H2A.Z-chromatin demonstrated via its altered colocalization with H3K9me3. When the effect of C9 treatment was studied in MEL1617 cells by RNA-seq, 564 genes were significantly down-regulated and 58 up-regulated (Fig. 6C). Interestingly, almost none of the down-regulated genes localize to the Giemsa-dark bands in the chromosome ideograms. At the same time, genes of different expression levels in the C9-untreated control cells and the H2A.Z ChIP-seq peaks of (SK-MEL-147) melanoma cells are rather evenly distributed among the Giemsa-light and dark bands (Fig. 6E-K). Gene subsets of particular pathways are overrepresented among the significantly down-regulated genes (Fig. 6L).

**Figure 5.**
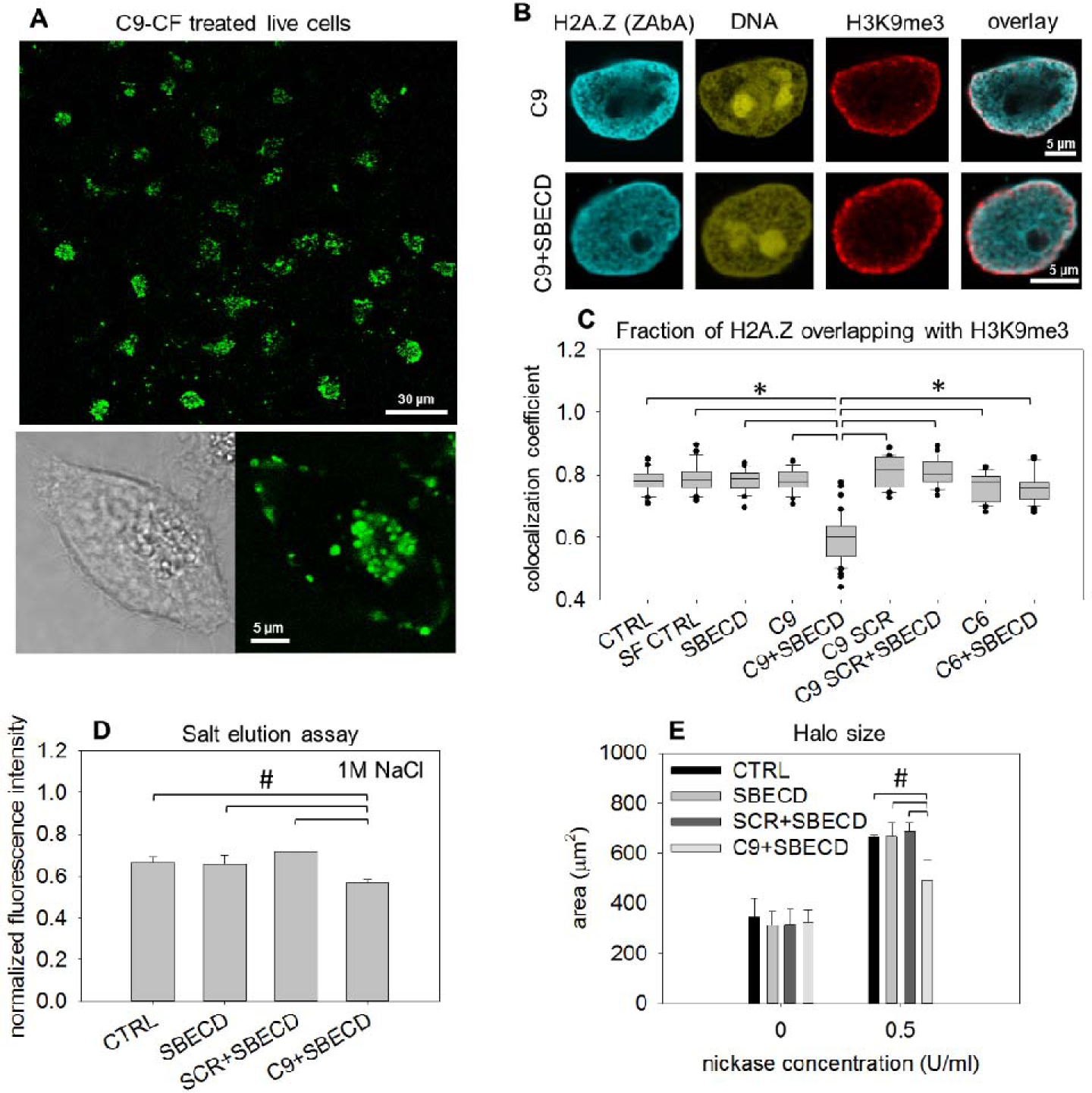
Localization and effects of C9 peptide introduced into live HeLa cells by the cyclodextrin derivative SBECD. (A) Localization of C9-CF (carboxyfluorescein conjugated C9) in live HeLa cells after 2 h treatment with C9-CF/SBECD followed by overnight culturing, visualized by CLSM. Zoom-n image of a cell from the C9-CF/SBECD treated sample is shown at the bottom. (B) Representative confocal images of H2A.Z-H3K9me3 co-labeled nuclei following introduction of C9 into live HeLa cells. Cells were treated with the C9, C9 SCR or the C6 peptide using SBECD (C9+SBECD, C9 SCR+SBECD or C6+SBECD), or with serum-free medium in the absence of peptides and cyclodextrins (SF CTRL). (C) Manders colocalization coefficients representing the fraction of H2A.Z overlapping with H3K9me3 calculated for the nuclei shown in panel B. Box- and-whisker plots were created from the data of ∼30 nuclei. Error bars represent SD of four parallel measurements. Statistical analysis was done using one-way ANOVA (* p≤0.001). (D) Resistance of H2A.Z nucleosomes to 1 M NaCl in permeabilized HeLa nuclei prepared from untreated, cyclodextrin-treated, SCR+SBECD- or C9+SBECD-treated HeLa cells. Bar charts show the mean fluorescence intensity, error bars represent the SD of 3 parallel measurements. Statistical analysis was performed using one-way ANOVA (# p≤0.025). (E) Measurement of chromatin sensitivity to nickase. Halo size of untreated (CTRL), SBECD-, SCR+SBECD- and C9+SBECD-treated nuclei were measured by LSC before and after nickase treatment. Bar charts show the mean fluorescence intensities, error bars represent the SD of 3 parallel measurements. Statistical analysis was done using one-way ANOVA (# p≤0.05).

**Figure 6.**
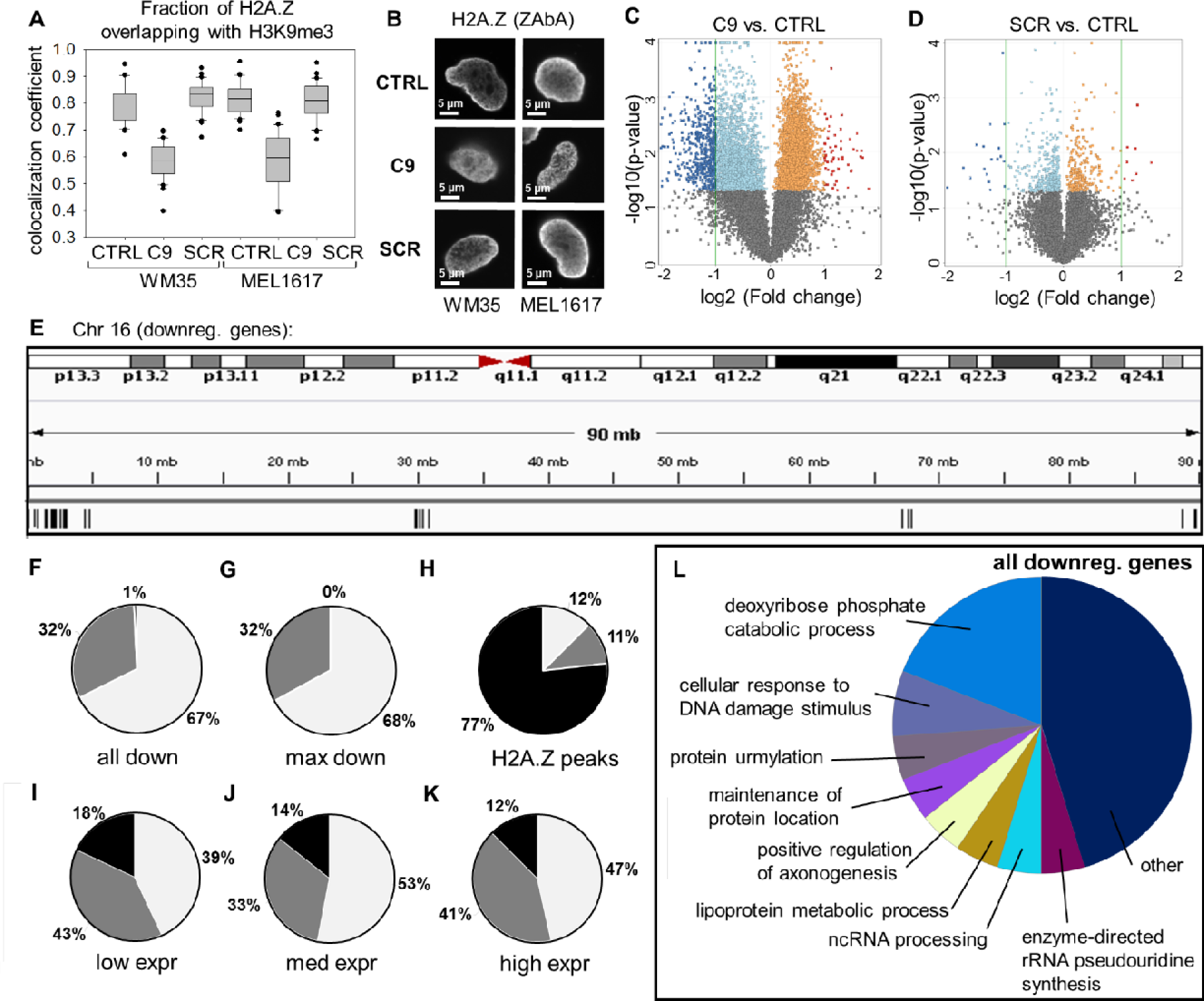
Effect of C9 treatment of live melanoma cell lines. (A) Colocalization of H2A.Z and H3K9me3 in WM35 and MEL1617 cells treated with C9, SCR or left untreated (CTRL). Manders colocalization coefficients representing the fraction of H2A.Z overlapping with H3K9me3. Box-and-whisker plots were created from the data of 30 nuclei. (D) CLSM images from the samples of panel (A) showing H2A.Z rearrangement. (C) and (D) Volcano plots showing differentially expressed genes defined as log2 fold change < or > 1.0 and p < 0.05 (dark blue and red dots). C9 vs. CTRL (C) and SCR vs. CTRL (D) data of MEL1617 cells are shown. (See the volcano plot of the differentially expressed genes in SCR vs. C9 in Sup l. Fig. 22.) (E) IGV screenshot of the chromosome 16 ideogram. The genes down-regulated by 9 treatment are shown below. (F) Pie chart showing the distribution of all down-regulated genes y C9 treatment among different Giemsa bands. (G) Pie chart showing the distribution of highly down-regulated genes (fold change < –2.35; ∼1/3 of all down-regulated genes) among different Giemsa bands. (H) Pie chart showing the distribution of HeLa H2A.Z ChIP-seq peaks (downloaded from ENCODE database, accession ENCFF094MFL) among different Giemsa bands. (I)-(K) Pie charts showing the distribution of all expressed genes grouped by the expression level of the corresponding gene. Distribution of genes with low (< –0.2), medium (– 0.2–0.5) and high (> 0.5) expression in control MEL1617 cells are shown in panel (I), (J) and (K), respectively. The distributions represented by the pie charts are normalized to the fraction of the corresponding areas in the genome. (L) Functional characterization of the genes down-regulated by C9 (C9 vs SCR).

## DISCUSSION

The data presented demonstrate the decisive role of the C-terminal unstructured tail of H2A.Z both in determining stability features of nucleosomes containing this variant and in their intranuclear landscape. Further, this study reveals the H2A.Z tail-dependence of certain local and global chromatin accessibility characteristics and offers means to modulate chromatin structure and function via interfering with the interactions involving its tail.

### H2A.Z tail-dependent changes in euchromatin and heterochromatin

The elution curves of QINESIn reflect the off-rate of particular histones released from the nucleosomes *in situ*, i.e. we look at nucleosome stability through the spectacles of H2A.Z dissociation, native or tagged, induced by salt treatment. Based on the localization patterns (see e.g. Fig. 1D, E), the tail-dependent changes and differences thereof (e.g. as shown in Fig. 2B-D, Fig. 3B, Fig. 5B, Fig. 6B, Suppl. Fig. 11F, G), colocalization with HP1 (Suppl. Fig. 10F), and the different stability features (Fig. 1A-C, Fig. 2A, Fig. 3A, Fig. 5D), we propose that ZAbA preferentially detects stable nucleosomes residing in peripheral heterochromatin (H2A.Z^hc^), while ZAbB recognizes the histone variant preferentially in unstable nucleosomes within the euchromatin (H2A.Z^eu^). Shielding of particular H2A.Z epitopes by the different molecular partners specific to the compartments could account for the different reactivity of the two classes of antibodies. The stable H2A.Z containing nucleosomes become destabilized to a canonical H2A-like state when the C-terminal tail is deleted or its binding to molecular partners may be disabled by the addition of a competing peptide. The tags apparently also hamper some of these interactions, with a destabilizing effect (Suppl. Fig. 4F). The N-terminal part of H2A.Z has a marked effect on the sliding of the nucleosome on the DNA ^34^, while the C-terminal tag is expected to directly interfere with interactions involving the tail. Tagged H2A.Z also appears to be localized in a scattered rather than dominantly peripheral manner in the nucleus, compared with ZAbA-detected native H2A.Z (Suppl. Fig. 4C, E).

The remarkable tail-dependence of chromatin organization as detected by ZAbA or anti-H3K9me3 staining suggest that these changes certainly involve heterochromatin. Sensitization of LINE1, centromeric and H3K9me3-rich regions to MNase by C9 treatment (Suppl. Fig. 17C) also shows that heterochromatin is affected by the presence of the tail and can be manipulated by treatment with excess C9. On the other hand, the gene expression differences between wt or DKO H2A.Z.1, and H2A.ZΔC-expressor DKO DT40 cells (Suppl. Fig. 14, Suppl. Table 3), the effect of C9 treatment on MNase sensitivity at TSS regions (Fig. 3F) and on the release of proteins associated with transcription, replication and repair (Fig. 3N, Suppl. Table 4) in the case of nuclei, and of the changed expression of altogether ∼620 genes in the case of live cell treatment (Fig. 6) collectively imply H2A.Z tail-dependent changes to be present also in euchromatin.

ChIP sequencing (ChIP-seq) data available for different cell types reveal that two of the antibodies detecting unusually stable nucleosomes (Abcam 4174 and Millipore 07-594; see Suppl. Fig. 2A and C, respectively, and refs. ^53, 54^) give peaks both in euchromatin and heterochromatin. The ChIP-seq data obtained using H2A.Z-GFP and anti-GFP antibodies were very similar to parallel ChIP-seq data obtained with Abcam 4174 ^52^. However, the experimental conditions of ChIP-seq and immunofluorescence studies are very different, as antibody labeling occurs after sonication in the first, while it is applied in the close-to-native state of the agarose-embedded, permeabilized nuclei in the second. Indeed, the distribution of H2A.Z reflected by CUT&RUN mapping, where antibody labeling is performed among conditions comparable to those of our elution assay, exhibits only partial overlap with ChIP-seq datasets ^55^. In view of the different preparation steps and also the ensemble character of the above genomic approaches, those data are not expected to reflect the picture obtained by immunolabeling using ZAbA or ZAbB.

### The presence of H2A.Z in the nuclear lamina

The fraction of the histone variant remaining in the nuclei after the maximal salt concentration used in the elution experiments (Suppl. Fig. 2F) was further investigated by mass spectrometric analyses and immunostaining of these nuclear halo preparations (see Materials and Methods). As Fig. 1F and Suppl. Table 1 show, a fraction of H2A.Z (H2A.Z^lmn^), as well as proteins implicated in laminar tethering and/or chromatin loop organization, could be detected. All these proteins appear to be in tight association with the nuclear lamina. We speculate that H2A.Z^lmn^ can interact with the nucleosomes and/or the internucleosomal DNA of peripheral heterochromatin, contributing to the laminar tethering of chromatin loops. The decreased presence of lamina-associated H3K9me3 signal in the ΔC DT40 cells (Suppl. Fig. 12) is in line with this hypothesis.

We note that the prominent H3K9me3 and H3K27me3-rich spots at the periphery of the nuclear halos give the impression of loop anchorage points detached from the lamina upon salt exposure (Suppl. Fig. 12). H3K9me3 detected at the periphery of the halos were more prominent in the case of the full-length H2A.Z containing nuclei as compared with those of the ΔC-expressors, suggesting involvement of tail-mediated molecular interactions in higher-order chromatin organization. Remarkably, the peculiar pattern of the changes in read coverage upon MNase digestion at CTCF sites (Fig. 3M) is strongly reminiscent of the unwrapping pattern of H2A.Z nucleosomes at the CTCF binding sites; see Fig. 3 of ref. ^19^. This observation raises the possibility that the reorganization of heterochromatin upon C9 treatment may be involved in the disruption of loop anchorage involving CTCF.

The separation of the two layers of the nuclear lamina with different Lamin B1 content (Fig. 1F) confirm the conclusions of the superresolution studies first describing this structural feature ^39^. This simple way of visualization, also in view of the laminar presence of proteins involved in higher-order chromatin organization (Suppl. Table 1), opens possibilities for further experimental investigation of the role of nuclear lamina in chromatin loop tethering.

### Tail-dependent switching of H2A.Z nucleosomes between the stable vs. unstable state, and between peripheral vs. scattered topography

The H2A.Z-containing nucleosomes detected by ZAbA are unusually stable in the sense that H2A.Z is evicted from the nucleosomes at higher salt concentrations than typical for H2A, when studied by QINESIn *in situ*. This was reproduced employing *in vitro* assembled chromatin (Fig. 1G), in line with previous studies where chicken erythrocyte chromatin was subjected to hydroxyapatite chromatography ^38^, suggesting that this is an intrinsic feature of the reconstituted nucleosome array containing this variant, i.e. it can be observed in the absence of nucleosome-unrelated potential interacting partners. The fact that the C9 peptide efficiently modulates the stability of H2A.Z-containing nucleosomes suggests that molecular interactions involving the tail rather than its influence on H2A.Z conformation, determine stability.

Nuclear architecture, when portrayed by ZAbA- or ZAbB-detected H2A.Z and H3K9me3, could also be modulated via addition of the C9 peptide to nuclei or to live cells (Fig. 3B-C, Fig. 5B, C, Fig. 6A, B; Suppl. Figs. 13C, E, Suppl. Fig. 21E, F), suggesting that the topographic differences in the nuclei containing the full-length or ΔC forms are the direct result of altered structure and functioning of the histone variant, rather than secondary, e.g. compensatory, consequences thereof developing in the cells expressing the truncated variant. To what extent differential masking/unmasking or changed accessibility of ZAbA- or ZAbB-binding epitopes in the different nuclear compartments may contribute to the altered nuclear landscape, have not been quantitatively determined.

Based on recent cryo-EM analyses ^45^, a H2A motif coined “regulating-octamer-folding” (ROF) plays an important role in nucleosome stability, interacting with the N-terminal H3 helix. This motif, residing between amino acids 106–111, is present both in H2A and H2A.Z.1. In the alternatively spliced variant H2A Z2.2, it is replaced by a noncanonical ROF and the terminal 14 amino acids are also missing as compared with H2A.Z.1. The destabilized nature of H2A.Z.2.2 containing nucleosomes, not investigated in our work, was attributed primarily to the disruptive effect of the noncanonical ROF in ^45^. The contribution to stability of the last 14 amino acids missing from H2A.Z.1 was not possible to judge, since this segment (encompassing the noncanonical ROF) was invisible in cryo-EM. Based on the efficient modulation of nucleosome stability with added C9, the contribution of the C-terminal tail to overall nucleosome stability may be very significant. According to another recent cryo-EM study ^21^, the H2A.Z-containing mononucleosomes formed on the Widom 601 sequence are less stable than those containing canonical H2A, a feature determined by the last 6 amino acids of its unstructured C-terminal tail, while the variant histone-containing nucleosome arrays assume a more regular and condensed state. Whether interactions between the enlarged acidic patch of the H2A.Z nucleosome and the H4 tail of the neighboring nucleosome described previously, or interactions involving the C-terminal tail, or both explain the more condensed state of the H2A.Z-containing chromatin fibers, have not been determined.

In view of the data presented, the tail may mediate interactions between neighboring chromatin fibers, increasing the stability of the nucleosomes connected. When this engagement is thwarted by reader protein binding, tail truncation or masking by a tag, the canonical stability features prevail. Tethering of peripheral heterochromatin to the lamina, perhaps involving H2A.Z^lmn^, could also account for the tail-dependence of increased stability of the nucleosomes involved. As a third alternative, the C-terminal tail could engage with a binding site within the same nucleosome rendering the variant histone more resistant to salt-induced dissociation; in this case the intranuclear localization and stability of H2A.Z-nucleosomes would not be directly linked. For a further, more complex scenario, see Suppl. Discussion.

The salt-sensitivity of the ZAbB-detected or tagged H2A.Z-containing nucleosomes were similar to each other and also to that of the canonical H2A-containing nucleosomes, what is at variance with the more open structure of the H2A.Z mononucleosomes in the cryoEM study ^21^. This may be because the degree of freedom of DNA termini in nucleosomes reconstructed with short pieces of DNA containing only the Widom sequence could be higher than what would be measured in the case of nucleosomes complete with longer linker DNA segments, in line with ref. ^56^. It is also possible that H2A.Z can stabilize the histone octamer in a tail-dependent fashion as measured in our assay, while simultaneously weakening the histone-DNA interactions at the DNA entry and exit points as detected by cryo-EM. However, recent AFM studies suggest that the N-terminal part of the histone, not the C-terminal, controls nucleosome-DNA binding strength ^34^.

### The stability of H2A.Z nucleosomes appear not to be directly affected by isotype specific features, PTMs and binding of PWWP2A

PTMs targeting the C-terminal region of H2A.Z ^57^ were considered as potential determinants of nucleosomal stability features. However, at least certain PTMs appear not to play a primary role in the case of H2A.Z containing nucleosomes (Suppl. Fig. 2I-L). This conclusion applies also to acetylation since we did not see any significant shift in the affinity of H2A.Z binding to nucleosomes when the global extent of histone acetylation was increased by treatment with HDAC inhibitors or when a non-acetylatable H2A.Z mutant was expressed in the cells (Suppl. Fig. 2I and L). The observation that the antibody supposed to be specific for acetylated H2A.Z ^58^ behaves as ZAbA (Suppl. Fig. 2L) is due to the fact that it readily recognizes the non-acetylated H2A.Z of the 5KR cells devoid of N-terminal acetylations; the H2A.Z-specificity of this antibody was verified in the silencing experiment shown in panel of Suppl. Fig. 5E.

Reader proteins such as PWWP2A ^59^, as well as chaperones/remodelers ^46, 59^ can bind to the H2A.Z C-terminus, raising the possibility that binding of these factors to H2A.Z can modulate the stability of H2A.Z containing nucleosomes. PWWP2A apparently does not contribute to the unusually high stability of the ZAbA detected H2A.Z-nucleosomes in our assay conditions, since it dissociates from the chromatin already at low salt (Suppl. Fig. 3A-C). *In vivo*, however, binding of this or any of the other readers may prevent the C-terminal tail from engaging with a neighboring nucleosome.

According to the H2A-like elution profiles of fluorescent protein-tagged H2A.Z.1 and Z2 (Suppl. Fig. 4A), the two tagged hypervariants may not differ significantly in their stability features at euchromatic locations. However, given that the elution curves, reflecting the nucleus-averaged stability of all H2A.Z species present, depend on even small tags, especially on their C-terminus (Suppl. Fig. 4F), it becomes difficult to make a comparison based on data obtained with tagged histones. Similar observations on the influence of tags have been reported on the Flag tag-dependent molecular interactions involving H2B ^60^. Thus, potential differences in the stability features of H2A.Z.1 and H2A.Z.2.1 could not be assessed based on the elution characteristics of the tagged hypervariants. However, expression of either hypervariant on a DKO background made such a comparison possible (Suppl. Fig. 2A), and have yielded negative results. In summary, the above negative data are in line with the finding that *in vitro* reconstituted nucleosomes are also stable (Fig. 1G) in line with ref. ^38^. This observation is in line with a model where the higher stability is the primary feature of H2A.Z-nucleosomes when assessed *in situ*, unless or until the inter- or intranucleosomal links mediated by the C-terminal tail are overridden by competing molecular associations.

### The destabilized state of the H2A.Z mononucleosomes and their aggregation induced stabilization may be synergistic

Nucleosome stability is assessed routinely by exposing them to challenging conditions like elevated salt concentration, which may itself perturb the structure. However, we observed entirely different elution characteristics for ZAbA- and ZAbB-detected, and tagged histone containing nucleosomes using the same salt concentration series; furthermore, salt resistance of the nucleosomes recognized by ZAbA was reverted upon nicking treatment. Thus, the stability features observed in the elution experiments are just revealed by, and are not the consequence of, the salt treatment. On the other hand, the nucleosomal binding of the labeled C9 peptide was enhanced to ∼90 nM in the presence of 700 mM salt, while DNA binding was unaffected (Fig. 4); this ionic strength favors loss of the histone dimers according to high precision FRET studies ^51^. A more open, nucleosomal structure, experimentally induced in the canonical nucleosomes by salt, could accommodate the C-terminal tail of a juxtaposed H2A.Z nucleosome. The tail of the histone variant is likely to protrude out of the nucleosome so as to be able to reach a binding site in a neighboring chromatin fiber ^61, 62^. The relaxed state of plasmid DNA in the reconstituted nucleosomal arrays of our hydroxyapatite experiments (see Materials and Methods; Fig. 1G) could also allow such interactions to occur between nucleosomes residing on the same plasmid and explain increased salt-resistance. On the other hand, an interpretation based on the possible interaction of the tail with a binding site formed within the same nucleosome with consequential strengthening of molecular cohesion and stability would also be compatible with our salt elution data. In this latter scenario, the increased stability of H2A.Z nucleosome arrays would be unrelated to the H2A.Z tail and could involve the acidic patch, as suggested ^21^.

As Suppl. Fig. 18 demonstrates, about 12 % of the peptide was found in the slower fraction representing C9-CF bound to nucleosomes prepared from recombinant histones and the 147 bp long Widom 601 core sequence ^63^, while the slow component was undetected in the case of the labeled scrambled peptide. Based on this value, the K_d_ of C9 binding to these nucleosomes is ∼320 nM, (calculated as in ^50^). Weaker association (K_d_ ∼2 µM) was obtained for nucleosomes containing longer (170 bp) DNA (Fig. 4). K_d_ values in the range of 1 nM to ≥ 1 μM measured for the binding of various H3K4-modified peptides to purified nucleosomes have been considered physiologically relevant ^64^. This relatively weak binding, as measured at low salt conditions, may be sufficient to support the formation of a condensed superstructure from juxtaposed nucleosomes; however, it could also be antagonized *in vivo* by structural components of the chromatin like H1 and HMGB1, with K_d_ values for DNA binding in the low micromolar range ^65^, washed out from the nuclei in the elution assay conditions. At low salt, the affinity of the tail to nucleosomes and DNA were also similar (Fig. 4), suggesting that it is the internucleosomal linker DNA that primarily engages with the tail in these conditions. Consolidation of the interaction to K_d_=90 nM at 700 mM salt was not observed in the case of pure DNA, so C9-CF is suggested to bind to the nucleosome core in these ionic strength conditions.

C9-CF-binding to H2A.Z-nucleosomes exceeded its binding to canonical nucleosomes at low ionic strength (Suppl. Fig. 18). It remains to be investigated how elevated salt concentrations influence this difference, to what extent C9 binds to the nucleosome core and to the linker DNA, how this binding depends on the length of the DNA containing the nucleosome positioning sequence and on the species origin of the histones. With these limitations, our FCS measurements support the possibility that binding of the H2A.Z C-terminal tail to nucleosomal components within the same nucleosome or mediating nucleosome-nucleosome contacts is a key factor in determining nucleosome stability and establish the experimental platform for further biophysical studies.

### Chromatin accessibility features are affected by interactions involving the H2A.Z tail

The parallel sensitization of chromatin to nickase, MNase and DNAse I in H2A.ZΔC-expressor DKO DT40 nuclei (Fig. 2E-G, Suppl. Fig. 13A) suggests that access to DNA rather than the exonucleolytic digestion subsequent to the initial nicking is the rate-limiting step in MNase digestion. The relaxation of internucleosomal superhelicity upon the initial single-strand cleavages significantly contributes to nucleosome destabilization ^35, 66^, rendering the entire chromatin loop more open to further cleavages, amplifying and extending the effect of the treatment.

The remarkable differences in overall nuclease sensitivity between the full-length H2A.Z.1-expressor and H2A.ZΔC-expressor DKO DT40 nuclei (Fig. 2) may reflect the altered contribution to what is measured over the entire nucleus by the dispersed peripheral heterochromatin, and may also reflect alterations in the dynamics of the euchromatic, e.g. of the promoter proximal H2A.Z; indeed, these nucleosomes exhibit remarkable heterogeneity in terms of accessibility features of the DNA packaged ^22^. The latter nucleosomes comprise a relatively minor fraction of the total chromatin, and even spreading of destabilization upon the initial nicking potentially to the whole loop ^66^ could not account for the increment of MNase sensitivity detected on a global level in ΔC vs. full-length H2A.Z.1-expressor DT40 DKO cells, or upon C9 treatment of HeLa nuclei (Fig. 3). Thus, further DNA regions must have also been affected by the C9 treatment. Indeed, comparison of the read coverages before and after MNase treatment revealed that the changes in total coverage were reflected by similar changes at certain repetitive elements (Suppl. Fig. 17C).

C9 treatment of nuclei at low ionic strength did not lead to eviction of either canonical or H2A.Z-containing nucleosomes (Suppl. Fig. 16A, B). Thus, the C9-induced increment in accessibility of a large fraction of genomic DNA to nucleases was not the experimental result of competition between the peptide and nucleosomes for DNA binding. We speculate that accessibility in peripheral heterochromatin is dependent upon chromatin condensation and/or laminar tethering mediated by the C-terminal tail of the histone variant. Loss of laminar anchorage or change in the condensation state may alter internucleosomal superhelicity what could lead to nucleosome destabilization involving the entire chomatin loop ^66^. The C9 treatment-enhanced release of a number of proteins of its euchromatic interactome by MNase (Suppl. Table 4) reveals that accessibility of DNA flanking H2A.Z-nucleosomes binding these molecular complexes is tail-dependent also in euchromatin. These DNA regions, however, may comprise only a small fraction of the total DNA content, not accounting for the overall increments in digestability by various nucleases.

### H2A.Z tail-dependence of gene expression and the possibility of epigenetic modulation

In the DT40 experimental system, the differences in gene expression were not extensive, especially in the case of ΔC vs. DKO H2A.Z.1 (Suppl. Fig. 14, Suppl. Table 3), perhaps due to the adaptation of the cells to their tailless histone variant, in comparison with the global contribution of H2A.Z to gene expression regulation ^36^, and with the massive differences in chromatin structure both in terms of H2A.Z nuclear localization and accessibility features (Figs. 2 and 3). Nonetheless, these differences have functional consequences as shown by the altered DNA damage response (DDR; Suppl. Fig. 15). The architectural changes exhibited by DKO H2A.ZΔC chromatin relative to full-length histone expressing cells, and the increased sensitivity of these cells to etoposide and doxorubicin (Suppl. Fig. 15 are closely reminiscent of the dramatic influence of H2A.Z on resistance to genotoxic stress in Saccharomyces cerevisiae ^67^. For further possible connections of the gene expression changes to DDR, see Supplementary Discussion.

The C9 peptide could be efficiently introduced into live HeLa and melanoma cells. Changes in gene expression could be primarily accounted for by modulation of euchromatin. On the other hand, alterations of constitutive, mostly peripheral heterochromatin, which comprises up to 40 % of the human genome, could be relevant in the context of aging, various neurological disorders, and carcinogenesis ^68^. The rearrangement of peripheral heterochromatin upon treatment of nuclei as well as of live cells with C9 demonstrates that global features of heterochromatin are amenable to modulation by the peptide. The CUT&RUN/MS data showing that the linker and/or nucleosome-free DNA around the H2A.Z containing nucleosomes become readily accessible upon C9 treatment (Suppl. Table 4) directly demonstrate that euchromatic molecular complexes of H2A.Z (likely in their inactive state) can also be readily modulated by the peptide. Indeed, introduction of the peptide into live cells leads to down-regulation of a cohort of genes (Fig. 6C), in line with the large number of genes found to be H2A.Z-regulated (ref. 127 in ^33^). Certain pathways are overrepresented among the down-regulated genes (Fig. 6L), so the effect cannot be due to nonspecific transcriptional suppression or cytotoxicity, also in view of the fact that the continuously growing cells showed no loss of viability (Suppl. Fig. 20). Intriguingly, the down-regulated genes themselves are exclusively localized outside the Giemsa-dark bands (Fig. 6E) containing late-replicating DNA ^69^. Thus, although H2A.Z is deposited all over the chromosomes (Fig. 6H), expression-determining interactions of its C-terminal tail appear to be limited to genes residing outside the late-replicating domains, raising the possibility of an intriguing connection between activation of early replication origins by H2A.Z ^70^ and tail-dependent transcriptional regulation involving the variant histone. For an assessment of the possible changes elicited by C9 treatment in the expression of repetitive elements see Supplementary Methods and Supplementary Discussion.)

In summary, the H2A.Z tail impacts chromatin at each level of its organization, from nucleosome to chromosome bands, and the enigmatic involvement of H2A.Z in both gene activating and silencing processes appears to be determined to a great part by the molecular interactions involving its unstructured C-terminal tail. We propose that the tail may be bound to the nucleosome core or internucleosomal DNA, increasing nucleosome stability via engagement of the tail within the same nucleosome or by promoting chromatin condensation, or alternatively, to reader proteins/chaperones/remodelers, preventing this engagement. A high salt-resistant fraction of H2A.Z was detected in the nuclear lamina, which may play a role in the tethering of heterochromatin. The alternative engagements of the tail affect the stability and landscape of H2A.Z-nucleosomes, chromatin accessibility features in both euchromatin and heterochromatin, with marked consequences on gene expression. Further investigation of this remarkable example of epigenetic modulation is warranted also in the hope that the C9 effect may be exploited to experimentally modulate different cellular states under H2A.Z control.

## MATERIALS AND METHODS

### Chemicals, peptides

All reagents were from Sigma-Aldrich (St. Louis, Missouri, USA) unless otherwise stated. Fmoc-Rink Amide MBHA resin and all amino acid derivatives used in this study were purchased from Iris Biotech GmbH (Marktredwitz, Germany). Peptides of different lengths, corresponding to the C-terminus of H2A.Z were synthesized manually on 2CTC resin (0.6 mmol/g loading capacity) following standard Fmoc/*^t^*Bu strategy using the *in situ* DIC/HOBt coupling method. The complete sequences were removed from the resin with 10 ml of cleavage mixture (95% TFA, 2.5% triisopropylsilane (TIS), 2.5% H_2_O), stirred for 1.5 h at RT. The crude products were precipitated in ice cold MTBE (methyl-tert-butyl ether) and centrifuged for 4 min at 4000 rpm. After washing three times with MTBE, the crude products were dissolved in water and the solution was purified by RP-HPLC and lyophilized.

The fluorescent peptide derivative of C9 was prepared similarly but was elongated on the C-terminus with an aminohexane (Ahx) group and a lysine. The peptide was labeled on resin, *via* coupling the lysine’s LJ-amino group with 5(6)carboxy-fluorescein (CF), which resulted the peptide H-GKKGQQKTV-Ahx-K(CF)-OH. ‘Scrambled’ reference peptide, used as a negative control, was generated by random shuffling of amino acids. The generated sequence was checked against Uniprot KB database. This was repeated until a peptide sequence not showing match with the database was found. The resulting control peptide was: H-KQGTGKVQK-OH.

### Plasmids

The plasmids carrying different isoforms of the histone variant H2A.Z (provided by Juan Ausio) were transformed into *Escherichia coli* DH5α by heat shock and selected on LB plates containing 100 μg/ml kanamycin.

### Cells

HeLa, H2B-GFP expressor HeLa, H3-GFP expressor HeLa ^71^(provided by Hiroshi Kimura, Yokohama, Japan), GFP-H2A.Z expressor HeLa (provided by H. T. Marc Timmers), 293T and MEL1617 cells (from Coriell Institute for Medical Research) were cultured in DMEM supplemented with 10% FCS, 2 mM L-glutamine, 100 μg/ml streptomycin, 100 U/ml penicillin. WM35 cells (Coriell Institute for Medical Research) were cultured in RPMI supplemented with 10% FCS, 2 mM L-glutamine, 100 μg/ml streptomycin, 100 U/ml penicillin. Wild type, H2A.Z.1 knock-out, H2A.Z.2.1 knock-out and mutant (5KR and ΔC) H2A.Z.1 expressing double knock-out DT-40 chicken B cells (provided by Masahiko Harata ^36, 37^) were cultured in DMEM supplemented with 2% chicken serum, 8% FCS, 2 mM l-glutamine, 100 μg/ml streptomycin, 100 U/ml penicillin.

### Embedding live cells into low melting point agarose

Prior to embedding, 8-well microscope chambers (Ibidi, Martinsried, Germany) were coated with 1% (m/v) low melting point (LMP) agarose. 150 μl liquid agarose diluted in distilled water was dispensed into each well and was immediately removed so that a thin agarose layer remained on the surfaces and was left to polymerize on ice for 2 minutes, then kept at 37°C until the surface of the wells dried out. This coating procedure was repeated once more on the same chambers. Embedding was performed keeping cells and agarose at 37°C. The cell suspension containing 6×10^6^ cells/ml was mixed with 1% LMP agarose diluted in PBS at a v/v ratio of 1:3. 22 μl of the cell-agarose suspension was dispensed in the middle of the wells and the chambers were covered with home-made rectangular plastic coverslips cut out from a 200 μm thick, medium weight polyvinyl chloride binding cover (Fellowes, Inc., Itasca, Illinois, USA). Cells were left to sediment on the surface of the coated wells for 4 minutes at 37°C, then kept on ice for 2 minutes. After polymerization of the agarose, 300 μl ice cold complete culture medium was added to each well, a step aiding removal of the coverslips.

### Preparation of nuclei/permeabilization, histone eviction by salt, nuclear halos

The agarose embedded cells at the bottom of the wells were washed with 500 μl ice cold PBS, 3 times for 3 minutes each, then permeabilized with 500 μl ice cold 1% (v/v) Triton X-100 dissolved in PBS/EDTA (5 mM EDTA in PBS), for 10 minutes. This step was repeated once more. After permeabilization, nuclei were washed with 500 μl ice cold PBS/EDTA 3 times for 3 minutes and were treated with different concentrations of NaCl or intercalator solutions on ice. Nuclei were incubated with 500 μl of ice cold salt or intercalator solution for 60 minutes. At this point nuclear halo samples were prepared by adding 2M NaCl dissolved in PBS/EDTA to the nuclei for 60 minutes on ice (neutral halos). After this treatment, nuclei were washed with 500 μl ice cold PBS/EDTA 3 times for 3 minutes. Since NaCl was diluted in PBS/EDTA, the salt concentrations indicated on the X axes of the graphs in all the figures show the total NaCl concentrations together with NaCl present in the PBS buffer. For the analyzis of the curves SigmaPlot 12.0 software was applied, using either the ‘Sigmoid 3 parameter’ (in the case of linear plots) or ‘Standard curves: Four Parameter Logistic Curve’ (in the case of logarithmic plots) curve-fitting subroutines. Elution curves were ’normalized to 0’ by substracting the smallest fluorescence intensity value from all the others, unless stated otherwise, and to ’1’ by dividing the mean fluorescence intensities represented by the data points by that of the non-treated sample. The number of analyzed G1 nuclei was between 200-2000/well, out of the about 500-5000 cells scanned. All the SEM values indicated in the figures were calculated from the datapoints of the cell population analyzed in the given experiment.

### Peptide treatment of permeabilized nuclei

The synthetic peptides were used at a final concentration of 30 μM, dissolved in PBS/EDTA/1%BSA. 200 μl of peptide solution/well was applied and nuclei were incubated with the peptide overnight at 4°C. After incubation, the peptide solution was washed out with 500 μl ice cold PBS/EDTA 3 times.

### Introduction of peptides into live cells

0.5×10^6^ HeLa or MEL1617 melanoma cells were placed in a 35 mm cell culture Petri dish and grown overnight before peptide treatment. Cyclodextrin/peptide complex formation was performed by mixing 30 μM peptide and 300 μM SBECD (Sulfobutylether-β-Cyclodextrin; CycloLab, Budapest, Hungary) diluted in colorless, serum free RPMI1640 and incubated for 1 h at room temperature. 2 ml SBECD/peptide complex was added to the cells and incubated for 2h at 37°C in 5% CO_2_ atmosphere. After incubation, cells were washed with complete RPMI medium once and further cultured overnight before agarose embedding.

### Immunofluorescence labeling

After salt or intercalator treatment the samples were incubated with 500 μl 5% (m/v) Blotto Non-Fat Dry Milk (Santa Cruz Biotechnology Inc., Santa Cruz, California, USA) in PBS/EDTA for 30 minutes on ice, to decrease nonspecific binding of the antibodies. The blocking solution was washed out with 500 μl ice cold PBS/EDTA 3 times for 3 minutes and indirect immunofluorescence labeling was performed using rabbit polyclonal anti-H2A.Z (ab97966 (ZAbA), ab4174, Abcam, Cambridge, UK; 1 mg/ml), sheep polyclonal anti-H2A.Z (acetyl K4+K7+K11, ab18262, Abcam, Cambridge, UK; 0.5 mg/ml), rabbit polyclonal anti-H2A.Z (PA5-17336 (ZAbB), Thermo Fisher Scientific, Waltham, Massachusetts, USA; 62 µg/ml), rabbit polyclonal anti-H2A.Z (07-594, Merck-Millipore, Darmstadt, Germany), rabbit polyclonal anti-PWWP2A (NBP2-13833, Novus Biologicals, Centennial, Colorado, USA; 0.2 mg/ml), rabbit polyclonal anti-H2A.X (ab11175, Abcam, Cambridge, UK; 1 mg/ml), rabbit polyclonal anti-H2A (ab18255, Abcam, Cambridge, UK; 1 mg/ml), rabbit polyclonal anti-H2B (ab52484, Abcam, Cambridge, UK; 1 mg/ml), mouse monoclonal anti-H1 (ab71591, Abcam, Cambridge, UK; 1 mg/ml), mouse monoclonal anti-HP1 (ab234085, Abcam, Cambridge, UK; 1 mg/ml), mouse monoclonal anti-γH2A.X (05-636, Merck-Millipore, Darmstadt, Germany), mouse monoclonal anti-H3K4me3 (^72^; 0,5 mg/ml), mouse monoclonal anti-H3K9me3 (^72^; 0,5 mg/ml) or mouse monoclonal anti-H3K27me3 (^73^; 0,5 mg/ml) primary antibodies, all diluted in 150 μl of PBS/EDTA/1% BSA (PBS/EDTA supplemented with 1% w/v bovine serum albumin), at 4°C, overnight. All the above antibodies were applied to the wells at a titer of 1:800. After labeling with the primary antibodies, the nuclei were washed with 500 μl ice cold PBS/EDTA 3 times for 10 minutes. Labeling with the secondary antibodies was performed in 150 μl PBS/EDTA for two hours on ice, using Alexa fluor 488 (A488) or Alexa fluor 647 (A647) conjugated goat anti-mouse IgG or goat anti-rabbit IgG antibodies (Thermo Fisher Scientific, Waltham, Massachusetts, USA; 2 mg/ml). In the case of sheep polyclonal anti-H2A.Z (acetyl K4+K7+K11, ab18262) Alexa fluor 633 (A633) conjugated goat anti-sheep IgG (Thermo Fisher Scientific, Waltham, Massachusetts, USA; 2 mg/ml) secondary antibody was used. The secondary antibodies were also used at a titer of 1:800, diluted in PBS/EDTA from 2 mg/ml stock solutions. After labeling with the secondary antibodies the agarose embedded nuclei were washed with 500 μl ice cold PBS/EDTA 3 times, for 10 minutes. Then the samples were fixed in 1% formaldehyde (dissolved in PBS/EDTA) at 4°C, overnight. After fixation the wells containing the embedded nuclei were washed with 500 μl ice cold PBS/EDTA 3 times for 3 minutes and were stained with 200 μl of 12,5 μg/ml propidium–iodide (PI, dissolved in PBS/EDTA) for 30 minutes, on ice. The stained nuclei were washed 3 times with 500 μl ice cold PBS/EDTA for 3 minutes. Fluorescence intensity distributions were recorded using an iCys laser scanning cytometer (LSC), as described below.

### Western blot

200 - 200 ng H2A.Z.1, H2A.Z.2, H2A (H2042, Sigma-Aldrich, St. Louis, Missouri, USA; 1 mg/ml) and H2B (H2167, Sigma-Aldrich, St. Louis, Missouri, USA; 1 mg/ml) histones were separated on a 12 % SDS–polyacrylamide gel and electro-blotted onto a 0.45 μm pore size PVDF membrane. The blot was saturated with milk blocking buffer (5% milk power / 0.2% Tween-20 / PBS) for 1 hour and then labeled by ZAbA or ZAbB. Both were applied at a final concentration of 0.4 μg/ml in 1:2500 dilution in milk blocking buffer overnight at 4°C. After washing with 0.2% Tween-20/PBS for 6 × 5 minutes, the membrane was incubated with goat-anti-mouse IgG secondary antibody conjugated with horseradish peroxidase (ab6789, Abcam, Cambridge, UK; 2 mg/ml) at a final concentration of 0.8 μg/ml / in 1:2500 dilution in milk blocking buffer for 1.5 hours at room temperature. After washing with 0.2% Tween-20/PBS for 6 × 5 minutes, the bands were visualized with SuperSignal™ West Pico PLUS Chemiluminescent Substrate (Thermo Fisher Scientific, Waltham, MA). The signal was detected by chemiluminescence using the FluorChem Q gel documentation system (Alpha Innotech Corp., San Leandro, CA).

### Etoposide treatment

Agarose embedded live cells were treated with etoposide (TEVA, Debrecen, Hungary) used at a final concentration of 25 µM. The drug was diluted in 300 µl complete DMEM medium and the cells were incubated together with the drug at 37°C in 5% CO_2_ atmosphere.

### Laser Scanning Cytometry (LSC)

Automated microscopic imaging was performed using an iCys instrument (iCys® Research Imaging Cytometer; CompuCyte, Westwood, Massachusetts, USA). Green fluorescent protein (GFP), SYBR Gold, A488 and PI were excited using a 488 nm Argon ion laser, A647 with a 633 nm HeNe laser. Fluorescence signals were collected via an UPlan FI 20× (NA 0.5) objective, scanning each field with a step size of 1.5 µm. GFP and A488 were detected through 510/21 nm and 530/30 nm filters, respectively, while A647 and PI were detected through a 650/LP nm filter.

Data evaluation and hardware control were performed with the iCys 7.0 software for Windows XP. Gating of G1 phase cells was according to the fluorescence intensity distribution of the DNA labeled with PI or SYBR Gold.

### Confocal Laser Scanning Microscopy (CLSM)

Confocal images were taken using an FLUOVIEW FV 1000 confocal microscope (Olympus, Center Valley, Pennsylvania, USA) based on an inverted IX-81 stand with an UPLS APO 60× (NA 1.35) oil immersion objective. GFP or A488 were excited by a 488 nm Argon ion laser. A647 and PI were excited by a 633 nm and 543 nm HeNe laser, respectively.

### Colocalization and texture analyzis

For image and texture analyzes, the Just Another Colocalization Plugin of the Image J software (http://imagej.nih.gov/ij/) and the MeasureGranularity module of the CellProfiler 2.2.0 software, respectively, were used. For colocalization measurements Manders colocalization coefficient (MCC) representing the fraction of overlapping pixels was calculated in Image J.

### MNase, Nickase and DNase I treatment

Live cells were embedded into agarose as described above and treated with 500 μl ice cold lysis buffer (0.4% (v/v) Triton X-100, 300 mM NaCl, 1 mM EDTA, 10 mM Tris-HCl, pH 8.0) for 10 minutes, followed by treatment with 500 μl ice cold 1% (v/v) Triton X-100 dissolved in PBS/EDTA, for 10 minutes, then washed 3 times with 500 μl ice cold PBS/EDTA. MNase, the frequent cutter Nt.CviPII nickase (recognition site: CCD; New England Biolabs Inc., Ipswich, Massachusetts, USA) and DNase I were applied after the washing steps following permeabilization. Where indicated, the permeabilized and washed nuclei were incubated with the peptides (C9 or SCR) overnight at 4 °C, and the peptides were washed out before nuclease addition. Before digestion, the samples were equilibrated with MNase buffer (1 mM CaCl_2_, 50 mM Tris-HCL pH 7.5), nickase buffer (10 mM Tris-HCl pH 8.0, 50 mM NaCl, 10 mM MgCl_2_, 1 mg/ml BSA) or with DNase I buffer (10 mM Tris-HCl pH 8.0, 0.1 mM CaCl_2_, 2.5 mM MgCl_2_) by washing 3 times with 500 µl of the buffer solutions. Nickase treatment was performed in 300 µl nickase buffer for 30 min at 37 °C, using the enzyme at a final concentration of 0.5 U/ml. MNase and DNase I digestion were performed in 300 µl MNase, or DNase I buffer for 10 min at 37 °C, at a final concentrations indicated on the figures. After enzymatic treatments, the samples were washed with 500 μl ice cold PBS/EDTA 3 times for 3 minutes.

### Microarray experiments

For microarray analyses, Cyanine3 (Cy3) labeled cRNA was prepared from 0.1 micro-g Total RNA using the Low Input Quick Amp Labeling Kit (Agilent Technologies) according to the manufacturer’s instructions, followed by RNeasy column purification (QIAGEN). For methodical details of the Agilent platform used, see Supplementary Methods.

### RNA-Seq and pathway analyzis

To obtain global transcriptome data high throughput mRNA sequencing analysis was performed on the Illumina sequencing platform. MEL1617 cells were grown in 6-well plates and treated with C9 or scrambled-C9 peptid for 2 hours (see details above). Total RNA was isolated from 1 million cells in triplicates using GenElute Mammalian Total RNA Miniprep Kit (Sigma) according to the manufacturer’s instructions. RNA sample quality was checked on Agilent BioAnalyzer using Eukaryotic Total RNA Nano Kit according to manufacturer’s protocol. Samples with RNA integrity number (RIN) value >7 were accepted for the library preparation process. For details of the preparation of RNA-Seq libraries, amplification and sequencing on an Illumina platform, see Supplementary Methods. For the steps of further sample processing and bioinformatic analyses of the fastq data, see Supplementary Methods.

### Whole genome sequencing for comparative mapping of MNase sensitivity before and after C9 treatment

HeLa cells were embedded into agarose in the wells of 8-well Ibidi chambers and permeabilized as described above. Different chambers were prepared parallel for LSC measurement (to measure the effect of the peptide on the overall degree of MNase digestion) and for whole genome sequencing. After peptide treatment, nuclei were digested with 10^−3^ U/ml MNase as described above. After enzymatic treatment, nuclei were washed with 500 μl ice cold 1×PBS/EDTA 3 times for 3 minutes, followed by fixation with 4% formaldehyde for 10 min, on ice. After fixation, samples were washed with 500 μl ice cold 1×PBS/EDTA 3 times, for 3 minutes each. For LSC measurement, the samples were stained with 200 μl 1×SYBRGold diluted in PBS/EDTA, on ice for 30 min, then washed with 500 μl ice cold 1×PBS/EDTA 3 times before LSC scanning. For whole genome sequencing, agarose embedded cells were scraped and removed from the wells. 12 wells of CTRL, CTR(MN), C9 or C9(MN) were pooled resulting in 400 μl samples. These were melted up to 70 °C for 10 min, then cooled down to 42 °C and digested with 4 U of agarase enzyme/400 μl sample (Thermo Fisher Scientific, Waltham, Massachusetts, USA) for 30 min. Samples were further digested with 40 μg/400 μl proteinase K (Thermo Fisher Scientific, Waltham, Massachusetts, USA) overnight at 55 °C. After digestion, DNA was sheared by sonication using a Diagenode Bioruptor instrument (Diagenode, Liège, Belgium) to generate ≈250 bp fragments, that were purified (MinElute PCR Purification Kit; Qiagen Inc., Valencia, California, USA) and stored at −70 °C until sequencing.

After shearing, size selection was performed to enrich for ∼280 bp DNA fragments using DNA Clean Beads (MGI, Shenzen, China). The sequencing library was generated using MGIEasy Universal DNA Library Prep Set (MGI) according to the manufacturer’s protocol. Briefly, the size selected DNA fragments went through end repair, A-tailing, adaptor ligation, and amplification steps. The double-stranded library fragments were circularized and one of the strands was digested. Circularized single stranded fragments were used for generating DNA nano balls (DNB) by rolling circle amplification. DNB sequencing was performed on a MGI DNBSEQ GS-400 instrument (MGI), to yield paired-end, ∼150 bp long sequencing reads. Library preparation, sequencing and primary data analysis were performed at the Genomic Medicine and Bioinformatics Core Facility of the University of Debrecen, Hungary.

Analyses of the Whole Genome Sequencing data was performed on the Galaxy platform (EU server: usegalaxy.eu) ^74^. For details of quality control, mapping of the reads to the reference genome, calculations related to read coverages, specification of the peak-sets download from ENCODE, see Supplementary Methods.

### Sample preparation for mass spectrometric (MS) measurements of the salt resistant fraction of nuclear proteins

80×10^6^ H3-GFP expressing HeLa cells in 2.5 ml cell culture medium were mixed with an equal amount of 1% LMP diluted in PBS. Agarose blocks were prepared where each blocks contained 90 µl of agarose/cell suspension. The blocks were washed 3 times in 15 ml 1×PBS for 10 min. Permeabilization of cells was performed in ice cold 15 ml 1% Triton-X 100 diluted in PBS/EDTA, twice for 30 min. Blocks were washed 5 times in 15 ml PBS/EDTA. Blocks were treated with 15 ml EBr diluted in PBS/EDTA/600 mM NaCl at a concentration of 100 µg/ml, or with 15 ml 1.2 M NaCl solution diluted in PBS/EDTA, for 60 min. Blocks were washed 3 times in 15 ml PBS/EDTA for 20 min. and the proteins that remained in the nuclei were eluted with 15 ml of 2 M NaCl diluted in PBS/EDTA for 60 min. All the washing steps were performed using ice cold solutions. Eluted proteins were concentrated using a 10K Amicon tube (Merck-Millipore, Darmstadt, Germany) and the buffer was changed to PBS. Proteins were eluted from the filter with 250 µl PBS and stored at −20°C for MS analyzes. For details of sample processing, Orbitrap MS analyses and protein identification, see Supplementary Methods.

### LC-MS/MS analyses of salt-eluted nuclear samples

The samples were digested in-solution with trypsin. Before digestion, samples were reduced using 10 mM dithiothreitol for one hour at 56°C, followed by alkylation with 20 mM iodoacetamide for 45 min. Overnight trypsin digestion was carried out using stabilized MS grade bovine trypsin (Sciex, Framingham, MA, USA) at 37 °C. The reaction was stopped by adding concentrated formic acid and the digests were desalted using C18 Pierce tips (ThermoScientific, Waltham, MA, USA) according to the manufacturer’s protocol. The desalted peptides were dried and re-dissolved in 10 μl 1% formic acid. Before MS analysis the peptides were separated using a 180 min water-acetonitrile gradient at 300 nl/min flow rate on an Easy nLC1200 UPLC (ThermoScientific, Waltham, MA, USA) and the eluted peptides were examined on an Orbitrap Fusion mass spectrometer (ThermoScientific, Waltham, MA, USA). Data-dependent MS/MS scans of the 14 most abundant ions were recorded (Orbitrap analyzer resolution: 60,000, AGC target: 4.0e5, collision-induced dissociation fragmentation in linear ion trap with 35% normalized collision energy, AGC target: 2.0e3, dynamic exclusion 45 s). Protein identification was done with MaxQuant 1.6.2.10 software ^75^ searching against the Human SwissProt database (release: 2020.02, 20,394 sequence entries). Cys carbamidomethylation along with Met oxidation and N-terminal acetylation were set as variable modifications. Maximum two missed cleavage sites were allowed. Those proteins were considered, where at least two peptides identified with 1% FDR were present.

### Sample preparation for CUT&RUN MS measurements

HeLa cells suspended in cell culture medium were mixed with equal volume of 1% LMP agarose diluted in PBS. Agarose blocks were prepared where each blocks contained 90 µl of agarose/cell suspension with 2×10^6^ cells. 5 blocks were used for each sample (sample 1: no ZAbA + pAG/MNase activated, sample 2: ZAbA + pAG/MNase not activated, sample 3: ZAbA + pAG/MNase activated). Following ZAbA or ZAbB labeling, anti-rabbit IgG was added as a secondary antibody to improve yield, where indicated (see Table 4). Blocks were washed 3 times in 25 ml PBS for 10 min. Permeabilization of cells was performed in 25 ml ice cold lysis buffer (0.4% (v/v) Triton X-100, 300 mM NaCl, 1 mM EDTA, 10 mM Tris-HCl, pH 8.0) for 30 minutes, followed by treatment with 25 ml ice cold 1% (v/v) Triton X-100 dissolved in PBS/EDTA, for 30 minutes. Blocks were washed 4 times in 25 ml ice cold CUT&RUN Wash (designated CR Wash) /EDTA (20 mM HEPES, 150 mM NaCl, 0.5 mM spermidine and 5 mM EDTA) for 10 min. Using 5 blocks/sample, samples 2 and 3 were labeled with ZAbA at 4°C, overnight at a titer of 1:800 in 2.4 ml CR Wash/EDTA. After labeling, the blocks were washed 4 times in 10 ml ice cold CR Wash buffer (without EDTA) for 30 min and incubated with 10^−3^ U/ml pAG/MNase in 10 ml CR Wash buffer for 2 h on ice. After digestion, MNase was washed out by 10 ml ice cold CR Wash buffer 4 times for 30 min on ice. Chromatin bound MNase was activated by 5 ml ice cold CR Wash buffer supplemented with 2 mM CaCl_2_ for 30 min on ice. MNase digestion was stopped by adding 250 µl of 0.5 M EDTA and the blocks were further incubated at 37°C, for 30 min. Eluted proteins were concentrated using a 10K Amicon tube (Merck-Millipore, Darmstadt, Germany) and the buffer was changed to PBS. Proteins were eluted from the filter with 250 µl 1×PBS and stored at −20°C for MS analyzes. For details of sample processing, Orbitrap MS analyses and protein identification, see Supplementary Methods.

### Chromatin assembly and hydroxyapatite chromatography

Chromatin assembly reactions employing *Xenopus laevis* N1/N2-(H3, H4), mouse recombinant H2A.Z/H2B or H2A/H2B ^6^, 900ng pXbsF201 plasmid DNA and topoisomerase I were carried out, followed by supercoiling assays to ensure an equal number of nucleosomes between reconstitutes (a physiological spacing one nucleosome per 180 bp), as described previously ^76^. Assembled chromatin was applied to a hydroxyapatite column (Bio-Rad HTP) with 0.1 M NaCl, 0.1M KPO_4_, pH 6.7 as described ^38^, and washed successively with 0.1 M NaCl, 0.2 M NaCl and 0.4 NaCl in 0.1M KPO_4_, pH 6.7. Histones were eluted by a stepwise increase in NaCl concentration from 0.6-1.2 M in 0.1M KPO_4_, pH 6.7. The eluted fractions were loaded onto a 18% sodium dodecylsulfate-polyacrylamide gel and then stained with silver as described previously ^76^.

### Fluorescence correlation spectroscopy (FCS). Sample preparation

Mononucleosomes were reconstituted by slow salt dialysis from recombinant histones and Cy5 labeled 170 bp long DNA fragments centered around the well positioning Widom sequence according to ^77, 78^. The Cy5 label was used to determine the diffusion properties of the nucleosomes independently from the fluorescein labeled peptides. The quality of the reconstituted mononucleosomes was controlled by absorption spectroscopy revealing eventual aggregation and PAAgel electrophoresis allowing the quantification of unbound DNA in the sample. Only preparations without detectable aggregation and with less than 5 % of unbound DNA were used.

Commmercial mononucleosomes containing H2A or H2A.Z histones (see experiments shown in Suppl. Fig. 18) were purchased from Tebubio (France) (cat. no. 16-0009 and 16-1014, respectively).

FCS measurements were carried out in 50 to 100 µl volume on a sylanized Sensoplate plus 384 well plate (Greiner) at room temperature. 20 nM CF-labeled C9 peptide was either mixed with 100 nM reconstituted nucleosome or DNA or was measured alone in 5 mM or 700 mM NaCl TE (pH=7.5). The mixture was allowed to equilibrate to RT for 0.5 h before measurement.

FCS measurements were performed on a Nikon A1 Eclipse Ti2 confocal laser-scanning microscope (Nikon, Tokyo, Japan), equipped with a Plan Apo 60× water immersion objective [NA=1.27] and a PicoQuant - TCSPC-FCS upgrade kit (PicoQuant, Berlin, Germany). Fluorescence of the CF-tagged peptide was excited by the 488 nm laser line, for Cy5 a 635 nm laser line was used. Fluorescence of CF and Cy5 were filtered with 500–550 nm and 660–740 nm bandpass filters, and detected with single photon counting detectors (PicoQuant, Berlin, Germany). Measurements of 20 x 10 second runs were taken with each samples. Fluorescence autocorrelation curves were calculated using the SymPhoTime64 software (PicoQuant, Berlin, Germany) at 200 time points from 300 ns to 1 s distributed on a quasi-logarithmic time scale. For fitting of the autocorrelation curves to obtain diffusion times and fraction of DNA- or nucleosome bounded C9-CF, see Supplementary Methods.

Since the diffusion coefficients (*D)* of the measured fluorescein and Cy5 samples depend on the viscosity of the environment (through the Stokes–Einstein equation), the calculated values of the lateral e^−2^ radius of the detection volumes are also affected by the refractive index of the solution ^79^ likely in a wavelength-dependent manner. Therefore, the ratio of the lateral e^−2^ radii of the detection volumes with eq.3 was calculated for 5 mM as well as 700 mM NaCl.

For determining the fraction of unlabeled gDNA bounded C9-CF, τ_D2_ was not fixed in the fitting of the autocorrelation functions as it may have a broad size distribution. All correlation curves were fitted using the QuickFit 3.0 software.

### Data Availability

The Whole Genome Sequencing datasets generated during the current study are available in the NCBI SRA repository, under the BioProject accession number PRJNA853352, WGS BioSample accession numbers (SAMN29379375, SAMN29379376, SAMN29379377, SAMN29379378). Microarray datasets were deposited into GEO database under accession number GSE225680. The RNA-seq datasets generated during the current study are available in the NCBI SRA repository, under the BioProject accession number PRJNA853352, BioSample accession numbers (SAMN32271849, SAMN32271848, SAMN32271847, SAMN32271846, SAMN32271845, SAMN32271844, SAMN32271843, SAMN32271842, SAMN32271841). Mass spectrometric data are available via ProteomeXchange with identifier PXD040998. All other datasets generated during and/or analysed during the current study are available from the corresponding author on reasonable request.

## ACKNOWLEDGEMENTS

The authors thank Cyclolab Hungary for providing cyclodextrins, Sandra B. Hake (Giessen, Germany) for the PWWP2A-GFP plasmid, Kerstin Bystricky (Toulouse, France) for the gift of anti-H2A.Z antibody ab4174, Zheng Zhou (Beijing, China) for explanations regarding their cryo-EM data on H2A.Z structure, Hiroshi Kimura (Tokyo, Japan) for valuable methodical advice and gift of antibodies, Adel Vezendine Nagy for technical help. Microscopy measurements were carried out at the Debrecen sub-Node of the Cellular Imaging Hungary Euro-BioImaging Node. Mass spectrometry analysis was carried out at the BMBI Proteomics Core Facility, Department of Biochemistry and Molecular Biology, University of Debrecen. The authors thank Mr. Soma Godó (UNICAM Ltd., Budapest, Hungary) for loaning the STEDYCON microscope. The authors acknowledge the support of the Freiburg Galaxy Team, Bioinformatics, University of Freiburg (Germany) funded by the Collaborative Research Centre 992 Medical Epigenetics (DFG grant SFB 992/1 2012) and the German Federal Ministry of Education and Research BMBF grant 031 A538A de.NBI-RBC.

## CONTRIBUTIONS

IL: LSC experiments, confocal microscopic studies, texture analyses, cytotoxicity measurements; MH and MK: microarray data; RB and ÁCs: bacterial production of plasmids; PN: analyses of microarray data provided by MH, bioinformatics analyses of sequencing data, preparation of reconstituted nucleosomes, doing the FCS measurements and data analyses together with GMo; GSz: conceptualization of the experiments, evaluation of data and writing of manuscript together with IL; MH: conceptualization of the DT40 experiments; GMe supervised peptide synthesis conducted by KNE; JA: advice and editing of the manuscript, MT developed the inducible GFP-H2A.Z system and provided advice regarding the CUT&RUN experiments; DT: contributed the hydroxyapatite elution data and edited the manuscript; EC and ZD: performed MS data analyses; IB conducted the titration experiments with cyclodextrins; BM: advice on melanoma cell lines; KT provided the constituents for and supervised the nucleosome reconstitution experiments, and supervised the analyses of the FCS data; GV discussed the FCS data and edited the manuscript; SP and BS contributed to the bioinformatic analyses of the RNA-seq data.

