## Supplementary figures and text for "EPIGENETIC MODULATION VIA THE C-TERMINAL TAIL OF H2A.Z"

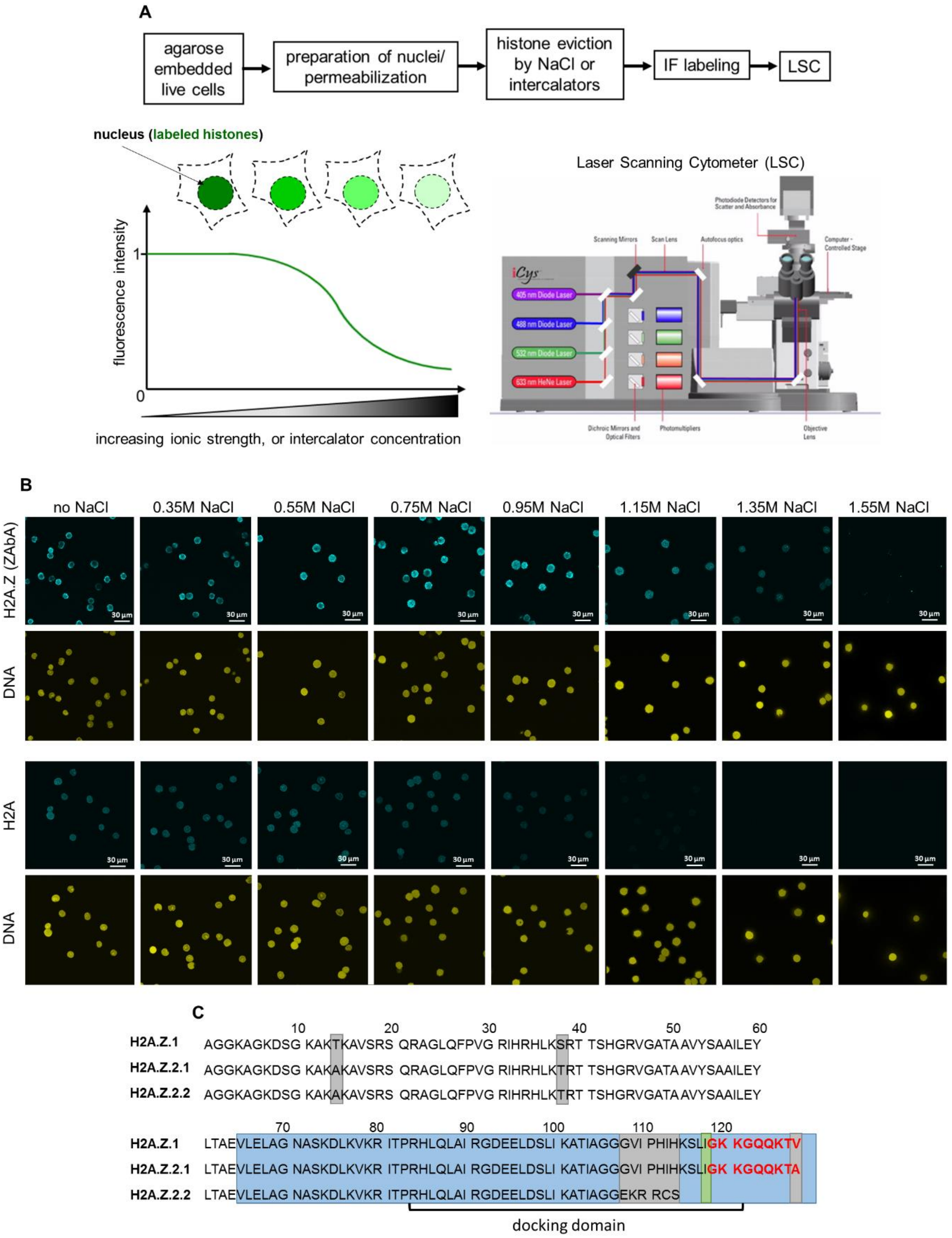

**Suppl. Fig. 1**

(A) Flow-chart of the QINESIn assay (Quantitative Imaging of Nuclei after Elution with Salt/Intercalators assay; [1]). Histones remaining in the nuclei after treatment with increasing concentration of NaCl solutions (or intercalator; an option not used in the current work) are detected by indirect immunofluorescence labeling (or directly if the histones are tagged with a fluorescent protein) and quantitatively analyzed by laser scanning cytometry (LSC). Our LSC instrument is equipped with four lasers: 405 nm, 488 nm, 563nm, 633 nm and four photomultiplier tubes (PMTs), each detecting a specific wavelength range of fluorescence excited by the scanning lasers. As the laser light intersects the sample, scattered or transmitted light is also simultaneously directed to one or more photomultiplier tubes. The PMT signals are converted into images and the events, such as cells, nuclei or other subcellular structures are identified and segmented on the basis of their fluorescence. From the segmented events, a variety of quantitative data are calculated: area, integral fluorescence, maximal pixel intensity, circularity, perimeter, *x* and *y* coordinates, among others. The numerical values are displayed in scattergrams and histograms, allowing assessment of relationships among the various features. (B) Representative CLSM images showing ZAbA-stained H2A.Z and immunolabeled H2A in permeabilized HeLa nuclei exposed to different concentrations of NaCl. (C) Amino acid sequences of human H2A.Z.1, H2A.Z.2.1, H2A.Z.2.2. Grey boxes highlight sequences that are different among H2A.Z isoforms. Blue square indicates the C-terminal epitope recognized by ZAbA starting with amino acid 65, comprising the H2A.Z docking domain and the C-terminal unstructured tail. ZAbB is generated against a peptide sequence surrounding amino acid residue 118 (green box) of the docking domain. (Further details are not provided by the manufacturer.) The 9 amino acid long deletion in the ΔC H2A.Z mutant is shown in red font.

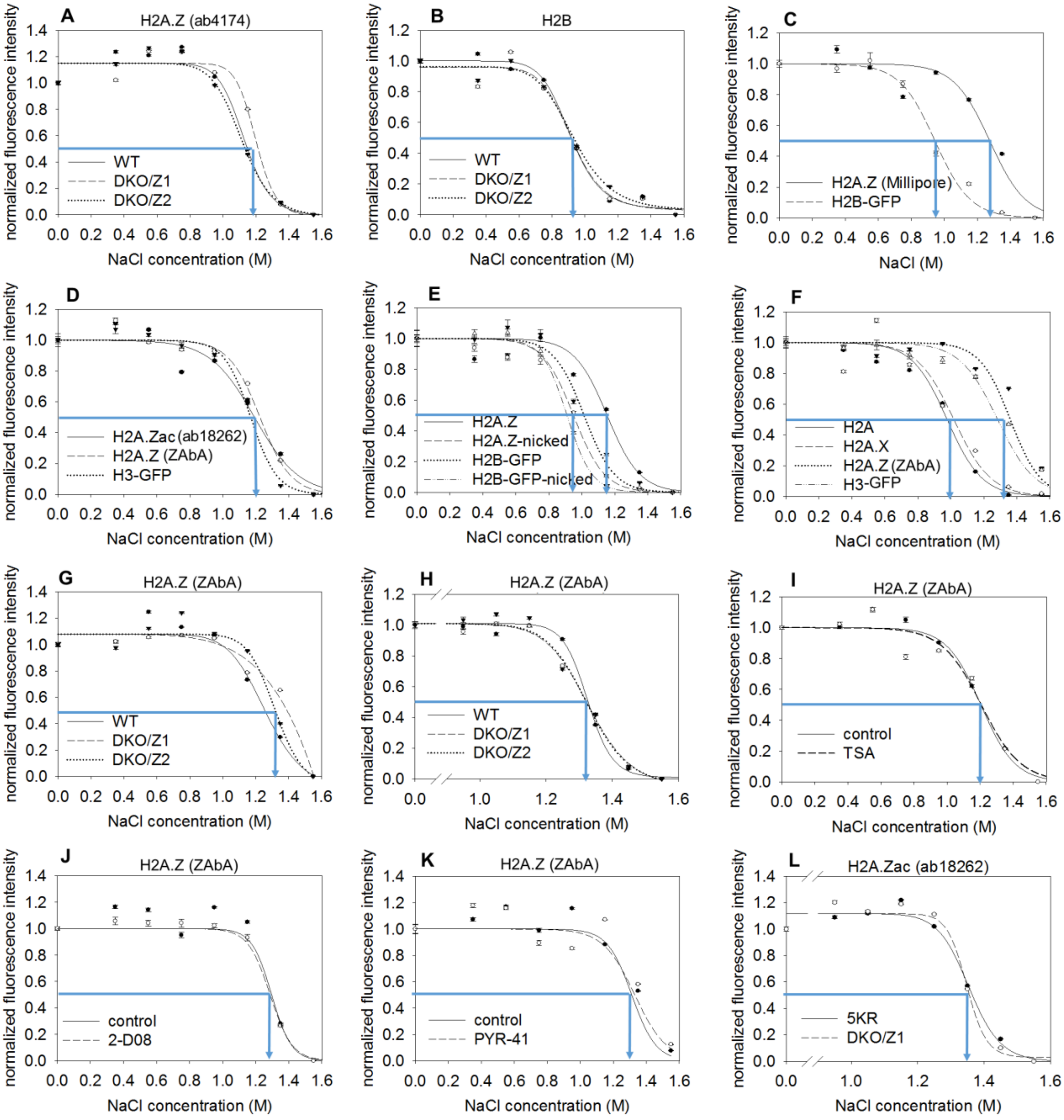

**Suppl. Fig. 2.**

(A) Salt elution profile of H2A.Z isoforms, detected by ab4174 (clone different from that of ZAbA; see Materials and Methods) in the nuclei of H2A.Z.1 (DKO/Z1) or H2A.Z.2 (DKO/Z2) expressor double knock-out chicken DT40 cells, compared to wild type (WT) cells. (B) H2B elution curves in the experiment shown in panel A; H2B, co-labeled with H2A.Z, serves as an internal control. (C) Salt elution profile of H2A.Z detected by the Merck-Millipore 07-594 antibody in H2B-GFP expressor HeLa cells. (D) Salt elution profile of H2A.Z detected by an antibody recognizing acetylated H2A.Z (ab18262). H3-GFP was used as inner control; bulk H2A.Z was detected by ZAbA and measured in a parallel sample. (E) Salt elution profiles of H2A.Z (detected by ZAbA) in H2B-GFP expressor HeLa nuclei before and after 0.5 U/ml nickase treatment. (F) Salt elution curves of Figure 1A (see main text) normalized only to the initial value (1.0). (G) Salt elution profile of H2A.Z isoforms labeled with ZAbA in the nuclei of H2A.Z.1 or H2A.Z.2 expressor DKO DT40 cells, compared to WT. (H) Experiment in panel G repeated to focus on the salt concentration range between 0.95 and 1.55 M. (I-K) The possible effect of acetylation (I), sumoylation (J) and ubiquitination (K) assessed by salt elution using ZAbA, after treatment with trichostatin (TSA), 2-D08 or PYR-41, respectively. (L) Salt elution profile of nucleosomes in H2A.Z.1-5KR (5KR) and H2A.Z.1 expressor DKO DT40 cells, detected by acetylated H2A.Z specific antibody (ab18262). In the H2A.Z.1-5KR mutant [2], 5 acetylatable lysines on the H2A.Z N-terminus were changed to arginines. The elution curves refer to G1 phase cells gated according to their DNA fluorescence intensity distribution and the error bars represent SEM of ~600 G1 nuclei measured by LSC. Blue arrows on the elution curves indicate EC50 values (also in the other figures).

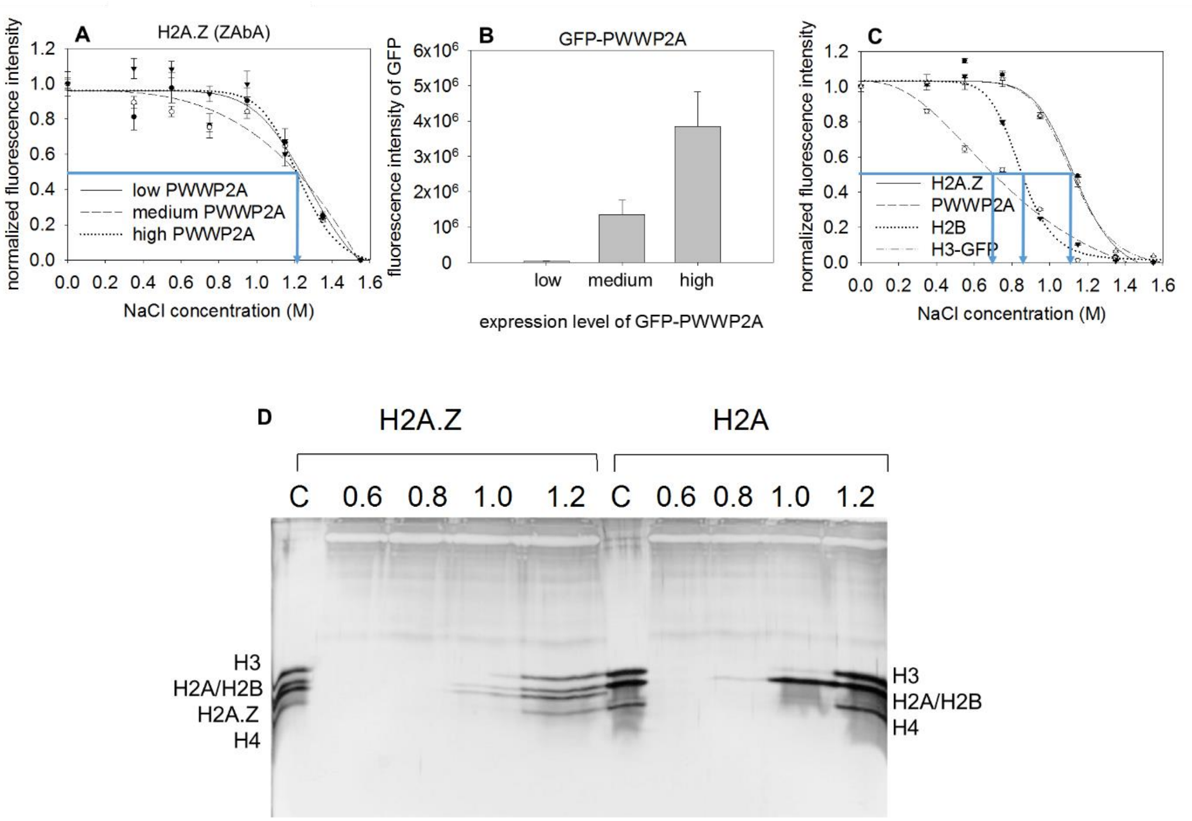

**Suppl. Fig. 3.**
(A) Salt elution profiles of H2A.Z detected by ZAbA in high, medium and low GFP-PWWP2A expressor HeLa nuclei. (B) GFP fluorescence levels in the different subpopulations of HeLa cells gated according to GFP-PWWP2A to record the distribution curves used for panel (A). (C) Salt elution profiles of PWWP2A (detected by immunofluorescence, H2B-GFP, H3-GFP, and H2A.Z (detected by ZAbA). PWWP2A was labeled in H2B-GFP expressor HeLa cells. H2A.Z was detected by ZAbA and measured in the nuclei of H3-GFP-expressor HeLa. (D) Full gel image of Fig. 1G

| Accession | Description | # Peptides | # PSMs | Coveage, % |
|---|---|---|---|---|
| O60216 | RAD21 | 3 | 3 | 9 |
| P02545 | Prelamin-A/C | 43 | 126 | 62 |
| P07305 | H1.0 | 3 | 3 | 12 |
| P0C0S5 | H2A.Z.1 | 3 | 34 | 24 |
| P10412 | H1.4 | 14 | 66 | 43 |
| P16401 | H1.5 | 10 | 16 | 31 |
| P16403 | H1.2 | 4 | 18 | 18 |
| P20700 | Lamin-B1 | 25 | 37 | 42 |
| P45973 | HP1α | 5 | 5 | 30 |
| Q03252 | Lamin-B2 | 13 | 15 | 21 |
| Q5QNW6 | H2B | 9 | 218 | 58 |
| Q71UI9 | H2A.Z.2 | 3 | 34 | 24 |

**Suppl. Table. 1.** Proteins detected by mass spectrometry in halo samples shown on Fig. 1F. Proteins identified in the Swissprot database. Swissprot database accession numbers, number of unique peptides identified (# Peptides), the number of spectra assigned to the protein (# PSM) and polypeptide sequence coverage % are also shown for these proteins not detected earlier in the lamina.

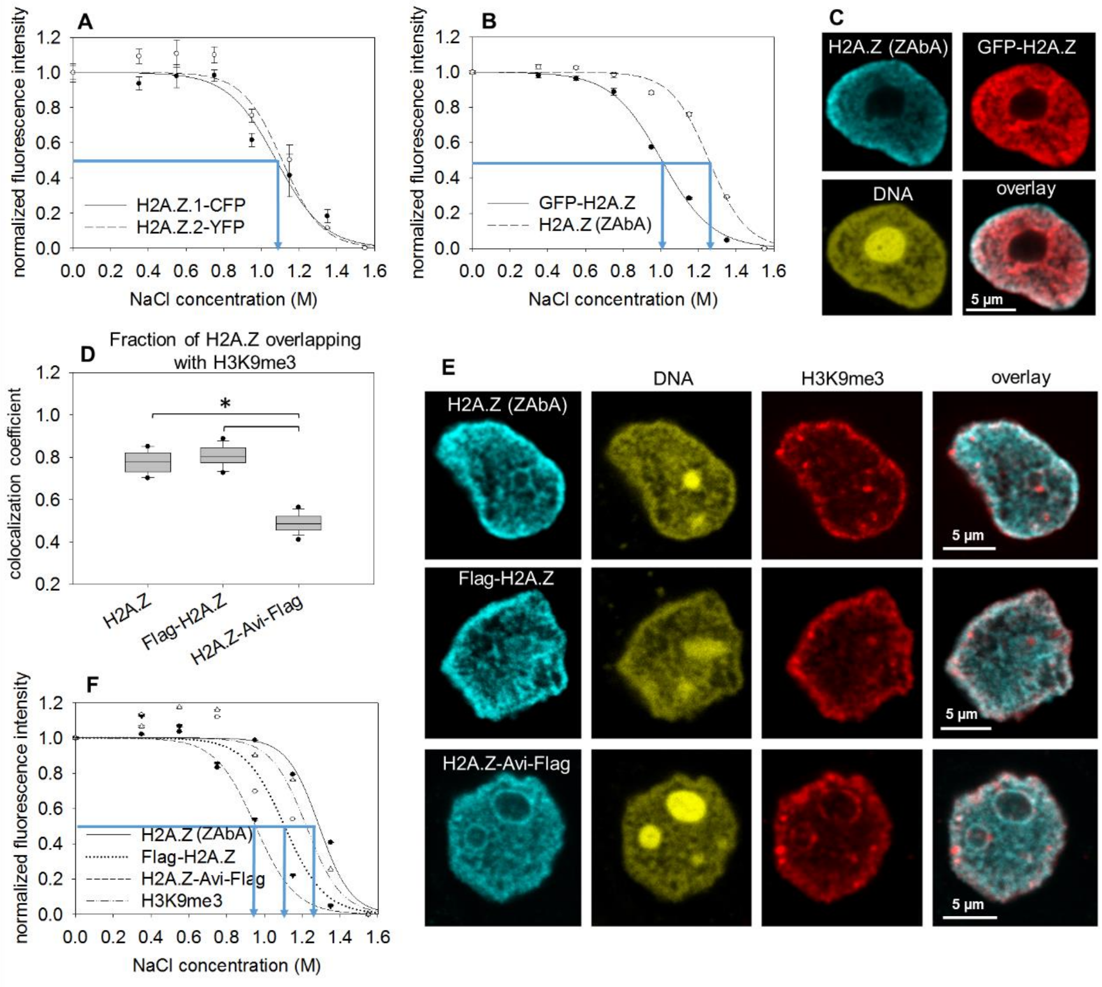

**Suppl. Fig. 4.**

(A) Salt elution curves of H2A.Z.1-CFP and H2A.Z.2-YFP tagged on their C-terminus, expressed from plasmids transfected into HeLa. (B) Salt elution curves of GFP-H2A.Z (tagged on its N-terminus) and of the endogenous H2A.Z labeled in the same sample using ZAbA. (C) CLSM images showing the localization of H2A.Z detected by ZAbA in GFP-H2A.Z expressor HeLa nuclei. (D) Measurement of colocalization of H2A.Z, Flag-H2A.Z or H2A.Z-Avi-Flag with H3K9me3. Manders colocalization coefficient was calculated showing the fraction of H2A.Z positive pixels overlapping those of H3K9me3 in HeLa nuclei. Box-and-whisker plot was created from the data of ~17 nuclei. (E) CLSM images showing the localization of H2A.Z, Flag-H2A.Z or H2A.Z-Avi-Flag co-labeled with H3K9me3 in HeLa nuclei. H2A.Z was detected by ZAbA. (F) Salt elution curves of Flag-H2A.Z and H2A.Z-Avi-Flag containing nucleosomes measured in HeLa nuclei. Tagged H2A.Z histones were detected by anti-Flag antibody. Non-tagged H2A.Z was detected by ZAbA and measured in a parallel sample. Elution curves refer to G1 phase cells gated according to their DNA fluorescence intensity distribution and the error bars represent SEM of ~600 G1 nuclei measured by LSC. Statistical analysis was done using one-way ANOVA (* $p \leq 0.001$)

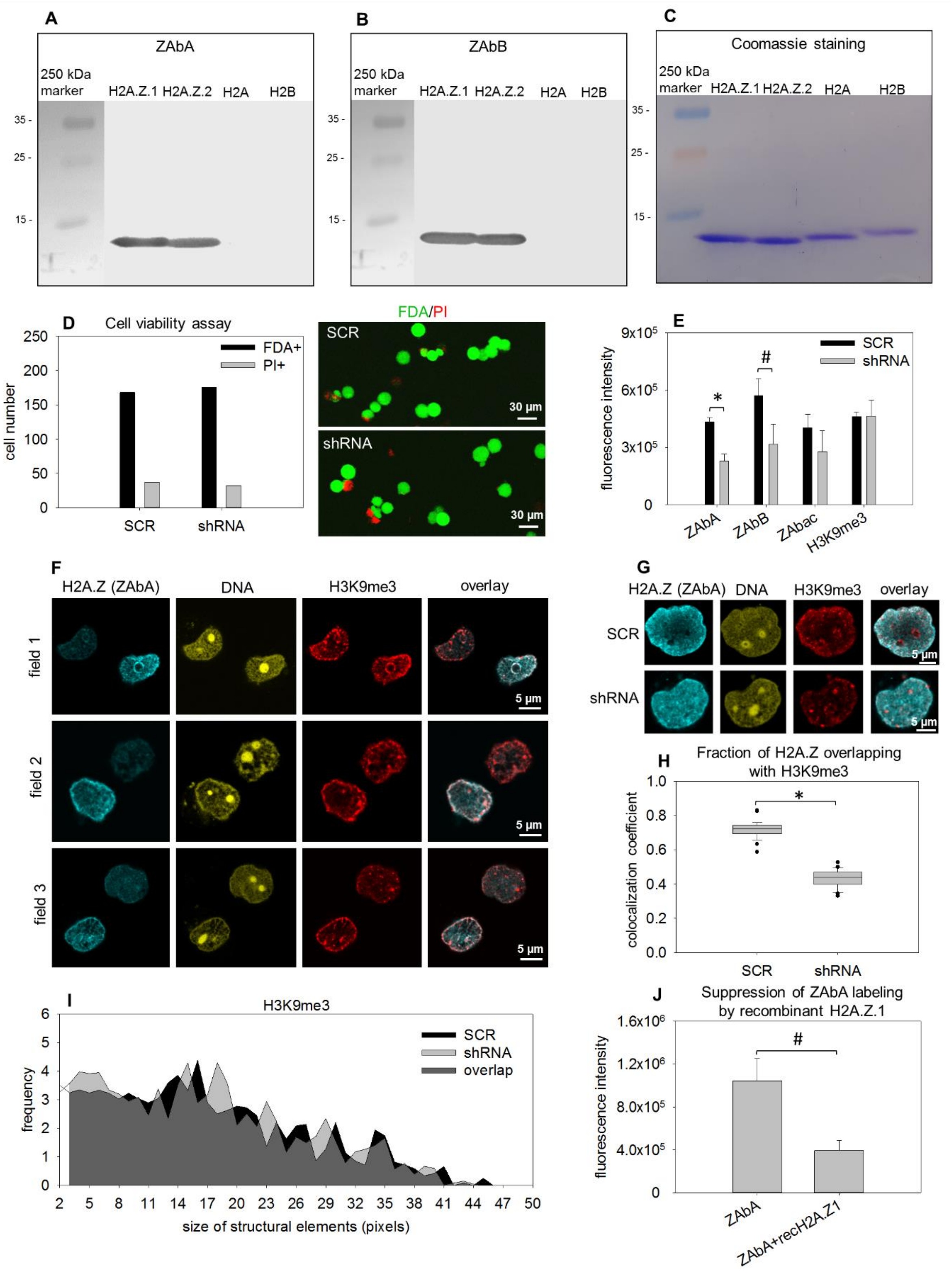

**Suppl. Fig. 5.**

(A and B) Specificity of ZAbA (A) and ZAbB (B) was validated on Western blots using recombinant human H2A.Z.1, H2A.Z.2, H2A and H2B histones (see Materials and Methods). (C)

SDS-PAGE analysis of the recombinant histones stained with Coomassie blue dye. (D) Viability analysis by fluorescein-diacetate (FDA) and propidium iodide (PI) staining of cells transfected with plasmid constructs expressing scrambled shRNA (SCR) or shRNA targeting the H2A.Z gene (shRNA). (E) Amount of H2A.Z, as detected by ZAbA, ZAbB, the antibody thought to recognize H2A.Zac (ab18262; ZAbac) and H3K9me3 in the nuclei of the transfected cells measured by LSC. Average and SD values calculated from 3 parallel measurements are shown on the bar chart. (Statistical analysis was done using one-way ANOVA, $^*p \leq 0.001$, $\# \ p \leq 0.05$, also in the subsequent figures). (F and G) CLSM images showing the nuclear localization of ZAbA detected H2A.Z co-labeled with H3K9me3 in the nuclei of cells transfected with the shRNA and SCR constructs. The fluorescence signals of the silenced sample (shRNA) was amplified relative to the control (SCR) in (G) to emphasize the shRNA-induced alteration in topology. (H) Colocalization analysis of of H2A.Z and H3K9me3 measured as the Manders colocalization coefficient showing the fraction of H2A.Z overlapping with H3K9me3. Box-and-whisker plot shows the median, $25^{th}$ and $75^{th}$ percentiles as vertical boxes with error bars, $5^{th}$, $95^{th}$ percentiles and outliers as dots created from the data of ~28 nuclei. (I) Texture analysis of SCR or shRNA transfected HeLa nuclei showing the size distribution of structural elements containing H3K9me3. Distribution curves were generated from the mean values of ~28 cells. (J) Suppression of ZAbA labeling by recombinant H2A.Z.1 in HeLa nuclei measured by LSC. Means of immunofluorescence intensity distributions with SD values calculated from 3 parallel measurements are shown.

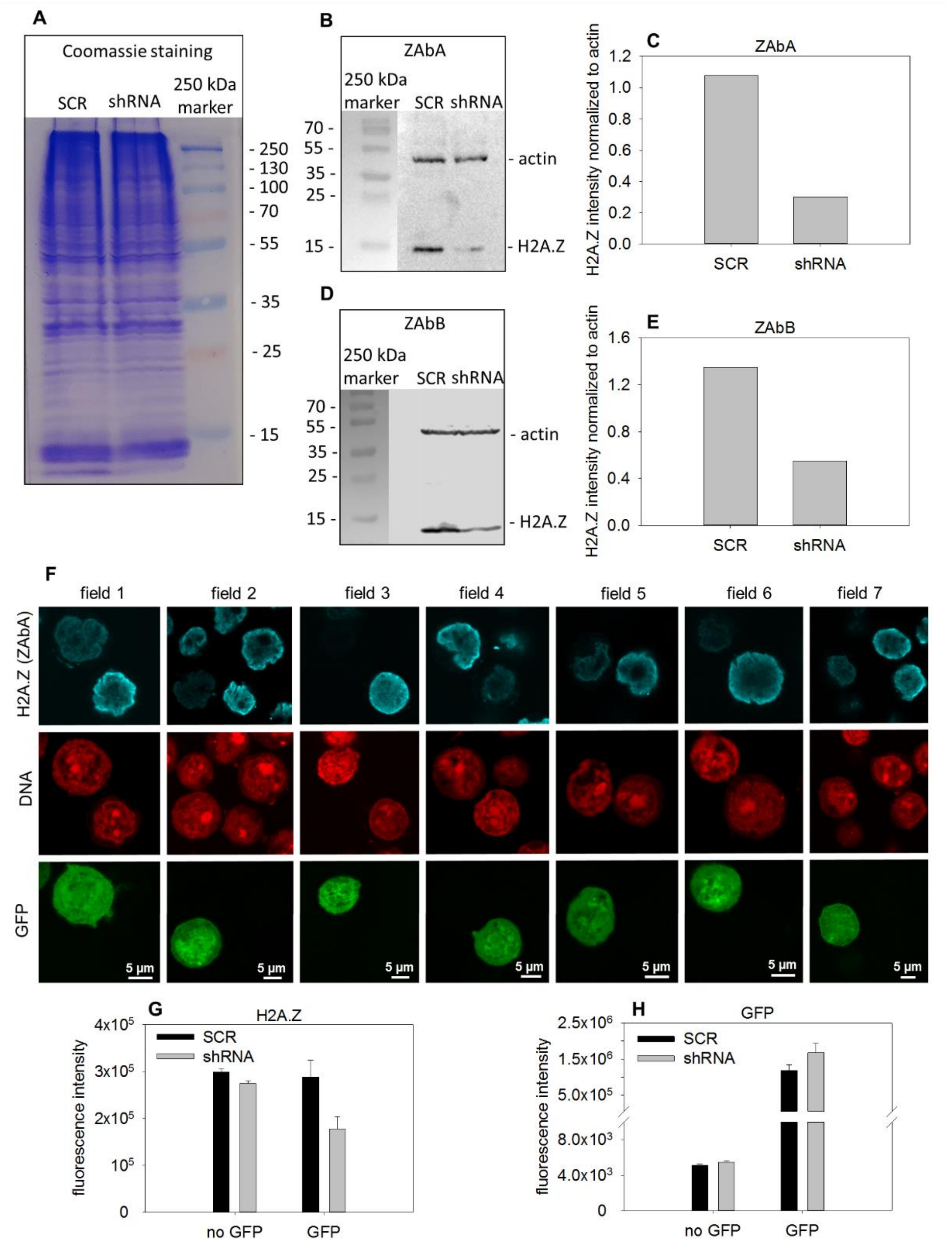

**Suppl. Fig. 6.**

(A) SDS-PAGE of cell lysates of HeLa cells transfected with SCR, or shRNA, stained with coomassie blue dye. (B) Western blot analysis of SDS-PAGE from panel (A) using ZAbA antibody. (C) Bands from panel (B) quantitatively analyzed by ImageJ. Bar chart shows the background subtracted mean pixel intensities of H2A.Z normalized to actin. (D) Western blot analysis of SDS-PAGE from panel (A) using ZAbB antibody. (E) Bands from panel (D) were quantitatively analyzed by ImageJ as in panel (C). (F) Representative fields from scans of 293T cells co-transfected with shRNA and GFP expressing plasmids. (G, H) Silencing of H2A.Z in co-transfected cells measured by LSC, gating GFP positive (GFP) and negative (no GFP) cells. Scrambled shRNA (SCR) was used as negative control. The shRNA constructs were from ref. [3]. Error bars represent SEM of ~600 G1 nuclei measured by LSC.

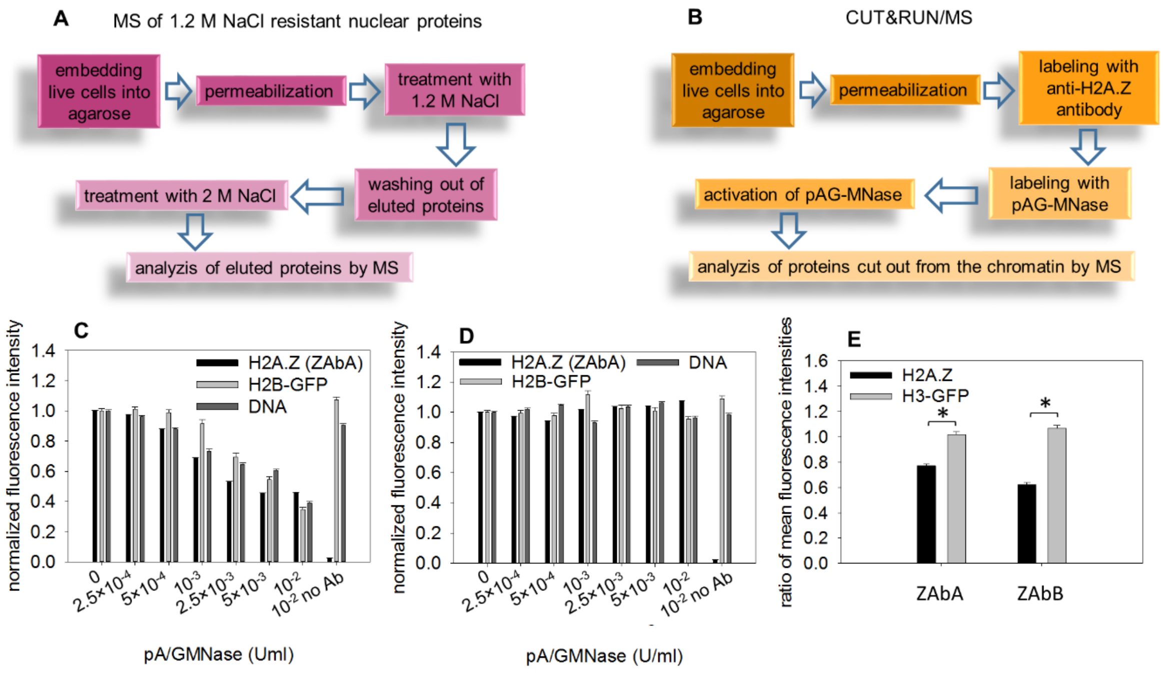

**Suppl. Fig. 7.**
(A) Flow chart of the elution experiment combined with mass spectrometry (MS). Proteins resistant to 1.2 M salt were detected by MS after high salt elution. (B) Flow chart of the CUT&RUN/MS experiment. ZAbA labeled nucleosomes released by the protein A/G-tagged MNase were detected by MS as described in Materials and Methods. (C) Titration of protein A/G-tagged MNase in H2B-GFP expressor HeLa nuclei. H2A.Z (labeled by ZAbA) and H2B-GFP were analyzed after MNase treatment by LSC. (D) The same as (C) when the MNase was not activated by $CaCl_2$. (E) The effect of protein A/G MNase treatment used at $10^{-3}$ U/ml concentration, using ZAbA and ZAbB, in H3-GFP HeLa nuclei. The ratios of mean fluorescence intensities observed after MNase activation and in the absence of $CaCl_2$ are shown. Statistical analysis was done using one-way ANOVA (* $p \leq 0.001$).

| CUT&RUN | | | | NaCl resistant fraction | | | | | |
|---|---|---|---|---|---|---|---|---|---|
| ZAbA | | ZAbB | C9+ZAbA | | | | | | |
| ACIN1 | NOLC1 | ANXA2 | ACIN1 | APEX1 | EBNA1BP2 | HNRNPU | PA2G4 | RNMT | S100A8 |
| ACTG1 | NONO | BASP1 | ALDOA | DDX6 | EEF1A1 | HP1BP3 | PABPC1 | RPF2 | S100A9 |
| AHNAK | NOP58 | BOP1 | BASP1 | mH2A.1 | EEF1G | HSP90AA1 | PABPC4 | RPL10A | SERBP1 |
| ALDOA | NUMA1 | CALM1 | CALM1 | ACTG1 | EEF2 | HSPA1A | PABPN1 | RPL11 | SERPINB3 |
| ANXA2 | PA2G4 | HP1α | HP1α | ACTN4 | EIF2AK2 | HSPA4 | PAK1IP1 | RPL13 | SF3A1 |
| BANF1 | PABPN1 | HP1β | HP1β | ACTR2 | EIF2S1 | HSPA8 | PARP1 | RPL17 | SF3A3 |
| BASP1 | PARP1 | HP1γ | HP1γ | ACTR3 | EIF4A1 | HSPH1 | PCBP1 | RPL22 | SF3B1 |
| BOP1 | PCBP2 | CFL1 | DDX21 | AHNAK | EIF4A3 | IFI16 | PCBP2 | RPL23 | SFN |
| CALM1 | PDIA3 | DHX9 | DHX9 | AIFM1 | ELAVL1 | IGF2BP3 | PDCD11 | RPL23A | SFPQ |
| HP1α | PES1 | EEF1A1 | EEF1A1 | AIMP1 | EMG1 | ILF2 | PDIA3 | RPL26 | SLC3A2 |
| HP1β | PKM | EEF1D | EEF1D | AIMP2 | ENO1 | ILF3 | PEBP1 | RPL27 | SND1 |
| HP1γ | PPIA | EEF1G | EIF6 | AKR1B1 | ERH | IPO5 | PFN1 | RPL3 | SNRNP40 |
| CFL1 | PRDX1 | EIF6 | FLNA | ALDH7A1 | EZR | KARS | PHGDH | RPL30 | SNRPB |
| DDX21 | PRPF19* | ENO1 | G3BP2 | ALDOA | FBL | KHSRP | PIP | RPL35 | SNRPD3 |
| DDX3X | PSMB6 | FLNA | H1.0 | ALYREF | FKBP4 | KPNB1 | PKM | RPL5 | SNU13 |
| DHX9 | RALY | H1.4 | H2A | ANXA1 | FLNA | KRR1 | PNO1 | RPL6 | SRP14 |
| DKC1 | RBBP7 | HSPA1A | H2B* | ANXA2 | FLNB | KYNU | PNO1 | RPL7 | SRSF1 |
| DSP | RRP9 | HSPA5 | HSPA1A | ANXA5 | FUS | LARP1 | PNP | RPL7A | SRSF3 |
| EEF1A1 | RRP9 | HSPA8 | HSPA5 | ARPC4 | G3BP1 | LARS | POLR1C | RPL7L1 | SRSF9 |
| EEF1B | RSL1D1 | HSPA9 | HSPA8 | ASS1 | GAPDH | LDHA | PPIB | RPL9 | SSB |
| EEF1D | SAFB | LMNA | HSPA9 | BANF1 | GAR1 | LGALS1 | PPP1CA | RPLP0 | STIP1 |
| EEF1G | SART1 | MKI67 | LMNA | BASP1 | GLO1 | LMNA | PPP2R1A | RPLP2 | STRAP |
| EEF2* | SF3A3 | MYH9 | LMNB1 | BAZ1B | GNAS | LRPPRC | PRDX1 | RPP30 | SYNCRIP |
| EIF6 | SF3B2 | NCL | MATR3 | BRIX1 | GPI | LRRC59 | PRDX5 | RPS10 | TALDO1 |
| FLNA | SF3B3 | PKM | MKI67 | BUB3 | GSTM3 | LSM2 | PRDX6 | RPS11 | TKT |
| FTSJ3 | SFPQ | PPIA | MYH9 | CALM1 | H1.0 | LTF | PRPF19 | RPS12 | TOP1 |
| FUBP1 | SLTM | PTBP1 | NCL | CALR | H1.10 | LYAR | PRPF3 | RPS13 | TOP2A |
| G3BP1 | SNRPA1 | RALY | NOLC1 | HP1γ | H1.2 | MAGOHB | PRPF4 | RPS14 | TPI1 |
| GAPDH | SNRPB | RPS7 | NUMA1 | CCDC86 | H2A | MARS | PSIP1 | RPS15A | TPT1 |
| H2A | SNRPD2 | RRP9 | PCBP2 | CCT4 | H2A.Z1 | MATR3 | PSMA4 | RPS16 | TRMT112 |
| H2B* | SNRPD3 | RUVBL2 | PHF5A | CCT8 | H2A.Z2 | MKI67 | PSMB4 | RPS17 | TROVE2 |
| HSP90AB1 | SNW1 | SAFB | RBBP4 | CFL1 | H2B | MRTO4 | PSMD11 | RPS19 | TUBA1B |
| HSPA1A | SPTBN1 | SF3B3 | RPS7 | CSE1L | H3.2 | MRTO4 | PSMD14 | RPS2 | TUBB |
| HSPA5 | SRSF2 | SFPQ | RRP9 | DAZAP1 | H4 | MSN | PSME3 | RPS24 | TUBB4B |
| HSPA8 | SRSF3 | SLTM | SAFB | DCAF13 | HDGF | MYBBP1A | PTBP1 | RPS25 | TXNRD1 |
| HSPA9 | TKT | SNRPD1 | SF3B3 | DCAF13 | HDLBP | MYO1C | PUM3 | RPS27A | U2AF2 |
| IK | TRA2A | SNRPD2 | SFPQ | DDX17 | HMGN2 | NAP1L1 | RACK1 | RPS3 | UBA1 |
| LAP2B | TUBA1B | SNRPD3 | SNRPD2 | DDX18 | HNRNPA0 | NAP1L4 | RALY | RPS3A | USP39 |
| LMNA | TUBB | TKT | SNRPD3 | DDX21 | HNRNPA1 | NCL | RAN | RPS4X | WDR1 |
| LMNB1 | UBA1 | TUBA1B | SRRM2 | DDX23 | HNRNPA2B1 | NHP2 | RANBP1 | RPS5 | WDR46 |
| LSM3 | VCP | TUBB | SRSF1 | DDX27 | HNRNPAB | NIFK | RANGAP1 | RPS6 | WDR74 |
| MATR3 | YWHAE | VCP | TUBA1B | DDX39B | HNRNPC | NIP7 | RBBP4 | RPS7 | XRCC5 |
| MDC1 | APEX1* | APEX1* | VCP | DDX3X | HNRNPD | NLE1 | RBM28 | RPS8 | XRCC6 |
| MKI67 | SMARCA5* | SMARCA5* | APEX1* | DDX47 | HNRNPDL | NONO | RBM34 | RPS9 | YBX1 |
| MSN | CDC5L* | CDC5L* | SMARCA5* | DDX5 | HNRNPF | NOP2 | RBM39 | RRP9 | YWHAE |
| MYH9 | H2A.Z1* | H2A.Z1* | CDC5L* | DEK | HNRNPK | NOP56 | RBMX | RRS1 | YWHAZ |
| MYO1C | MCM4* | MCM4* | H2A.Z1* | DHX9 | HNRNPL | NOP58 | RCC1 | RSL1D1 | |
| NCL | MCM5* | MCM5* | MCM4* | DKC1 | HNRNPM | NPM1 | RCL1 | RTCB | |
| NOL6 | MCM7* | MCM7* | MCM5* | DRG1 | HNRNPR | NSUN2 | REXO4 | RTRAF | |
| | CSE1* | PRPF19* | MCM7* | | | | | | |
| | PSMA6* | CSE1* | PRPF19* | | | | | | |
| | U2AF1* | PSMA6* | CSE1* | | | | | | |
| | | H2B* | PSMA6* | | | | | | |
| | | EEF2* | EEF2* | | | | | | |
| | | U2AF1* | U2AF1* | | | | | | |

**Suppl. Table 2.** Proteins released by 1.2 M salt- or pA/G-MNase.

Proteins known to be H2A.Z associated [4, 5, 6] are labeled in the table with blue color. Histones are labeled with green and proteins responsible for higher order chromatin structure (heterochromatin, matrix associated proteins) are labeled with red. Asterisk indicates proteins detected in the experiment related to Fig. 3N and Suppl. Table 4, using also a secondary antibody.

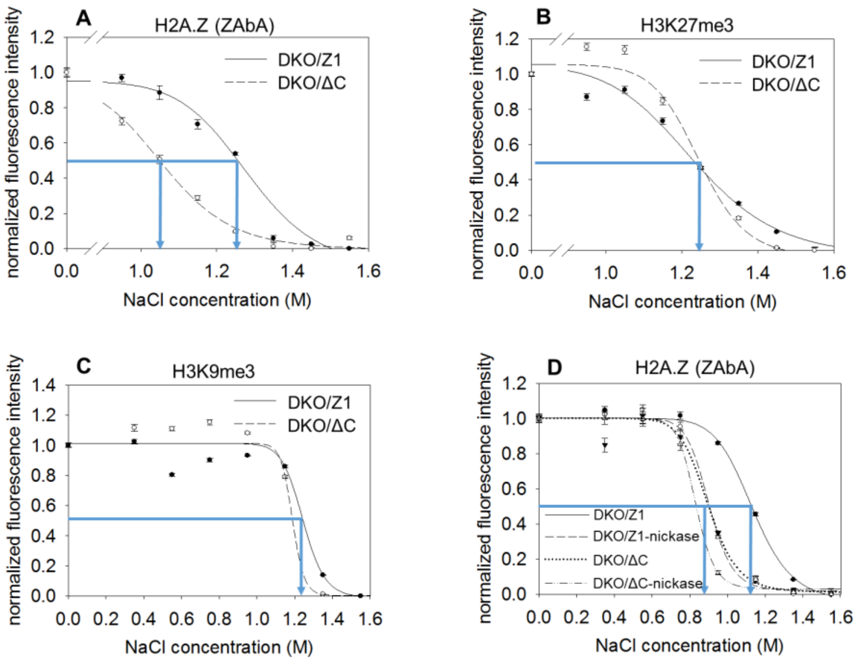

**Suppl. Fig. 8.**

(A) Experiment in Fig. 2A was reproduced focusing on the salt concentration range of 0.95-1.55 M. (B and C) Salt elution profiles of H3K27me3 (B) and H3K9me3 (C) measured in DKO H2A.Z.1 (DKO/Z1) and H2A.Z.1ΔC (DKO/ΔC) DT40 nuclei (resolving the salt concentration range as in panel A of this figure or as in Fig. 2A, respectively). (D) Salt elution profiles of H2A.Z (detected by ZAbA) in DKO/ΔC and DKO/Z1 nuclei before and after 0.5 U/ml nickase treatment. The elution curves refer to G1 phase cells gated according to their DNA fluorescence intensity distribution and the error bars represent SEM of ~600 G1 nuclei measured by LSC.

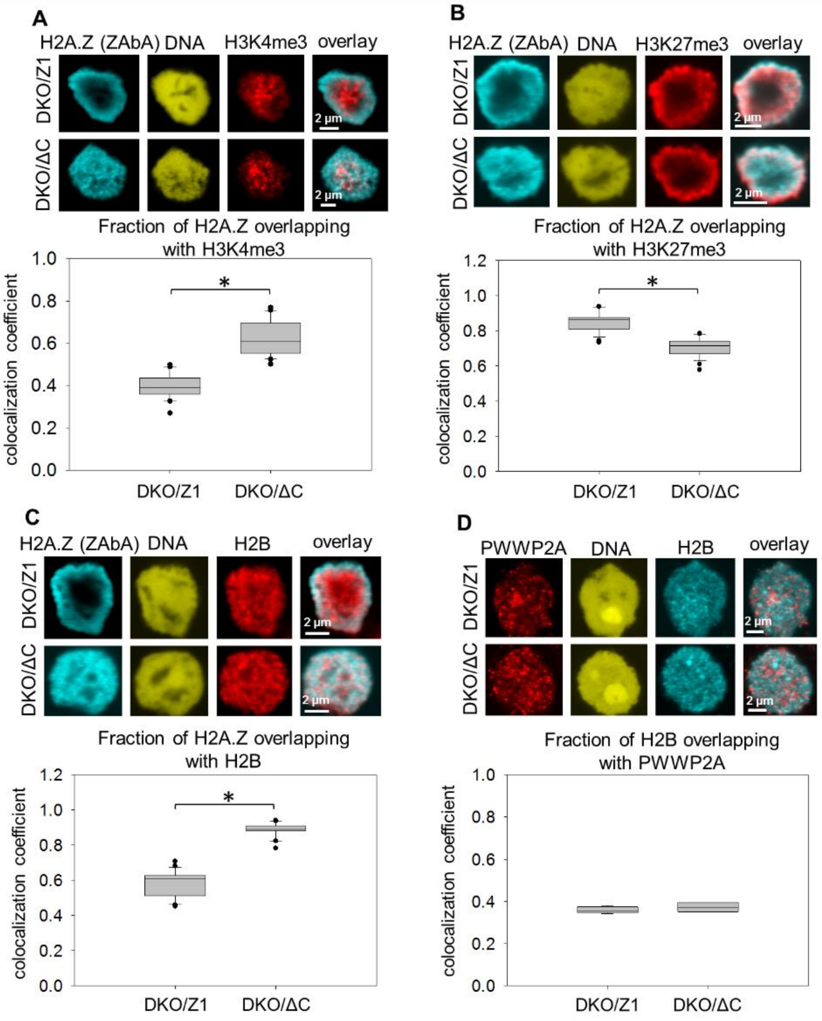

**Suppl. Fig. 9.**

(A-D) Colocalization measurements: (A) H2A.Z and H3K4me3, (B) H2A.Z and H3K27me3, (C) H2A.Z and H2B, (D) H2B and PWWP2A colocalization in DKO/Z1 and DKO/ΔC nuclei. H2A.Z was detected by ZAbA, PWWP2A was labeled with antibody specific for the protein. Manders colocalization coefficients representing the fraction of H2A.Z overlapping with H3K4me3 (A), with H3K27me3 (B), with H2B (C), or the fraction of H2B with PWWP2A (D) are shown. Statistical analysis was done using one-way ANOVA (* $p \leq 0.001$). Box-and-whisker plot shows the median, $25^{th}$ and $75^{th}$ percentiles as vertical boxes with error bars, $5^{th}$, $95^{th}$ percentiles and outliers as dots created from the data of 20 (A), ~27 (B), ~24 (C) or ~4 (D) nuclei.

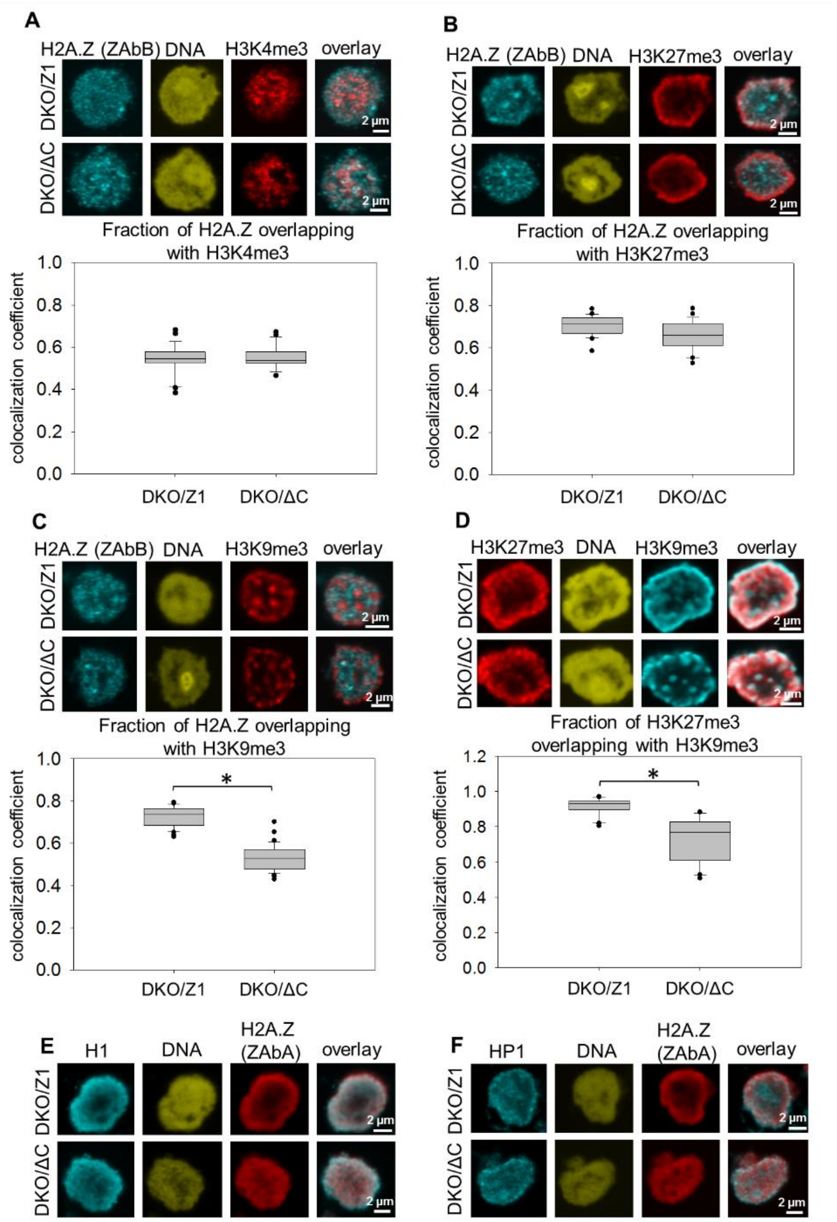

**Suppl. Fig. 10.**

(A-D) Colocalization measurements: (A) H2A.Z and H3K4me3, (B) H2A.Z and H3K27me3 (C), H2A.Z and H3K9me3, (D) H3K27me3 and H3K9me3 colocalization in DKO/Z1 and DKO/ΔC nuclei. H2A.Z was detected using ZAbB. Manders colocalization coefficients showing the fraction of H2A.Z overlapping with H3K4me3 (A), H3K27me3 (B) and H3K9me3 (C), and of H3K27me3 overlapping with H3K9me3 (D) were calculated. Statistical analysis was done using one-way ANOVA (* $p \leq 0.001$). Box-and-whisker plot shows the median, 25$^{th}$ and 75$^{th}$ percentiles as vertical boxes with error bars, 5$^{th}$, 95$^{th}$ percentiles and outliers as dots created from the data of ~24 (A), ~26 (B), ~42 (C) or 25 (D) nuclei. (E and F) CLSM images of H2A.Z-containing nucleosomes detected by ZAbA co-labeled with H1 (E) or heterochromatin protein 1 (HP1) (F) in DKO/ΔC and DKO/Z1 nuclei.

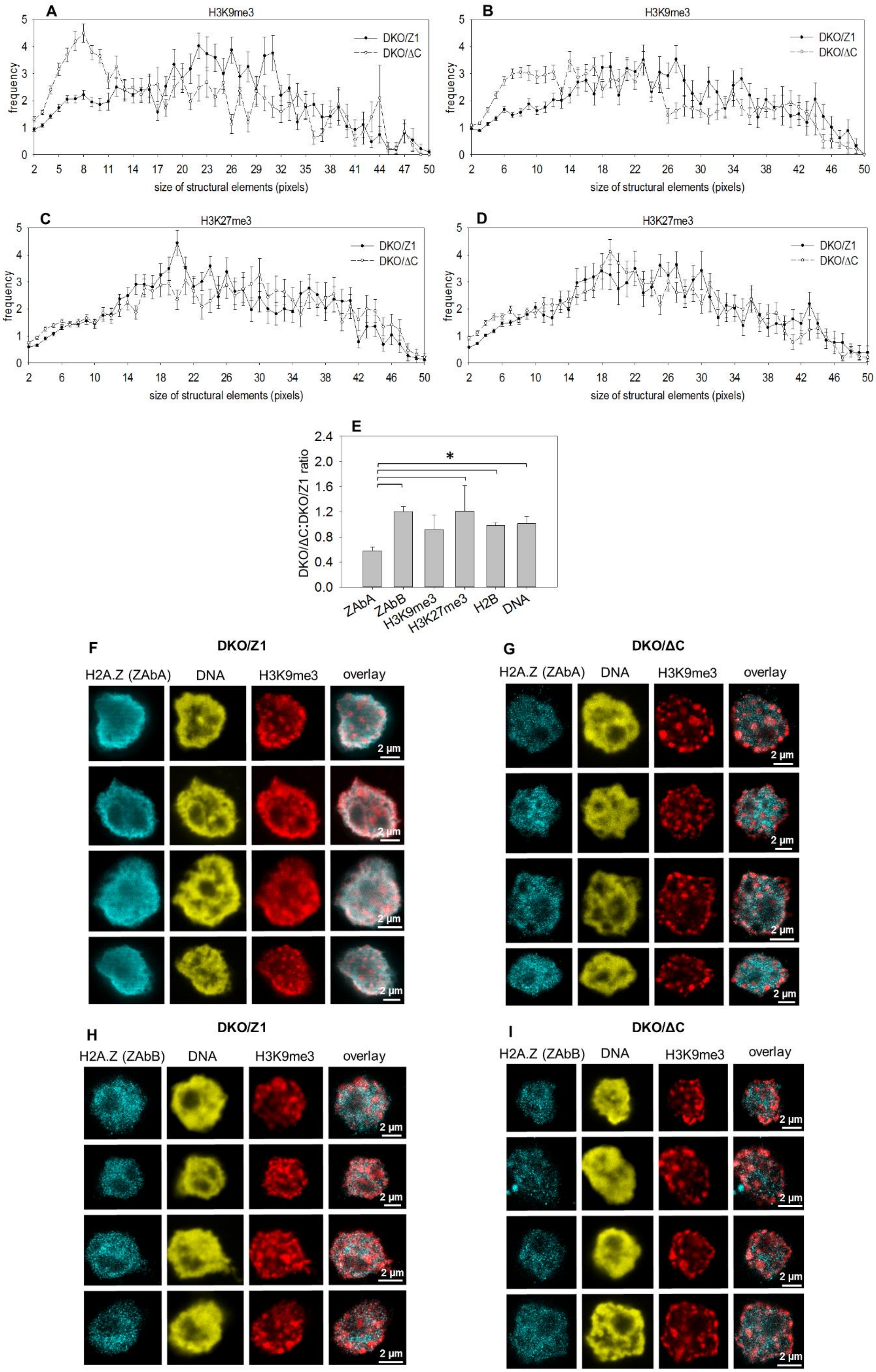

A
H3K9me3
DKO/Z1
DKO/ΔC
frequency
size of structural elements (pixels)
B
H3K9me3
DKO/Z1
DKO/ΔC
frequency
size of structural elements (pixels)
C
H3K27me3
DKO/Z1
DKO/ΔC
frequency
size of structural elements (pixels)
D
H3K27me3
DKO/Z1
DKO/ΔC
frequency
size of structural elements (pixels)
E
*
DKO/ΔC:DKO/Z1 ratio
ZAbA
ZAbB
H3K9me3
H3K27me3
H2B
DNA
F
DKO/Z1
H2A.Z (ZAbA)
DNA
H3K9me3
overlay
2 μm
G
DKO/ΔC
H2A.Z (ZAbA)
DNA
H3K9me3
overlay
2 μm
H
DKO/Z1
H2A.Z (ZAbB)
DNA
H3K9me3
overlay
2 μm
I
DKO/ΔC
H2A.Z (ZAbB)
DNA
H3K9me3
overlay
2 μm

**Suppl. Fig. 11.**

(A-D) Texture analysis of H3K9me3 and H3K27me3 in DT40 cells expressing full length (DKO/Z1) or C-terminally truncated H2A.Z (DKO/ΔC). Curves show the size distribution of structural elements containing H3K9me3 or H3K27me3. Results of two independent experiments. (Experiment 1: A, C; 2: B, D.) Error bars represent SEM. (E) Ratios of proteins (mean fluorescence intensities) detected in DKO/ΔC vs. DKO/Z1 nuclei by ZAbA, ZAbB, or the antibodies specific for H3K9me3, H3K27me3 or H2B, and of PI-stained DNA. Bar chart shows the average and SD values of 4 independent measurements. Statistical analysis was done using one-way ANOVA (* $p \leq 0.001$). (F and G) Superresolution (STED; see Supplementary Materials and Methods) images of H2A.Z containing nucleosomes detected by ZAbA in DKO/Z1 (F) and DKO/ΔC (G) DT40 nuclei, co-labeled with anti-H3K9me3. (H and I) STED images of H2A.Z containing nucleosomes detected by ZAbB in DKO/Z1 (H) and DKO/ΔC (I) DT40 nuclei, co-labeled with anti-H3K9me3.

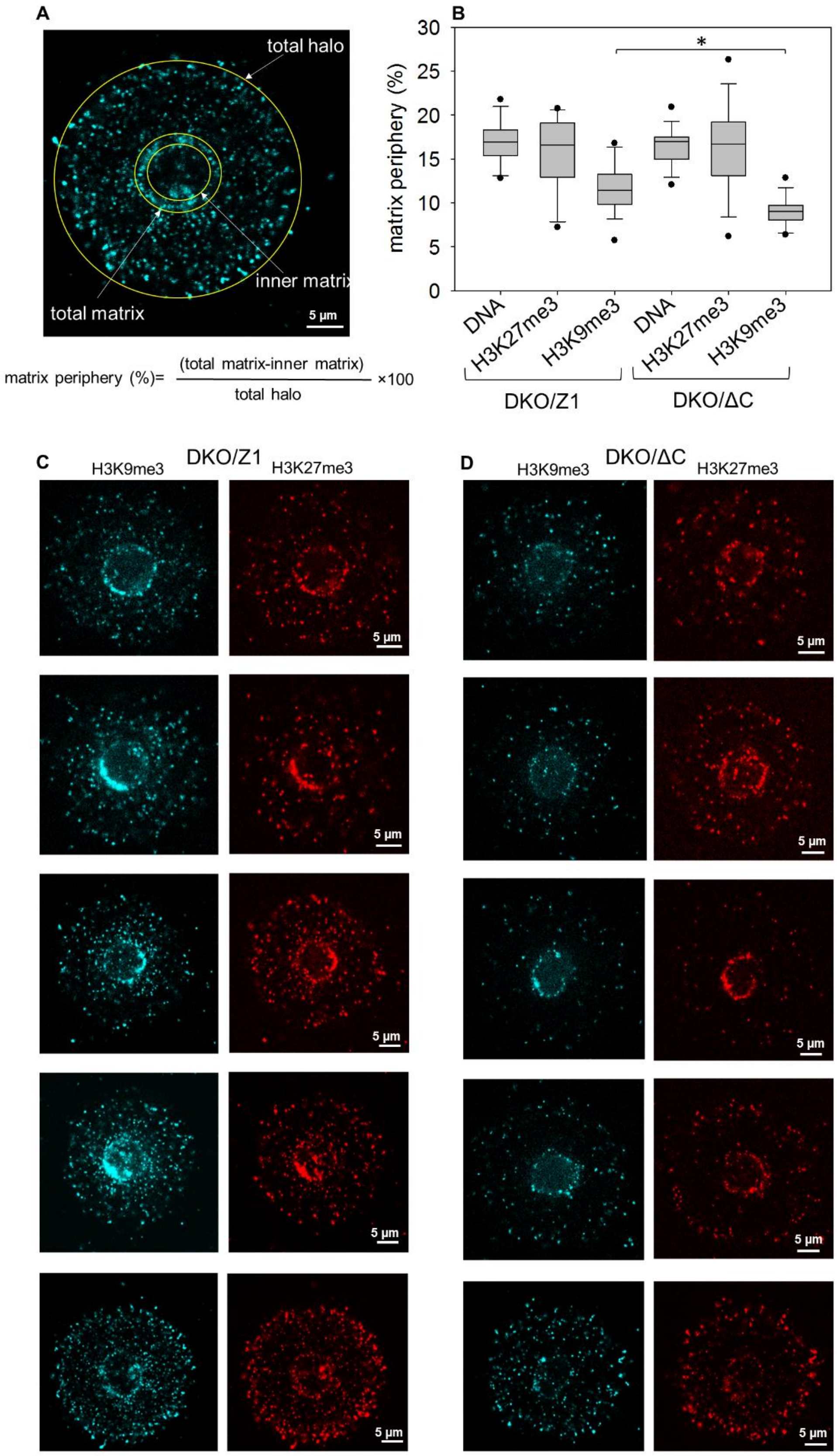
A
total halo
inner matrix
total matrix
5 µm
matrix periphery (%)= (total matrix-inner matrix) / total halo ×100
B
30
25
20
15
10
5
0
matrix periphery (%)
*
DNA
H3K27me3
H3K9me3
DNA
H3K27me3
H3K9me3
DKO/Z1
DKO/ΔC
C
DKO/Z1
H3K9me3
H3K27me3
5 µm
D
DKO/ΔC
H3K9me3
H3K27me3
5 µm

**Suppl. Fig. 12.**

(A-D) Immunofluorescence staining of H3K9me3 and H3K27me3 in halo samples prepared from DKO/ΔC and DKO/Z1 cells. The overall fluorescence signal of the periphery of the nuclei ("designated matrix") was calculated according to panel (A). Box plots in panel (B) show the % of the fluorescence at the nuclear matrix periphery relative to the total fluorescence calculated from 14-17 halos. Representative CLSM images are shown on panel (C) and (D). Statistical analysis was done using one-way ANOVA (* $p \leq 0.001$). Box-and-whisker plot shows the median, 25$^{th}$ and 75$^{th}$ percentiles as vertical boxes with error bars, 5$^{th}$, 95$^{th}$ percentiles and outliers as dots created from the data of ~20 nuclei.

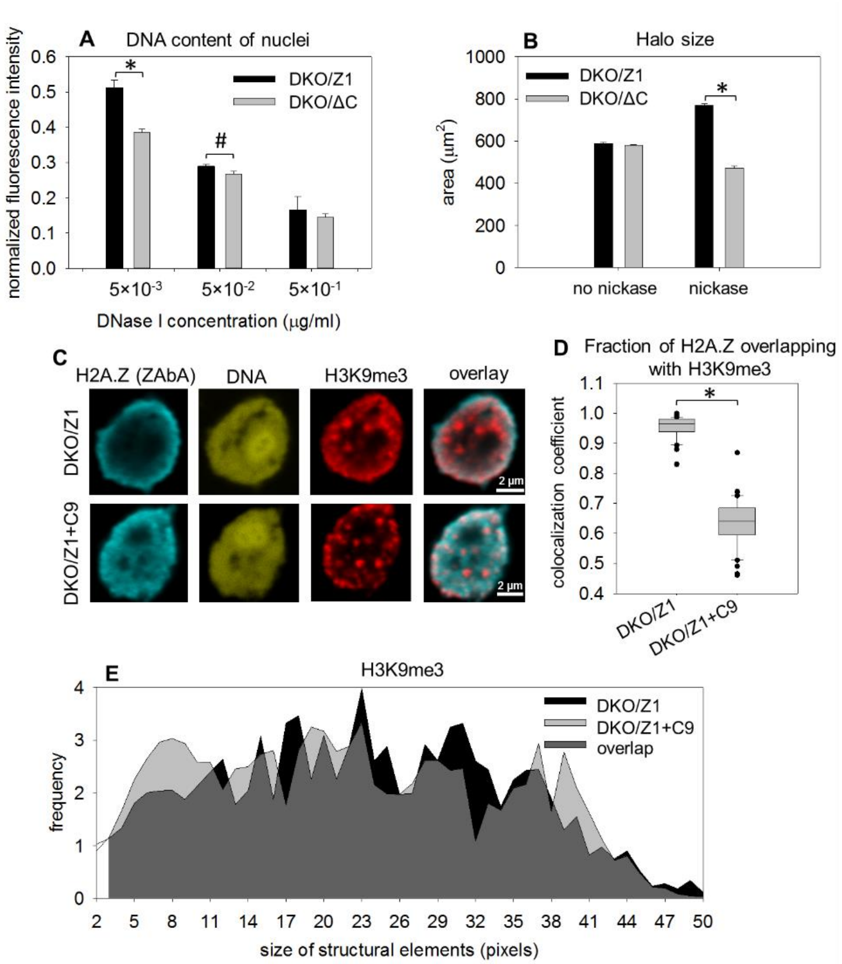

**Suppl. Fig. 13.**

(A) Sensitivity of chromatin to DNase I in DKO/ΔC and DKO/Z1 DT40 nuclei. The DNA content was measured by LSC before and after DNase I endonuclease treatment, at the enzyme concentrations indicated on the figure. (B) Comparison of sensitivities to nickase. Halo size of DKO/ΔC and DKO/Z1 nuclei was measured by LSC before and after 0.5 U/ml nickase treatment, as indicated on the figure. (C) Representative CLSM images showing nuclear localization of H2A.Z recognized by ZAbA and of H3K9me3 co-labeled with H2A.Z, in H2A.Z.1 DKO (DKO/Z1) or C9 treated (DKO/Z1+C9) nuclei. (D) Colocalization analysis of H2A.Z and H3K9me3 in H2A.Z.1 DKO cells, before (DKO/Z1) or after the addition of the C9 peptide

(DKO/Z1+C9). The change of the Manders colocalization coefficient representing the fraction of H2A.Z overlapping with H3K9me3 are shown. Box-and-whisker plot was created from the data of ~40 nuclei. (E) Texture analysis of H2A.Z.1 DKO (DKO/Z1) and C9 treated H2A.Z.1 DKO (DKO/Z1+C9) nuclei showing the size distribution of structural elements containing H3K9me3.

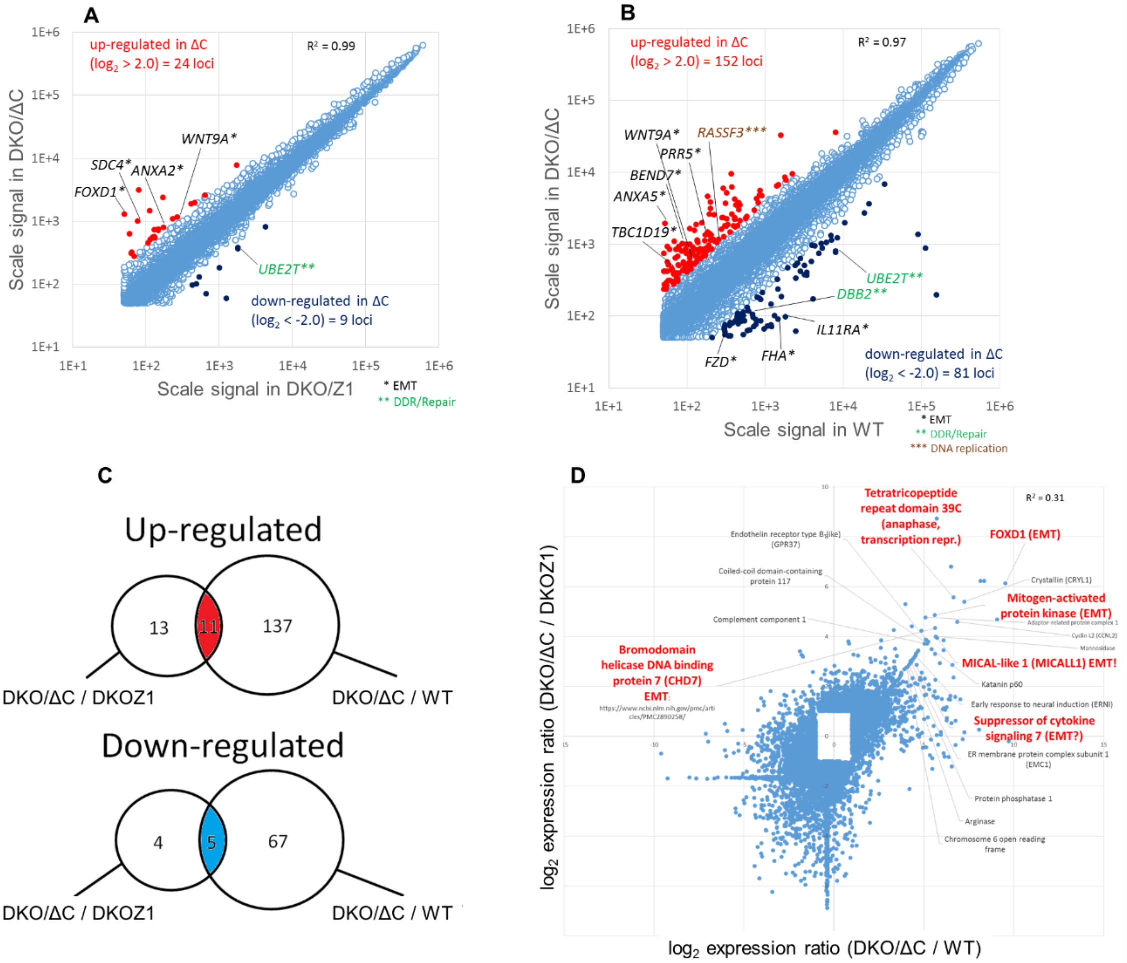

**Suppl. Fig. 14.**

(A-D) Comparison of the gene expression profiles of DKO H2A.Z.1 (designated as DKO/Z1 above), DKO H2A.Z.1ΔC (designated as DKO/ΔC above) and wild-type (WT) DT40 cells. (A) Scatter plot comparing the mRNA expression profiles of DKO/Z1 and DKO/ΔC. Scale signals of gene loci showing moderate or high expression levels (Scale Signal > 50) were used to reduce noise derived from genes with low expression levels. Numbers of up-regulated and down-regulated genes (> 2-fold; red and blue dots, respectively) in DKO/ΔC cells are indicated. The correlation coefficient value ($R^2$) is also shown. (B) Scatter plot comparing the expression profiles of WT and DKO/ΔC. Numbers of up- and down-regulated genes in DKO/ΔC are indicated as described above. *, ** and *** mark proteins contributing to epithelial mesenchymal transition (EMT), DNA repair or DNA replication pathways, respectively. (C) Genes differentially expressed in DKO/ΔC as compared to both DKO/Z1 and WT. Gene loci showing moderate or high expression levels (Scale Signal > 50) in WT were used. In the Venn diagrams depicting the up-regulated and down-regulated genes, the overlaps represent the red and dark-blue dots of panels A and B. (D) Correlation of the DKO/ΔC / DKO/Z1 and DKO/ΔC / WT log2 expression ratios. The correlation coefficient value ($R^2$) is also shown. The group of genes with ratios tightly correlating with each other are indicated. Statistical analysis was done using one-way ANOVA (* p≤0.001, # p≤0.05).

| In ΔC compared to DKO-Z H2A.Z1: | Name | Pathway | Function | ΔC / DKO-Z H2A.Z1 ratio | log2 ratio |
|---|---|---|---|---|---|
| Upregulated: (in ΔC) | | | | | |
| FOXD1 | Forkhead box D1 | FoxO signaling pathway | EMT | 25.76988374 | 4.69 |
| SDC4 | Syndecan 4 | Suppresses EMT | EMT | 13.1187226 | 3.71 |
| ALCAM | Activated leukocyte cell adhesion molecule | Cell adhesion molecule | EMT | 5.11217528 | 2.35 |
| ANXA2 | Annexin A2 | NOD-like receptor signaling pathway | EMT | 4.65439743 | 2.22 |
| WNT9A | Wingless-type MMTV integration site family, member 9A | mTOR signaling pathway | EMT | 4.447939452 | 2.15 |
| Downregulated: (in ΔC) | | | | | |
| UBE2T | ubiquitin-conjugating enzyme E2T | The Fanconi anemia pathway | Repair | 0.206585866 | -2.28 |
| In ΔC compared to WT: | Name | Pathway | Function | ΔC / WT ratio | log2 ratio |
| Upregulated: (in ΔC) | | | | | |
| ANXA5 | Annexin A5 | NOD-like receptor signaling pathway | EMT | 37.59155883 | 5.23 |
| FRZB | Frizzled-related protein | WNT signaling | EMT | 8.380238435 | 3.07 |
| PRR5 | Proline rich 5 | mTOR signaling pathway | EMT | 7.925581779 | 2.99 |
| RASGRP1 | RAS guanyl releasing protein 1 | RAS pathway | EMT | 5.660633439 | 2.50 |
| TBC1D19 | TBC1 domain family member 19 | | EMT | 5.527378464 | 2.47 |
| KLF13 | Kruppel-like factor 13 | | EMT | 5.297023364 | 2.41 |
| WNT9A | Wingless-type MMTV integration site family, member 9A | | EMT | 4.588883338 | 2.20 |
| BEND7 | BEN domain containing 7 | FoxO signaling pathway | EMT | 10.24565168 | 3.36 |
| RASSF3 | Ras association (RalGDS/AF-6) domain family member 3 | | DNA Replication | 4.38985302 | 2.13 |
| Downregulated: (in ΔC) | | | | | |
| FHA | Forkhead-associated (FHA) phosphopeptide binding domain 1 | | EMT | 0.062139246 | -4.00835 |
| RPS6KB1 | Ribosomal protein S6 kinase | WNT signaling | EMT | 0.136636894 | -2.87158 |
| FZD | Frizzled family receptor | | EMT | 0.157378259 | -2.66769 |
| IL6ST | Interleukin 6 signal transducer | | EMT | 0.210157084 | -2.25046 |
| DDB2 | Damage-specific DNA binding protein 2 | | Repair | 0.154487789 | -2.69444 |
| UBE2T | Ubiquitin-conjugating enzyme E2 | Fanconi anemia pathway | Repair | 0.10193932 | -3.29422 |

**Suppl. Table 3.**

List of genes up- or down-regulated in DKO/ΔC relative to DKO/Z1 and WT cells, grouped according to major pathways H2A.Z functioning has been implicated in. Genes with changes of expression level >2 are shown. Comparing DKO/ΔC to DKO/Z1, 74 genes had log2 signal ratio >1.5 (upregulated) and 29 had < –1.5 (downregulated). Comparing DKO/ΔC to WT, 293 genes had log2 signal value >1.5 and 209 had < –1.5, out of 24,530 genes.

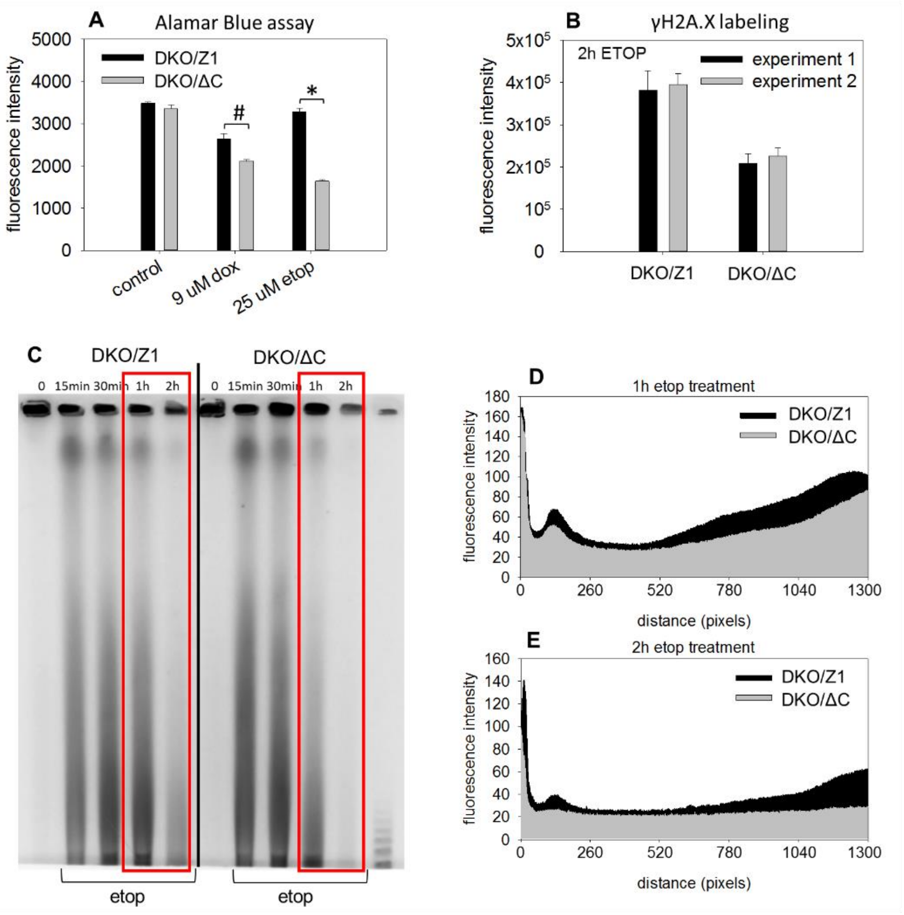

**Suppl. Fig. 15.**

(A) Viability of DKO/ΔC and DKO/Z1 cells after 2h doxorubicin (dox) or etoposide (etop) treatment, as measured by the Alamar Blue assay. (B) γH2A.X expression in DKO/ΔC and DKO/Z1 nuclei detected by immunofluorescence using LSC in two independent experiments. Cells were exposed to 25 µM etoposide for 2h. Bar charts show the mean and SD values of three parallel samples in each experiment. (C) Double-strand break distribution in DKO/ΔC and DKO/Z1 genomic DNA detected by CHEF. Cells were exposed to 25 µM etoposide for 15, 30, 60 and 120 minutes. (D and E) Line-scans corresponding to 60, and 120 minutes treatment in panel C. Statistical analysis was done using one-way ANOVA (* $p \leq 0.001$, # $p \leq 0.05$).

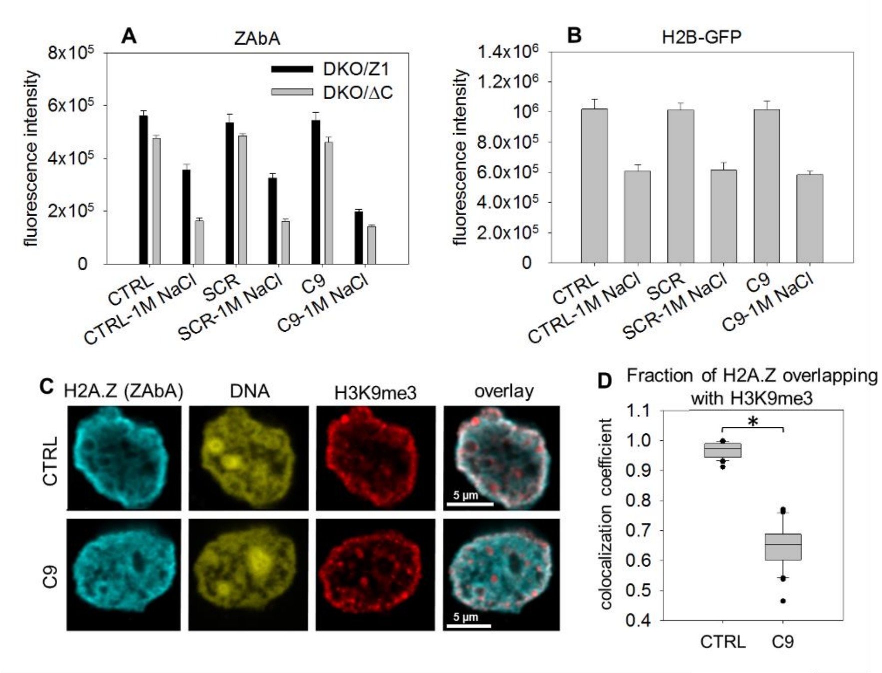

**Suppl. Fig. 16.**

(A) Resistance of H2A.Z nucleosomes to 1 M NaCl in permeabilized DKO/Z1 or DKO/ΔC DT40 nuclei treated with SCR or C9 peptides compared to untreated control (CTRL). H2A.Z was detected by ZAbA. Bar charts show the mean fluorescence intensity, error bars represent the SD of four parallel measurements. (B) Resistance of H2B nucleosomes to 1 M NaCl in permeabilized H2B-GFP expressor HeLa nuclei treated with SCR or C9 peptides compared to untreated control (CTRL). Bar charts show the mean fluorescence intensity, error bars represent the SD of four parallel measurements. (C, D) Colocalization analyses of H2A.Z and H3K9me3 before and after incubation of agarose-embedded Hela nuclei with the C9 peptide used at a concentration of 30 µM. H2A.Z was detected by ZAbA and Manders colocalization coefficients showing the fraction of H2A.Z overlapping with H3K9me3. Statistical analysis was done using one-way ANOVA (* $p \leq 0.001$, # $p \leq 0.05$). Box-and-whisker plot shows the median, 25$^{th}$ and 75$^{th}$ percentiles as vertical boxes with error bars, 5$^{th}$, 95$^{th}$ percentiles and outliers as dots created from the data of ~36 nuclei.

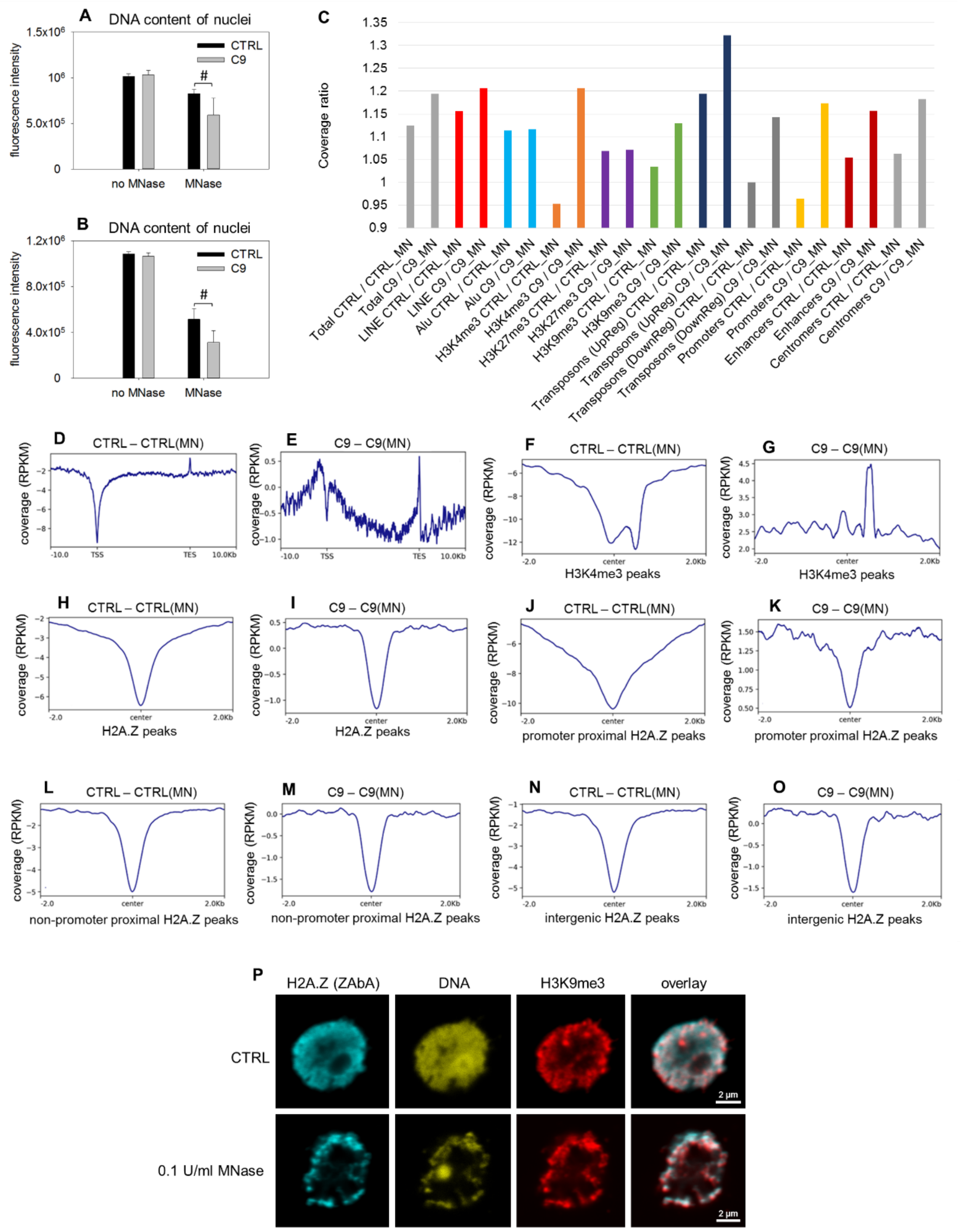

**Suppl. Fig. 17.**

(A and B) Two independent replicates of the experiment shown in Figure 3F. Bar chart shows the average and SD values of 4 parallel measurements. Statistical analysis was done using one-way ANOVA (# $p \leq 0.05$). (C) Effect of C9 on MNase sensitivity. Coverage ratios comparing the MNase treated (MN) and non-treated samples for both C9 treated (C9) and untreated (CTRL) samples were calculated for the whole genome (total – indicated with grey), LINE (red) and Alu (light blue) repetitive elements, H3K4me3 (yellow), H3K27me3 (purple) and H3K9me3 (green) peaks, transposons upregulated in our RNA-seq (UpReg – dark blue), downregulated in our RNA-seq (DownReg - dark grey), promoter (yellow) and enhancer (dark red) elements and centromers (grey). Peaksets used were downloaded from ENCODE database, data accession numbers are indicated in Suppl. Methods. Centromer and Alu/LINE elements were downloaded using the UCSC Main Table Browser and RepeatMasker. (D and E) Metaplots showing the distribution of CTRL – CTRL(MN) and C9 – C9(MN) read coverage differences (D and E, respectively) in the 10 kb vicinity of transcription start sites. (F-O) Anchor plots of CTRL – CTRL(MN) (F, H, J, L, N) and C9 – C9(MN) (G, I, K, M, O) read coverage differences around H3K4me3 (F and G) and H2A.Z (H and I) peaks. Anchor plots of CTRL – CTRL(MN) and C9 – C9(MN) coverage differences around promoter proximal (+/- 2 kb) (panels J and K), non-promoter proximal (panels L and M) and intergenic H2A.Z peaks (panels N and O). (P) CLSM images showing the localization of H2A.Z and H3K9me3 containing nucleosomes labeled with ZAbA and goat anti-rabbit secondary antibody in DKO/Z1 DT40 nuclei, after treatment with 0.1 U/ml MNase or without treatment (CTRL).

| Accession | Description | MW [kDa] | Abundance Ratio |
|---|---|---|---|
| P27695 | APEX1 | 35.5 | 6.7* |
| O60264 | SMARCA5 # | 121.8 | 17.5** |
| Q99459 | CDC5L | 92.2 | 16.8** |
| P0C0S5 | H2A.Z.1 | 13.5 | 43.7** |
| P33992 | MCM5 # | 82.2 | 50.2** |
| Q9UMS4 | PRPF19 # | 55.1 | 8.9* |
| P55060 | CSE1L | 110.3 | 79.0** |
| P33991 | MCM4 # | 96.5 | 20.9** |
| P33993 | MCM7 # | 81.3 | 7.2* |
| P60900 | PSMA6 | 27.4 | 12.9** |
| P62807 | H2B | 13.9 | 7.7* |
| P13639 | EEF2 | 95.3 | 22.2** |
| Q01081 | U2AF1 # | 27.9 | 5.4 |

**Suppl. Table 4.** Effect of C9 treatment on the accessibility of H2A.Z-containing chromatin to ZAbA/MNase.

Nuclear proteins of a ZAbA/anti-rabbit antibody/CUT&RUN experiment enriched >5x in the samples after C9 pretreatment as compared to the peptide-untreated control. Proteins marked with # are expressed in a H2A.Z-dependent manner based on correlation of their expression level with that of H2A.Z across the NCI60 panel of human cancer cell lines (cellminercdb; https://discover.nci.nih.gov/rsconnect/cellminercdb/) (** significant at $p<0.01$; * significant at $p<0.05$).

Accession: protein identifier in the Swissprot database; abundance ratio: ratio of protein abundances detected in the C9 treated sample compared to the untreated control. Protein abundances are calculated as the summed abundance of peak areas of the peptides identifying the respective protein.

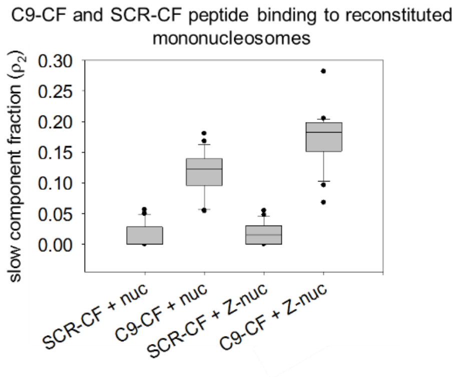

**Suppl. Fig. 18.** FCS analysis of C9-CF and SCR-CF peptide binding to reconstituted mononucleosomes.

Slow fraction of C9-CF and SCR-CF peptides after addition of H2A containing nucleosomes (nuc) or H2A.Z containing nucleosomes (z-nuc). The peptide was incubated with nucleosomes at 5 mM NaCl concentration. The slow fraction $\rho_2$ was calculated in ACF fits as described in the Materials and Methods and presented as a box-and-whisker plot.

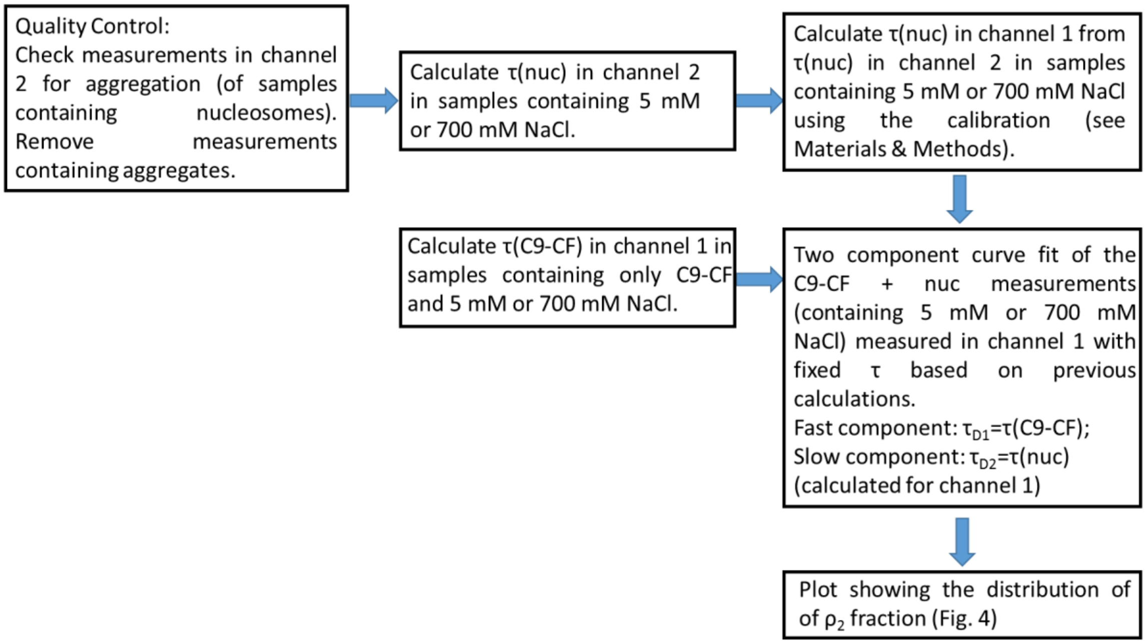

**Suppl. Fig. 19.**

Pipeline of FCS data analysis in Fig. 4.

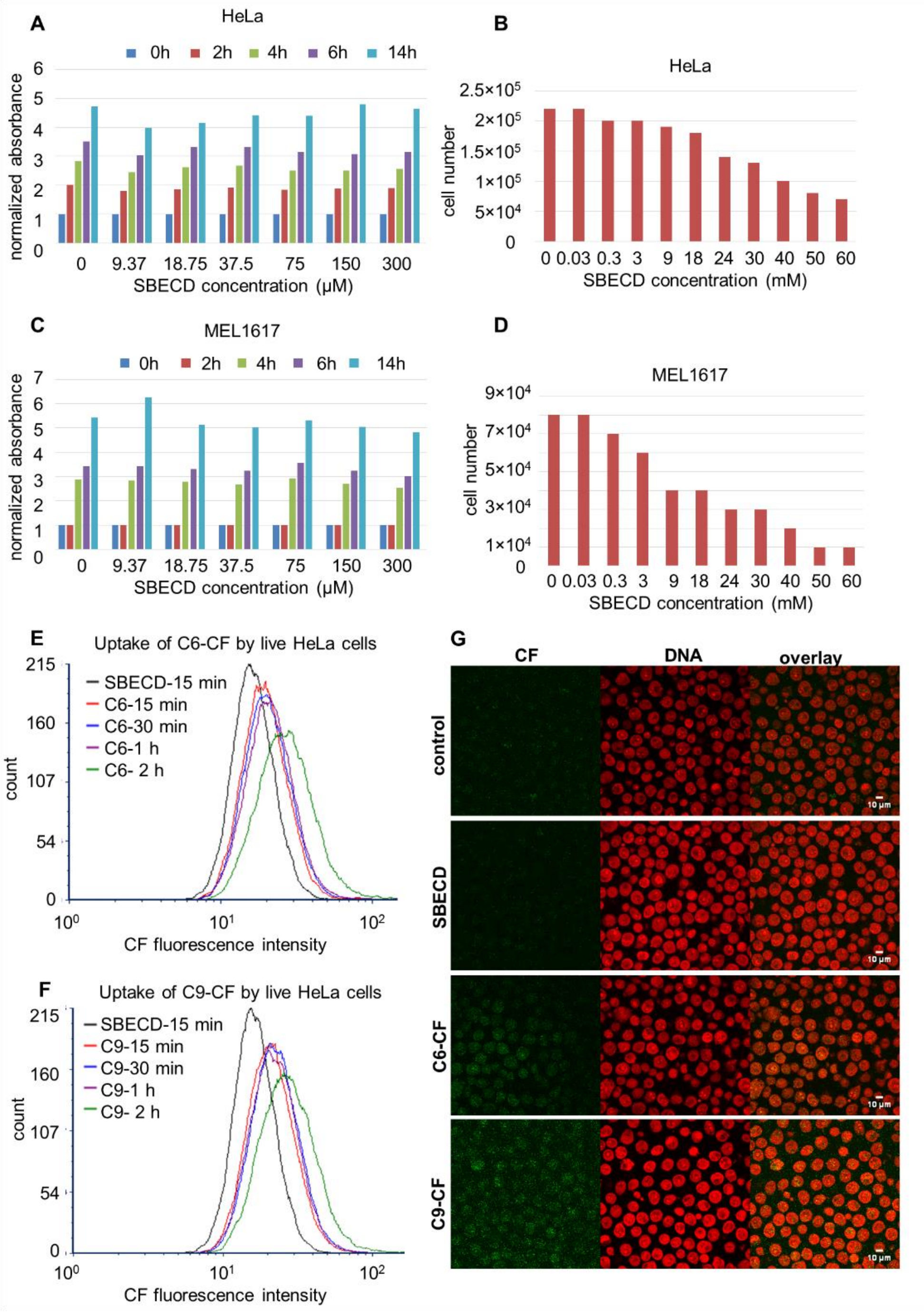

**Suppl. Fig. 20.**

(A) Titration of SBECD concentration in the micromolar range in the case of HeLa cells measuring cell viability by the Alamar Blue assay. (B) Titration of SBECD concentration in the case of HeLa cells in the millimolar range quantifying cell number by cell counting. (C and D) SBECD titration performed using melanoma cells (MEL1617), as in panel (A) and (B), respectively. (E and F) Kinetics of carboxyfluorescein-C6 (E) or carboxyfluorescein-C9 (F) uptake by live HeLa cells analyzed by flow cytometry. Complex formation was performed using 300 μM SBECD and 30 μM peptide. (G) Accumulation of the fluorescent peptides in the nucleus of HeLa cells visualized by CLSM, after fixation of the cells in 1% formaldehyde.

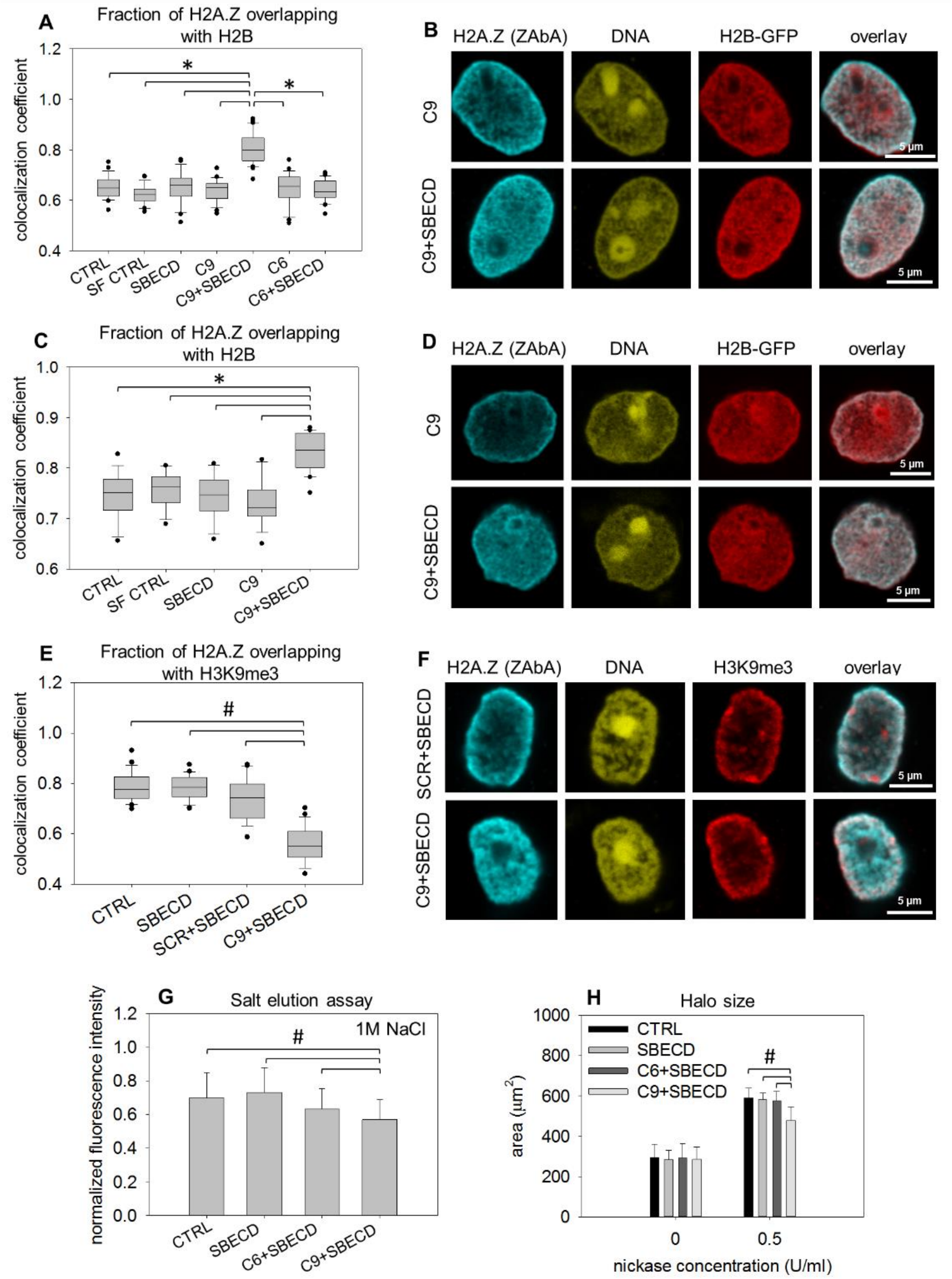

**Suppl. Fig. 21.**

(A-D) Colocalization analysis of H2A.Z and H2B after incubation of live Hela cells with serum free medium (SF CTRL), cyclodextrin (SBECD), C9 (C9), C9 in complex with SBECD

(C9+SBECD) used at a concentration of 30 µM and 300 µM, respectively, C6 (C6) or C6 in complex with SBECD (C6+SBECD) used at the same peptide/cyclodextrin ratio. Results of two independent experiments are shown in panels A, B and in panels C, D. H2A.Z was detected by ZAbA. Manders colocalization coefficients showing the fraction of H2B overlapping H2A.Z were calculated (A, C). CLSM images showing the nuclear localization of H2A.Z-containing nucleosomes detected by ZAbA and H2B-GFP in HeLa cells treated with C9 only or with the peptide in complex with SBECD (B, D). (E) Manders colocalization coefficients representing the fraction of H2A.Z overlapping with H3K9me3 calculated for the nuclei shown in panel F are shown. (F) Representative confocal images of H2A.Z-H3K9me3 double-stained nuclei following introduction of C9 into live HeLa cells. Cells were treated with the C9 or C9 SCR using SBECD (C9+SBECD, C9 SCR+SBECD), or with cyclodextrin in the absence of peptides (SBECD). (G) Resistance of H2A.Z nucleosomes to 1 M NaCl in permeabilized HeLa nuclei prepared from untreated, cyclodextrin-treated, C6+SBECD- or C9+SBECD-treated HeLa cells. Bar charts show the mean fluorescence intensity, error bars represent the SD of two parallel measurements united. (H) Comparison of chromatin sensitivity to nickase. Halo size of untreated (CTRL), SBECD-, C6+SBECD- and C9+SBECD-treated nuclei was measured by LSC before and after nickase treatment. Bar charts show the mean fluorescence intensity, error bars represent the SD of 4 parallel measurements. Statistical analysis was done using one-way ANOVA (* $p \leq 0.001$, # $p \leq 0.05$). Box-and-whisker plot shows the median, 25$^{th}$ and 75$^{th}$ percentiles as vertical boxes with error bars, 5$^{th}$, 95$^{th}$ percentiles and outliers as dots created from the data of ~37 (A), ~20 (C) or ~28 (E) nuclei.

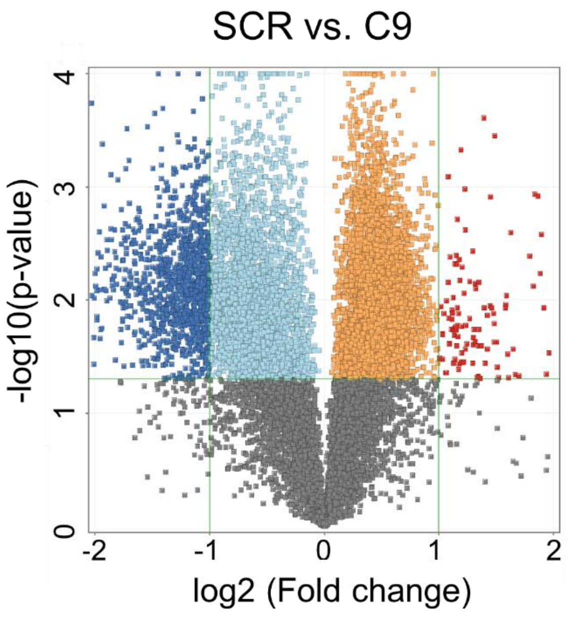

**Suppl. Fig. 22.**
Volcano plot showing differentially expressed genes defined as log2 fold change < or > 1.0 and $p < 0.05$ (dark blue and red dots) comparing SCR vs. C9 treated samples of MEL1617 cells.

SUPPLEMENTARY MATERIALS AND METHODS

### Microarray analysis

Dye incorporation and cRNA yield were checked with the NanoDrop ND-2000 Spectrophotometer. 0.6 micro-g of Cy3-labeled cRNA was fragmented at 60°C for 30 minutes in a reaction volume of 25 micro-L containing 1x Agilent fragmentation buffer and 2x Agilent blocking agent following the manufacturer's instructions. On completion of the fragmentation reaction, 25 micro-L of 2x Agilent hybridization buffer was added to the fragmentation mixture and hybridized to G. gallus (Chicken) Oligo Microarray v2 (Agilent Technologies) for 17 hours at 65°C in a rotating Agilent hybridization oven. After hybridization, microarrays were washed 1 minute at room temperature with GE Wash Buffer 1 (Agilent Technologies) and 1 minute with 37°C GE Wash buffer 2 (Agilent Technologies).Slides were scanned immediately after washing on the Agilent SureScan Microarray Scanner (G2600D) using one color scan setting for 8x60k array slides (Scan Area 61x21.6 mm, Scan resolution 3 micro-m, Dye channel is set to Green PMT is set to 100%).
The scanned images were analyzed with Feature Extraction Software (Agilent Technologies) using default parameters to obtain background subtracted and spatially detrended Processed Signal intensities. Processed signal intensities were normalized by the global scaling method. A trimmed mean probe intensity was determined by removing 2% of the lower and the higher end of the probe intensities in order to calculate the scaling factor. Normalized signal intensities were then calculated from the target intensity on each array using the scaling factor, so that the trimmed mean target intensity of each array was arbitrarily set to 2500.

### RNA-Seq analysis

RNA-Seq libraries were prepared from total RNA using Ultra II RNA Sample Prep kit (New England BioLabs) according to the manufacturer's protocol. Briefly, poly-A RNA was captured on oligo-dT conjugated magnetic beads then the mRNA was eluted and fragmented at 94 $^{o}$C. First strand cDNA was generated by random priming reverse transcription, followed by second strand synthesis yielding double stranded cDNA. After end repair and A-tailing, the adapter ligated fragments were amplified in enrichment PCR and finally sequencing libraries were generated. Sequencing runs were executed on Illumina NextSeq 500 instrument using single-end 75 cycles sequencing.

### Bioinformatic analysis of the fastq data

Raw sequencing data (fastq) were aligned to the human reference genome GRCh38 using the HISAT2 (daehwankimlab.github.io/hisat2) algorithm and BAM files were generated. Downstream analysis was performed using StrandNGS (www.strand-ngs.com). BAM files were imported into the DESeq algorithm for normalization. Moderated T-test with Benjamini-Hochberg FDR was used to determine differentially expressed genes between the compared conditions. Dysregulated genes were converted into BED format with UCSC Main Table Browser (genome.ucsc.edu/cgi-bin/hgTables). CytoScape v3.4 software with ClueGo v2.3.5. application was used for identifying over- represented Gene ontology (GO) terms of the list of differentially expressed genes. Two-sided hypergeometric test with Benjamini-Hochberg correction was performed and GO Biological process database was used.

**Bioinformatic analysis of the Whole Genome Sequencing data**

Quality control was performed on the raw sequenced reads using the FastQC tool (https://www.bioinformatics.babraham.ac.uk/projects/fastqc/). The paired end sequencing reads were mapped to the reference genome (hg19) with BWA-MEM [7]. Samtools [8] was used to filter the mapped reads (BAM files), the minimum MAPQ quality score was 20. BamCompare (deeptools package [9] was used to divide or subtract two BAM files from each other, computeMatrix and plotProfile tools were used to create anchor- and metaplots). The LINE1 and Alu repetitive elements were downloaded using the UCSC Main Table Browser (https://genome.ucsc.edu/cgi-bin/hgTables). PlotCoverage (deeptools) was used to calculate median per base coverage for either the whole genome or the LINE1 and Alu repeats.

The following peaksets or coverage data for HeLa cells were downloaded from the ENCODE database [10]: CTCF ChIP-seq peaks and coverage, ENCFF768OPK and ENCFF799KLZ, respectively; H2A.Z peaks: ENCFF415ESA; H3K9me3 peaks: ENCFF712ATO; H3K4me3 peaks: ENCFF447CLK. Lamin A and Lamin B1 ChIP-seq peaks were downloaded from the NCBI GEO database (accession numbers: GSM1376181 and GSM1376181, respectively).

**Bioinformatic analyses of C9-induced changes in transposable element expression**

Based on https://pubmed.ncbi.nlm.nih.gov/29508296/, the transposable element (TE) annotation file for hg38 was downloaded through http://hammelllab.labsites.cshl.edu/software, and was used to analyze the expression of TEs from the bam files using the featurecounts and DESeq2 programs on the Galaxy.eu server (https:usegalaxy.eu).

**LC-MS/MS analysis of CUT&RUN samples**

The samples were digested with trypsin according to the Strap micro high recovery protocol (https://files.protifi.com/protocols/s-trap-micro-high-recovery-quick-card-2.pdf, accessed on 23 February 2023). Briefly, samples were reduced using TCEP (Tris(2-carboxyethyl)phosphine, 6mM) for 15 min at 37°C, followed by alkylation with MMTS (S-methyl methanethiosulfonate, 25 mM) for 15 min at room temperature before digesting with MS-grade trypsin (Thermo Scientific, Waltham, MA, USA) for 2h at 47 °C. 40% of the peptide resulting peptide mixtures were analyzed with online LC-MS/MS using a nanoAcquity nanoLC (Waters) - Orbitrap Elite mass spectrometer (Thermo Scientific) system. Peptides were separated using a 40-min water-acetonitrile gradient at 250 nl/min followed by MS/MS analysis of the top 20 most abundant multiply charged precursor ions (Orbitrap analyzer resolution: 60,000, AGC target: 1.0e6, collision-induced dissociation in the linear ion trap with 35% normalized collision energy, AGC target: 5.0e4, dynamic exclusion 60 s). Raw data were processsed using the Proteome Discoverer software (v2.4.1.15). Proteins were identified using the Byonic search engine against the Swissprot human database (release: 2022.03, 20306 entries). Methylthio derivatization of Cys was set as fixed modification, oxidation of Met, pyroglutamic acid formation from peptide N-terminal Gln, and acetylation and/or Met cleavage of protein N-terminus were set as variable modifications allowing maximum 2 variable modifications per peptide. Trypsin was specified as enzyme allowing maximum two missed cleavage sites per peptide. A minimum score of 200 was specified for peptide-spectrum matches (PSM) and protein-level FDR was set to maximum 1%. Protein abundances were normalized to the total peptide amount and semiquantitative comparison was performed using label free quantification.

### Stimulated emission depletion (STED) microscopy

STED superresolution images were taken with a STEDYCON (Abberior GmbH, Göttingen, Germany) confocal/STED extension mounted on an Olympus BX43 upright microscope equipped with an Olympus 100× UPLXAPO oil immersion objective NA 1.45. Abberior Star Orange and Star Red dyes were excited at 561 and 640 nm, then illuminated with a 775 nm STED laser to deplete the excited state. Fluorescence was detected by PMTs between 575-625 and 650-700 nm, the optical resolution was 70 and 50 nm in the two channels. 8 µm × 8 µm areas (307 × 307 pixels) were scanned with a pixel size of 25 nm, pixel dwell time of 3 µs (confocal) and 10 µs (STED), pinhole size of 32 µm. The nucleus was counterstained with DAPI and imaged with confocal resolution (exc: 405 nm, em: 420-475 nm).

### Fitting of FCS autocorrelation curves

Autocorrelation curves of the C9-CF peptide were fitted to a model assuming a single 3D free diffusion component with triplet state correction:

$$G(\tau) = \frac{1-T+Te^{-\frac{\tau}{\tau_{trip}}}}{N(1-T)} \frac{1}{1+\frac{\tau}{\tau_D}} \frac{1}{\sqrt{1+\frac{\tau}{S^2\tau_D}}} \quad \text{eq.1}$$

$N$ is the average number of fluorescent molecules in the detection volume, $T$ is the fraction of molecules in the triplet state, $\tau_{trip}$ is the triplet correlation time. The rate of diffusion is characterized by the diffusion time, $\tau_D$, which is the average time that a molecule spends in the illuminated volume. $S$ corresponds to the aspect ratio of the ellipsoid-shaped confocal volume, defined as the ratio of its axial to radial dimensions. This parameter for both channels was estimated by fitting the autocorrelation curves of 100 nM Alexa 488 or 100 nM Alexa 647 dyes (dissolved in 10 mM Tris-EDTA buffer, pH 7.4).

Autocorrelation curves of the C9-CF peptide mixed with DNA or nucleosomes (nuc) were fitted with a model function including triplet state transition, and assuming two diffusing species: a fast component describing the diffusion of unbound peptide, and a slow component corresponding to the diffusion of peptide bound to nucleosomes or DNA:

$$G(\tau) = \frac{1-T+Te^{-\frac{\tau}{\tau_{trip}}}}{N(1-T)} \left(\rho_1 \frac{1}{1+\frac{\tau}{\tau_{D1}}} \frac{1}{\sqrt{1+\frac{\tau}{S^2\tau_{D1}}}} + \rho_2 \frac{1}{1+\frac{\tau}{\tau_{D2}}} \frac{1}{\sqrt{1+\frac{\tau}{S^2\tau_{D2}}}}\right) \qquad \text{eq.2}$$

$\tau_{D1}$ and $\tau_{D2}$ are the diffusion times of the fast and slow components, and $\rho_1$ and $\rho_2$ are their fractions. For determining the fraction of DNA- or nucleosome-bound C9-CF, the diffusion times of the fast and slow components were fixed. The diffusion time of the fast component ($\tau_{D1}$) was determined by fitting the ACF from a sample containing only C9-CF according to equation 1. Although the diffusion times of the DNA- or nucleosome-bound C9-CF may rightfully be assumed to be equal to that of the diffusion times of Cy5-labeled nucleosome/DNA, the diffusion times measured in green (CF) and red (Cy5) channels may differ since the geometries of the observation volumes are different in these channels. Therefore, for estimating the diffusion time of DNA/nucleosome-bound C9-CF in the CF channel, the following equation was used:

$$\tau_{D2} = \omega_{green}^2/\omega_{red}^2 \ \tau_{nuc} \qquad \text{eq.3}$$

$\tau_{D2}$ is the calculated (would-be) diffusion time of the nucleosome/DNA-bound C9-CF in the CF channel; $\tau_{nuc}$ is the measured diffusion time of the Cy5-labeled nucleosome/DNA in the Cy5 channel. $\omega_{green}$ and $\omega_{red}$ are the lateral $e^{-2}$ radii of the detection volumes in the CF and Cy5 channels, which were determined by measuring the diffusion times of 100 nM fluorescein and 100 nM Cy5 dyes (dissolved in 10 mM Tris-EDTA buffer, pH 7.4, $D_{Fluor}$ = 425 µm$^2$/s and $D_{Cy5}$ = 360 µm$^2$/s, at T = 22.5°C) in their respective channels and substituting them into the following equation:

$$\omega_{xy}^2 = 4\tau_D D \qquad \text{eq.4}$$

### SUPPLEMENTARY DISCUSSION

H2A.Z may affect HP1's switch to a crosslinking-competent conformation [11], what could influence the formation and tethering of the phase-separated heterochromatin. Indeed, H2A.Z collaborates with HP1 in an H1-dependent manner to maintain the H3K9me3 marked constitutive heterochromatin in a lamina-tethered, phase-separated state [12, 13, 14, 15]. According to ref. [13], HP1α interacts with the linker DNA unoccluded by linker histones, and also with the nucleosome core, and this latter mode of binding is augmented by both H2A.Z and H3K9me3. HP1 and H2A.Z may also directly interact with each other [16]. A scenario including H1 in the picture could also be considered based on the observation that the C-terminal tail of H2A.Z disfavors H1 binding [13, 17, 18], although H1 appears to accompany H2A.Z in the DKO H2A.Z.1 as well as in DKO H2A.ZΔC cells (Suppl. Fig. 10E) and that observation was also challenged based on data obtained with alternative methods [19].

The special significance of the C-terminus in DDR was also supported independently by mutational analysis [20]. The chromatin remodeling steps following double-strand break induction involve sequential H2A.Z deposition and removal [21]; in view of the waned DDR of the ΔC cells in response to DNA damaging agents, its C-terminus may play an important role in the dynamics or in the functioning of the histone variant. Proteins belonging to the Fanconi anemia pathway involved in topoisomerase II poison-elicited DDR [22] were expressed in a H2A.Z C-terminal tail-dependent manner (e.g. UBE2T; see in Suppl. Table 3), potentially also explaining the relative vulnerability of H2A.ZΔC cells to such drugs. Involvement of H2A.Z in DNA repair may also be related to its role in the retention of transiently stalled replication forks, based on yeast analogy [23].

C9-induced changes of transposable element expression: Out of the more than 4.7 million TEs listed in the annotation file, only about 7000 TEs had detectable expression (raw counts > 20), and 251 had significantly different expression (adj. p value $\leq 0.05$ and abs(log2FC) $\geq 1.2$). The 96 upregulated TEs included mostly LINEs, plus 16 DNA transposons, 12 LTRs, and 6 SINEs. The 155 downregulated TEs were skewed towards SINEs, in addition to 27 LINEs, 17 DNA transposons, 14 LTRs, 2 satellites and 1 retroposon. These data suggest that the alteration in chromatin structure (Suppl. Fig. 17C) is accompanied by changes in transcriptional activity also in these heterochromatic regions. Limitations of our analysis include that only polyadenylated TE transcripts could be detected and quantified reliably and that only a very small percentage of all TEs was detected. (Remark: Although the cellular sources of the MNase and RNA-seq experiment were different, this is perhaps of a lesser concern in the case of the repetitive elements.)
